# Clipper: p-value-free FDR control on high-throughput data from two conditions

**DOI:** 10.1101/2020.11.19.390773

**Authors:** Xinzhou Ge, Yiling Elaine Chen, Dongyuan Song, MeiLu McDermott, Kyla Woyshner, Antigoni Manousopoulou, Ning Wang, Wei Li, Leo D. Wang, Jingyi Jessica Li

## Abstract

High-throughput biological data analysis commonly involves identifying features such as genes, genomic regions, and proteins, whose values differ between two conditions, from numerous features measured simultaneously. The most widely-used criterion to ensure the analysis reliability is the false discovery rate (FDR), which is primarily controlled based on p-values. However, obtaining valid p-values relies on either reasonable assumptions of data distribution or large numbers of replicates under both conditions. Clipper is a general statistical framework for FDR control without relying on p-values or specific data distributions. Clipper outperforms existing methods for a broad range of applications in high-throughput data analysis.

## Introduction

High-throughput technologies are widely used to measure system-wide biological features, such as genes, genomic regions, and proteins (“high-throughput” means the number of features is large, at least in thousands). The most common goal of analyzing high-throughput data is to contrast two conditions so as to reliably screen “interesting features,” where “interesting” means “enriched” or “differential.” “Enriched features” are defined to have higher expected measurements (without measurement errors) under the experimental (i.e., treatment) condition than the background (i.e., negative control) condition. The detection of enriched features is called “enrichment analysis.” For example, typical enrichment analyses include calling protein-binding sites in a genome from chromatin immunoprecipitation sequencing (ChIP-seq) data [1, 2] and identifying peptides from mass spectrometry (MS) data [3]. In contrast, “differential features” are defined to have different expected measurements between two conditions, and their detection is called “differential analysis.” For example, popular differential analyses include the identification of differentially expressed genes (DEGs) from genome-wide gene expression data (e.g., microarray and RNA sequencing (RNA-seq) data [4–10]) and differentially interacting chromatin regions (DIRs) from Hi-C data [11–13] (Fig. 1a). In most scientific research, the interesting features only constitute a small proportion of all features, and the remaining majority is referred to as “uninteresting features.”

**Figure 1.**
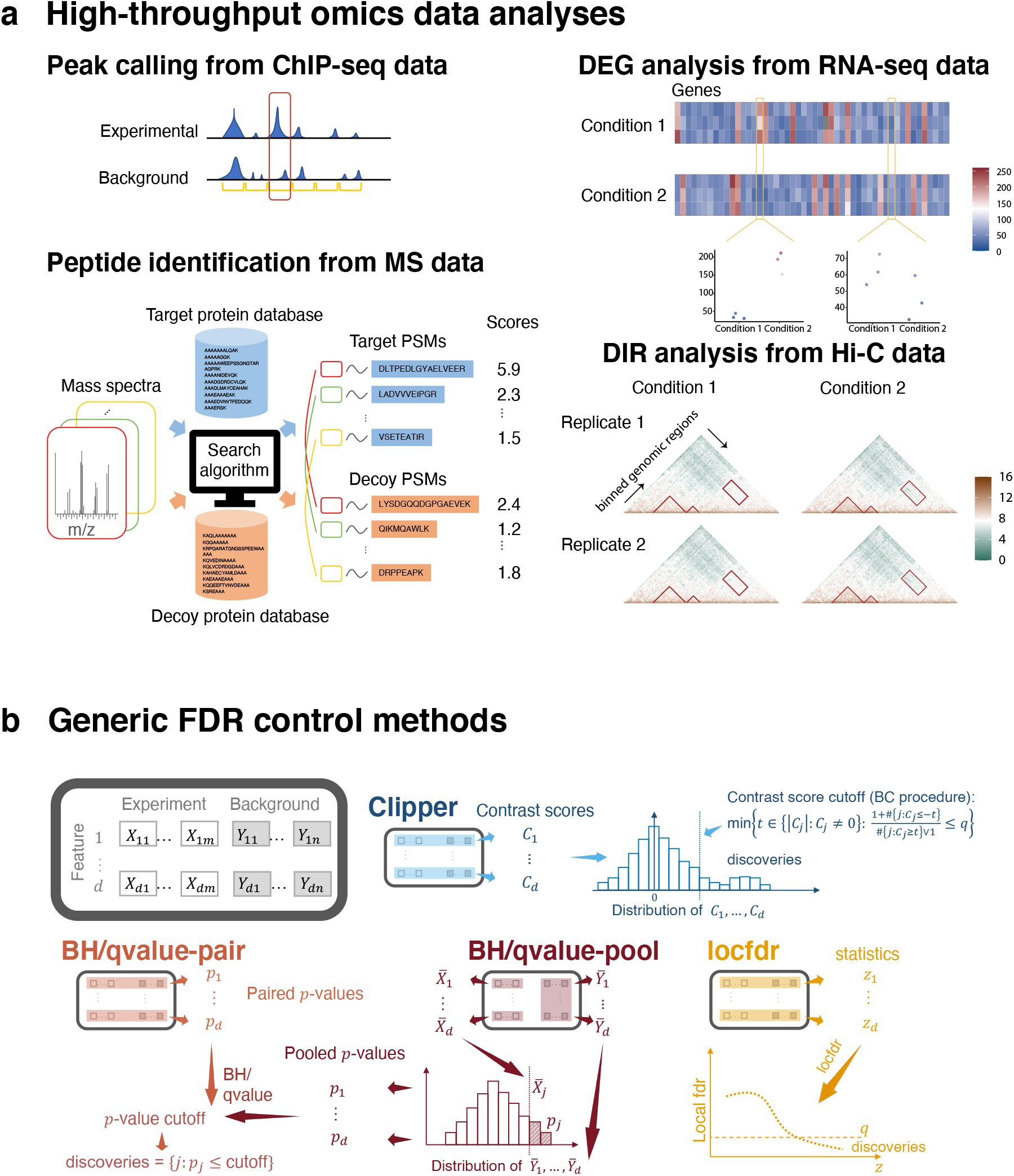
High-throughput omics data analyses and generic FDR control methods. **(a)** Illustration of four common high-throughput omics data analyses: peak calling from ChIP-seq data, peptide identification from MS data, DEG analysis from RNA-seq data, and DIR analysis from Hi-C data. In these four analyses, the corresponding features are genomic regions (yellow intervals), peptide-spectrum matches (PSMs; a pair of a mass spectrum and a peptide sequence), genes (columns in the heatmaps), and chromatin interacting regions (entries in the heatmaps). **(b)** Illustration of Clipper and five generic FDR control methods: BH-pair (and qvalue-pair), BH-pool (and qvalue-pool), and locfdr. The input data are *d* features with *m* and *n* repeated measurements under the experimental and background conditions, respectively. Clipper computes a contrast score for each feature based on the feature’s *m* and *n* measurements, decides a contrast-score cutoff, and calls the features with contrast scores above the cutoff as discoveries. (This illustration is Clipper for enrichment analysis with *m* = *n*.) BH-pair or qvalue-pair computes a p-value for each feature based on the feature’s *m* and *n* measurements, sets a p-value cutoff, and calls the features with p-values below the cutoff as discoveries. BH-pool or qvalue-pool constructs a null distribution from the *d* features’ average (across the *n* replicates) measurements under the background condition, calculates a p-value for each feature based on the null distribution and the feature’s average (across the *m* replicates) measurements under the experimental condition, sets a p-value cutoff, and calls the features with p-values below the cutoff as discoveries. The locfdr method computes a summary statistic for each feature based on the feature’s *m* and *n* measurements, estimates the empirical null distribution and the empirical distribution of the statistic across features, computes a local fdr for each feature, sets a local fdr cutoff, and calls the features with local fdr below the cutoff as discoveries.

The identified features, also called the “discoveries” from enrichment or differential analysis, are subject to further investigation and validation. Hence, to reduce experimental validation that is often laborious or expensive, researchers demand reliable discoveries that contain few false discoveries. Accordingly, the false discovery rate (FDR) [14] has been developed as a statistical criterion for ensuring discoveries’ reliability. Technically, under the frequentist statistical paradigm, the FDR is defined as the expected proportion of uninteresting features among the discoveries. In parallel, under the Bayesian statistical paradigm, other criteria have been developed, including the Bayesian false discovery rate [15], the local false discovery rate (local fdr) [16], and the local false sign rate [17]. Among all these frequentist and Bayesian criteria, the FDR is the dominant criterion for setting thresholds in biological data analysis [1, 10, 18–24] and is thus the focus of this paper.

FDR control refers to the goal of finding discoveries such that the FDR is under a pre-specified threshold (e.g., 0.05). Existing computational methods for FDR control primarily rely on p-values, one per feature. Among the p-value-based FDR control methods, the most classic and popular ones are the Benjamini-Hochberg (BH) procedure [14] and the Storey’s q-value [25]; later development introduced methods that incorporate feature weights [26] and/or covariates—e.g., independent hypothesis weighting (IHW) [27], adaptive p-value thresholding [28], and Boca and Leek’s FDR regression [29]—to boost the detection power. All these methods set a p-value cutoff based on the pre-specified FDR threshold. However, the calculation of p-values requires either distributional assumptions, which are often questionable, or large numbers of replicates, which are often unachievable in biological studies (see Results). Due to the difficulty of valid p-value calculation in high-throughput biological data analysis, bioinformatics tools often output ill-posed p-values. This issue is evidenced by serious concerns about the widespread miscalculation and misuse of p-values in the scientific community [30]. As a result, bioinformatics tools using questionable p-values either cannot reliably control the FDR to a target level [23] or lack power to make discoveries [31]; see Results. Therefore, p-value-free control of FDR is desirable, as it would make data analysis more transparent and thus improve the reproducibility of scientific research.

Although p-value-free FDR control has been implemented in the MACS2 method for ChIP-seq peak calling [1] and the SAM method for microarray DEG identification [32], these two methods are restricted to specific applications and lack a theoretical guarantee for FDR control. (Although later works have studied some theoretical properties of SAM, they are not about the exact control of the FDR [33, 34].) More recently, the Barber-Candès (BC) procedure has been proposed to achieve theoretical FDR control without using p-values [35], and it has been shown to perform comparably to the BH procedure using well-calibrated p-values [36]. The BC procedure is advantageous because it does not require well-calibrated p-values, so it holds tremendous potential in various high-throughput data analyses where p-value calibration is challenging [37]. For example, a recent paper has implemented a generalization of the BC procedure to control the FDR in peptide identification from MS data [38].

Inspired by the BC procedure, we propose a general statistical framework Clipper to provide reliable FDR control for high-throughput biological data analysis, without using p-values or relying on specific data distributions. Clipper is a robust and flexible framework that applies to both enrichment and differential analyses and that works for high-throughput data with various characteristics, including data distributions, replicate numbers (from one to multiple), and outlier existence.

## Results

Clipper consists of two main steps: construction and thresholding of contrast scores. First, Clipper defines a contrast score for each feature, as a replacement of a p-value, to summarize that feature’s measurements between two conditions and to describe the degree of interestingness of that feature. Second, as its name suggests, Clipper establishes a cutoff on features’ contrast scores and calls as discoveries the features whose contrast scores exceed the cutoff (see Methods and Additional File 1: Section S2). Clipper is a flexible framework that only requires a minimal input: all features’ measurements under two conditions and a target FDR threshold (e.g., 5%) (Fig. 1b).

Clipper only relies on two fundamental statistical assumptions of biological data analysis: (1) measurement errors (i.e., differences between measurements and their expectations, with the expectations including both biological signals and batch effects) are independent across all features and replicates; (2) every uninteresting feature has measurement errors identically distributed across all replicates under both conditions. These two assumptions are used in almost all bioinformatics tools and are commonly referred to as the “measurement model” in statistical genomics [39]. With these two assumptions, Clipper has a theoretical guarantee for FDR control under both enrichment and differential analyses with any number of replicates (see Methods and Additional File 1: Section S2).

To verify Clipper’s performance, we designed comprehensive simulation studies to benchmark Clipper against existing generic FDR control methods (Additional File 1: Section S1). We also benchmarked Clipper against bioinformatics tools in studies including peak calling from ChIP-seq data, peptide identification from mass spectrometry data, DEG identification from bulk and single-cell RNA-seq data, and DIR identification from Hi-C data. Notably, our benchmarking results for peptide identification are based on our in-house data, the first MS data standard with a realistic dynamic range.

### Clipper has verified FDR control and power advantage in simulation

Simulation is essential because we can generate numerous datasets from the same distribution with known truths to calculate the FDR, which is not observable from real data. Our simulation covers both enrichment and differential analyses. In enrichment analysis, we consider four “experimental designs”: 1vs1 design (one replicate per condition), 2vs1 design (two and one replicates under the experimental and background conditions, respectively), 3vs3 design (three replicates per condition), and 10vs10 design (ten replicates per condition). In differential analysis, since Clipper requires that at least one condition has two replicates, we only consider the 2vs1 and 3vs3 designs. For each analysis and design, we simulated data from three “distributional families”—Gaussian, Poisson, and negative binomial—for individual features under two “background scenarios” (i.e., scenarios of the background condition): homogeneous and heterogeneous. Under the homogeneous scenario, all features’ measurements follow the same distribution under the background condition; otherwise, we are under the heterogeneous scenario, which is ubiquitous in applications, e.g., identifying DEGs from RNA-seq data and calling protein-binding sites from ChIP-seq data. By simulation setting, we refer to a combination of an experimental design, a distributional family, and a background scenario. The details of simulation settings are described in Additional File 1: Section S4.

For both enrichment and differential analyses and each simulation setting, we compared Clipper against generic FDR control methods, including p-value-based methods and local-fdr-based methods. The p-value-based methods include BH-pair, BH-pool, qvalue-pair, and qvalue-pool, where “BH” and “qvalue” stand for p-value thresholding procedures, and “pair” and “pool” represent the paired and pooled p-value calculation approaches, respectively. The local-fdr-based methods include locfdr-emp and locfdr-swap, where “emp” and “swap” represent the empirical null and swapping null local-fdr calculation approaches, respectively. See Methods for detail.

The comparison results are in Fig. 2 and Additional File 1: Figs. S1–S14. A good FDR control method should have its actual FDR no larger than the target FDR threshold and achieve high power. The results show that Clipper controls the FDR and is overall more powerful than the other methods, excluding those that fail to control the FDR, under all settings. Clipper is also shown to be more robust to the number of features and the existence of outliers than the other methods. In detail, in both enrichment analyses (1vs1, 2vs1, 3vs3, and 10vs10 designs) and differential analyses (2vs1 and 3vs3 designs), Clipper consistently controls the FDR, and it is more powerful than the generic methods in most cases under the realistic, heterogeneous background, where features do not follow the same distribution under the background condition. Under the idealistic, homogeneous background, Clipper is still powerful and only second to BH-pool and qvalue-pool, which, however, cannot control the FDR under the heterogeneous background.

**Figure 2.**
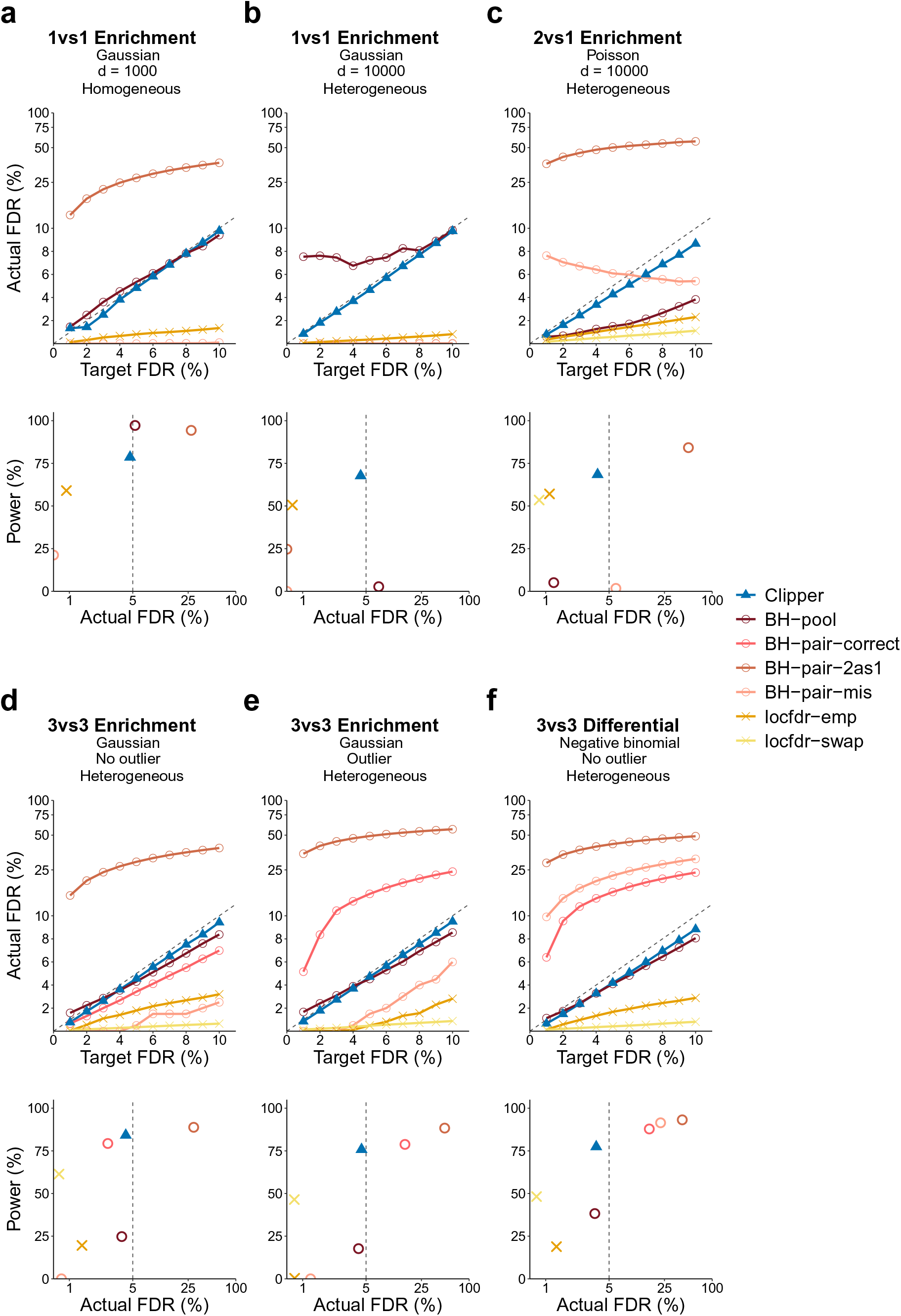
Comparison of Clipper with generic FDR control methods in terms of their FDR control and power in six example simulation studies. **(a)** 1vs1 enrichment analysis with 1000 features generated from the Gaussian distribution with a homogeneous background; **(b)** 1vs1 enrichment analysis with 10,000 features generated from the Gaussian distribution with a heterogeneous background; **(c)** 2vs1 enrichment analysis with 10,000 features generated from the Poisson distribution with a heterogeneous background; **(d)** 3vs3 enrichment analysis with 10,000 features generated from the Gaussian distribution without outliers and with a heterogeneous background; **(e)** 3vs3 enrichment analysis with 10,000 features generated from the Gaussian distribution with outliers and with a heterogeneous background; **(f)** 3vs3 differential analysis with 10,000 features generated from the negative binomial distribution with a heterogeneous background. At target FDR thresholds *q* ∈ {1%, 2%, … , 10%}, each method’s actual FDRs and power are approximated by the averages of false discovery proportions (see Additional File 1: Eq. (S14)) and power evaluated on 200 simulated datasets. In each panel, the top row shows each method’s actual FDRs at target FDR thresholds: whenever the actual FDR is larger than the target FDR (the solid line is higher than the dashed line), FDR control is failed; the bottom row shows each method’s actual FDRs and power at the target FDR threshold *q* = 5%: whenever the actual FDR is greater than *q* (on the right of the vertical dashed line), FDR control is failed. Under the FDR control, the larger the power, the better. Note that BH-pair-correct is not included in (a)–(c) because it is impossible to correctly specify the model with only one replicate per condition; locfdr-swap is not included in (a)–(b) because it is inapplicable to the 1vs1 design.

Here we summarize the performance of the generic FDR control methods. First, the two p-value-based methods using the pooled approach—BH-pool and qvalue-pool—are the most powerful under the idealistic, homogeneous background, which is their inherent assumption; however, they cannot control the FDR under the heterogeneous background (Fig. 2b). Besides, they cannot control the FDR when the number of features is small (Fig. 2a and Additional File 1: Fig. S1). These results show that the validity of BH-pool and qvalue-pool requires a large number of features and the homogeneous background assumption, two requirements that rarely both hold in biological applications.

Second, the four p-value-based methods using the paired approach with misspecified models or misformulated tests (BH-pair-mis, qvalue-pair-mis, BH-pair-2as1, and qvalue-pair-2as1; see Methods) fail to control the FDR by a large margin in most cases, and rarely when they control the FDR, they lack power (Fig. 2c–d and Additional File 1: Figs. S1–S8). These results confirm that BH-pair and qvalue-pair rely on the correct model specification to control the FDR; however, the correct model specification is hardly achievable with no more than three replicates per condition.

Third, even when models are correctly specified (an idealistic scenario), the p-value-based methods that use the paired approach—BH-pair-correct and qvalue-pair-correct (see Methods)—fail to control the FDR in the existence of outliers (Fig. 2e and Additional File 1: Figs. S3 and S7) or for the negative binomial distribution with unknown dispersion (Fig. 2f and Additional File 1: Fig. S9). It is worth noting that even when they control the FDR, they are less powerful than Clipper in most cases except for the 3vs3 differential analysis with the Poisson distribution (Fig. 2d and Additional File 1: Figs. S4 and S8).

Fourth, the two local-fdr-based methods—locfdr-emp and locfdr-swap—achieve the FDR control for all designs and analyses; however, they are less powerful than Clipper in most cases (Additional File 1: Figs. S1–S4).

Fifth, when the numbers of replicates are large (10vs10 design), non-parametric tests become applicable. We compared Clipper with three BH-pair methods that use different statistical tests: BH-pair-Wilcoxon (the non-parametric Wilcoxon rank-sum test), BH-pair-permutation (the non-parametric permutation test), and BH-pair-parametric (the parametric test based on the correct model specification, equivalent to BH-pair-correct). Although all the three methods control the FDR, they are less powerful than Clipper (Additional File 1: Fig. S10).

Moreover, the above five phenomena are consistently observed across the three distributions (Gaussian, Poission, and negative binomial) that we have examined, further confirming the robustness of Clipper.

In addition, for the 3vs3 enrichment analysis, we also varied the proportion of interesting features as 10%, 20%, and 40%. The comparison results in Additional File 1: Fig. S3 (columns 1 and 3 for 10%) and Additional File 1: Fig. S11 (for 20% and 40%) show that the performance of Clipper is robust to the proportion of interesting features.

The above results are all based on simulations with independent features. To examine the robustness of Clipper, we introduced feature correlations to our simulated data, on which we compared Clipper with the other generic FDR control methods. The comparison results in Additional File 1: Fig. S12 show that even when the feature independence assumption is violated, Clipper still demonstrates strong performance in both FDR control and power.

### Clipper has broad applications in omics data analyses

We then demonstrate the use of Clipper in four omics data applications: peak calling from ChIP-seq data, peptide identification from MS data, DEG identification from bulk or single-cell RNA-seq data, and DIR identification from Hi-C data. The first two applications are enrichment analyses, and the last two are differential analyses. In each application, we compared Clipper with mainstream bioinformatics methods to demonstrate Clipper’s superiority in FDR control and detection power.

#### Peak calling from ChIP-seq data (enrichment analysis I)

ChIP-seq is a genome-wide experimental assay for measuring binding intensities of a DNA-associated protein [40], often a transcription factor that activates or represses gene expression [41, 42]. ChIP-seq data are crucial for studying gene expression regulation, and an indispensable analysis is to identify genomic regions with enriched sequence reads in ChIP-seq data. These regions are likely bound by the target protein and thus of biological interest. The identification of these regions is termed “peak calling” in ChIP-seq data analysis.

As the identified peaks are subject to experimental validation that is often expensive [43], it is essential to control the FDR of peak identification to reduce unnecessary costs. The two most highly-cited peak-calling methods are MACS2 [1] and HOMER [2], both of which claim to control the FDR for their identified peaks. Specifically, both MACS2 and HOMER assume that the read counts for each putative peak (with one count per sample/replicate) follow the Poisson distribution, and they use modified paired approaches to assign each putative peak a p-value and a corresponding Storey’s q-value. Then given a target FDR threshold 0 < *q* < 1, they call the putative peaks with q-values ≤ *q* as identified peaks. Despite being popular, MACS2 and HOMER have not been verified for their FDR control, to our knowledge.

To verify the FDR control of MACS2 and HOMER (Additional File 1: Section S5.1), we used ENCODE ChIP-seq data of cell line GM12878 [44] and ChiPulate [45], a ChIP-seq data simulator, to generate semi-synthetic data with spiked-in peaks (Additional File 1: Section S6.1). We examined the actual FDR and power of MACS2 and HOMER in a range of target FDR thresholds: *q* = 1%, 2%, … , 10%. Fig. 3a shows that MACS2 and HOMER cannot control the FDR as standalone peak-calling methods. However, with Clipper as an add-on (Additional File 1: Section S7.1), both MACS2 and HOMER can guarantee the FDR control. This result demonstrates the flexibility and usability of Clipper for reducing false discoveries in peak calling analysis.

**Figure 3.**
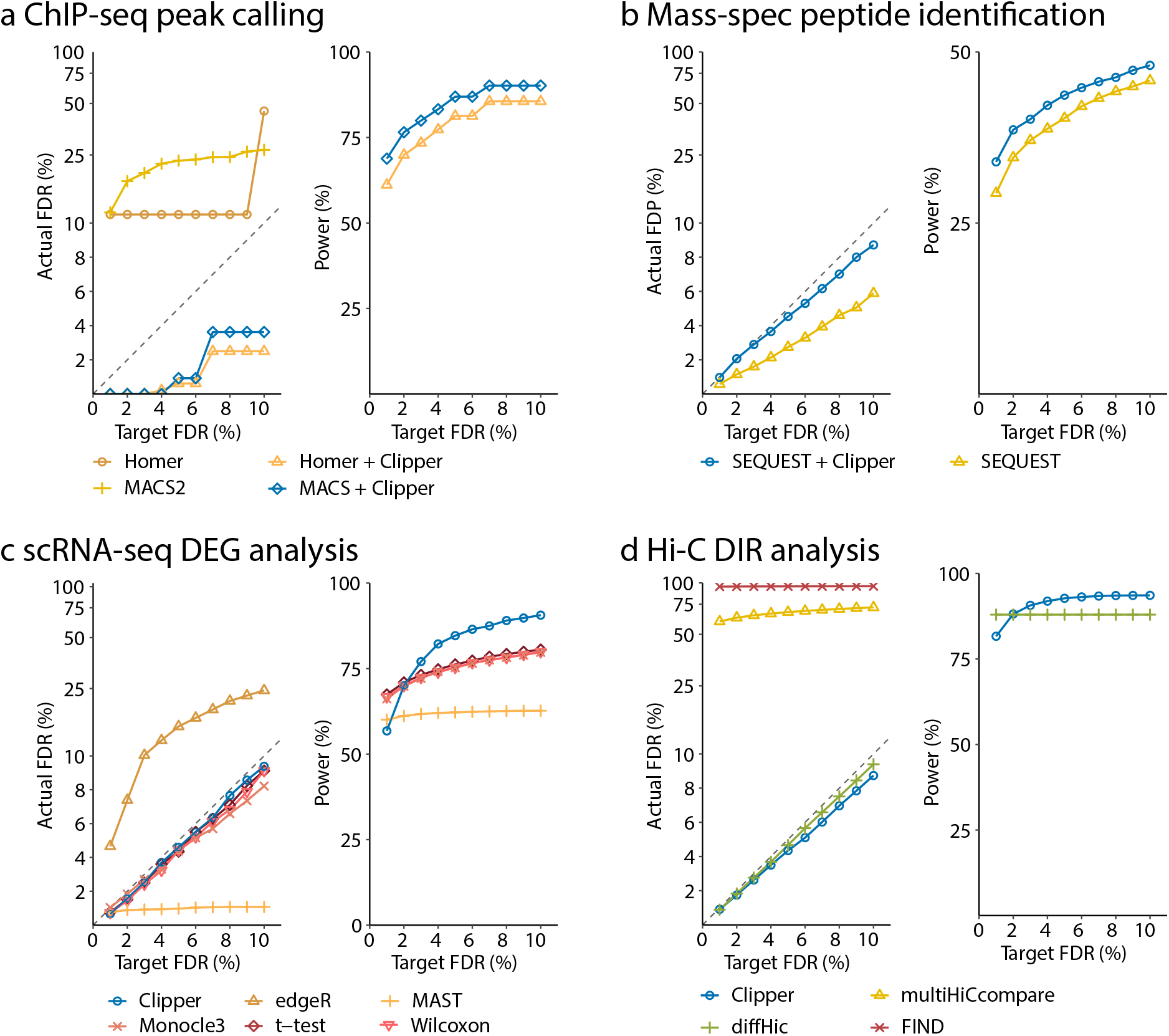
Comparison of Clipper and popular bioinformatics methods in terms of FDR control and power. **(a)** peaking calling analysis on semi-synthetic ChIP-seq data; **(b)** peptide identification on real proteomics data; **(c)** DEG analysis on semi-synthetic 10x Genomics scRNA-seq data; **(d)** DIR analysis on semi-synthetic Hi-C data. In all four panels, the target FDR threshold *q* ranges from 1% to 10%. In the “Actual FDR vs. Target FDR” plot of each panel, points above the dashed diagonal line indicate failed FDR control; when this happens, the power of the corresponding methods is not shown, including MACS2 and HOMER in (a), edgeR in (c), and multiHICcompare and FIND in (d). In all four applications, Clipper controls the FDR while maintaining high power, demonstrating Clipper’s broad applicability in high-throughput data analyses.

Technically, the failed FDR control by MACS2 and HOMER is attributable to the likely model misspecification and test misformulation in their use of the paired approach. Both MACS2 and HOMER assume the Poisson distribution for read counts in a putative peak; however, it has been widely acknowledged that read counts are over-dispersed and thus better modeled by the negative binomial distribution [46]. Besides, MACS2 uses one-sample tests to compute p-values when two-sample tests should be performed. As a result, the p-values of MACS2 and HOMER are questionable, so using their p-values for FDR control has no guaranteed success. (Note that MACS2 does not use p-values to control the FDR but instead swaps experimental and background samples to calculate the empirical FDR; yet, we emphasize that controlling the empirical FDR does not guarantee the FDR control.) As a remedy, Clipper strengthens both methods to control the FDR while maintaining high power.

As a side note, it is known that uninteresting regions tend to have larger read counts in the control sample than in the experimental (ChIP) sample; however, this phenomenon does not violate Clipper’s theoretical assumption for FDR control (Additional File 1: Section S7.1).

#### Peptide identification from MS data (enrichment analysis II)

The state-of-the-art proteomics studies use MS experiments and database search algorithms to identify and quantify proteins in biological samples. In a typical proteomics experiment, a protein mixture sample is first digested into peptides and then measured by tandem MS technology as mass spectra, which encode peptide sequence information. “Peptide identification” is the process that decodes mass spectra and converts mass spectra into peptide sequences in a protein sequence database via search algorithms. The search process matches each mass spectrum to peptide sequences in the database and outputs the best match, called a “peptide-spectrum match” (PSM). The identified PSMs are used to infer and quantify proteins in a high-throughput manner.

False PSMs could occur when mass spectra are matched to wrong peptide sequences due to issues such as low-quality spectra, data-processing errors, and incomplete protein databases, causing problems in the downstream protein identification and quantification [47]. Therefore, a common goal of database search algorithms is to simultaneously control the FDR and maximize the number of identified PSMs, so as to maximize the number of proteins identified in a proteomics study [3, 48, 49]. A widely used FDR control strategy is the target-decoy search, where mass spectra of interest are matched to peptide sequences in both the original (target) database and a decoy database that contains artificial false protein sequences. The resulting PSMs are called the target PSMs and decoy PSMs, respectively. The decoy PSMs, i.e., matched mass spectrum and decoy peptide pairs, are known to be false and thus used by database search algorithms to control the FDR. Mainstream database search algorithms output a q-value for each target or decoy PSM. Discoveries are the target PSMs whose q-values are no greater than the target FDR threshold *q*.

We generated the first comprehensive benchmark dataset from an archaea species *Pyrococcus furiosus*, and we used it to examine the FDR control and power of a popular database search algorithm SEQUEST [3] (Additional File 1: Section S5.2). Using this benchmark dataset (Additional File 1: Section S6.2), we demonstrate that, as an add-on, Clipper improves the power of SEQUEST. Specifically, Clipper treats mass spectra as features. For each mass spectrum, Clipper considers the experimental (or background) measurement as the − log_10_-transformed SEQUEST q-value of the target (or decoy) PSM that includes the mass spectrum. Then Clipper decides which mass spectra and their corresponding target PSMs are discoveries (Additional File 1: Section S7.2). Based on the benchmark dataset, we examined the empirical FDR, i.e., the FDP calculated based on the true positives and negatives, and the power of SEQUEST with or without Clipper as an add-on, for a range of target FDR thresholds: *q* = 1%, 2%, … , 10%. Fig. 3b shows that although SEQUEST and SEQUEST+Clipper both control the FDR, SEQUEST+Clipper consistently improves the power, thus enhancing the peptide identification efficiency of proteomics experiments.

While preparing this manuscript, we found a recent work [38] that used a similar idea to identify PSMs without using p-values. Clipper differs from this work in two aspects: (1) Clipper is directly applicable as an add-on to any existing database search algorithms that output q-values; (2) Clipper is not restricted to the peptide identification application.

#### DEG identification from bulk RNA-seq data (differential analysis I)

RNA-seq data measure genome-wide gene expression levels in biological samples. An important use of RNA-seq data is the DEG analysis, which aims to discover genes whose expression levels change between two conditions. The FDR is a widely used criterion in DEG analysis [4–9].

We compared Clipper with two popular DEG identification methods: edgeR [4] and DESeq2 [5] (Additional File 1: Section S5.3). Specifically, when we implemented Clipper, we first performed the trimmed mean of M values (TMM) normalization [50] to correct for batch effects; then we treated genes as features and their normalized expression levels as measurements under two conditions (Additional File 1: Section S7.3). We also implemented two versions of DESeq2 and edgeR: with or without IHW, a popular procedure for boosting the power of p-value-based FDR control methods by incorporating feature covariates [27]. In our implementation of the two versions of DESeq2 and edgeR, we used their standard pipelines, including normalization, model fitting, and gene filtering (edgeR only). To verify the FDR control, we generated four realistic semi-synthetic datasets from two real RNA-seq datasets—one from classical and non-classical human monocytes [51] and the other from yeasts with or without *snf2* knockout [52]—using simulation strategies 1 and 2 (Additional File 1: Section S6.3).

In detail, in simulation strategy 1, we used bulk RNA-seq samples from two conditions to compute a fold change for every gene between the two conditions; then we defined true DEGs as the genes whose fold changes exceeded a threshold; next, we randomly drew three RNA-seq samples and treated them as replicates from each condition (*m* = *n* = 3 as in Methods); using those subsampled replicates of two conditions, we preserved the true DEGs’ read counts and permuted the read counts of the true non-DEGs, i.e., the genes other than true DEGs, between conditions. In summary, simulation strategy 1 guarantees that the measurements of true non-DEGs are i.i.d., an assumption that Clipper relies on for theoretical FDR control.

In simulation strategy 2, which we borrowed from a benchmark study [53], we first randomly selected at most 30% genes as true DEGs; next, we randomly drew six RNA-seq samples from one condition (classical human monocytes and yeasts without knockout) and split the samples into two “synthetic conditions,” each with three replicates (*m* = *n* = 3 as in Methods); then for each true DEG, we multiplied its read counts under one of the two synthetic conditions (randomly and independently picked for each gene) by a randomly generated fold change (see Additional File 1: Section S6.3); finally, for the true non-DEGs, we preserved their read counts in the six samples. In summary, simulation strategy 2 preserves batch effects, if existent in real data, for the true non-DEGs (the majority of genes). As a result, the semi-synthetic data generated under strategy 2 may violate the Clipper assumption for theoretical FDR control and thus can help evaluate the robustness of Clipper on real data.

The four semi-synthetic datasets have ground truths (true DEGs and non-DEGs) to evaluate each DEG identification method’s FDR and power for a range of target FDR thresholds: *q* = 1%, 2%, … , 10%. Our results in Fig. 4a and Additional File 1: Figs. S15a–S17a show that Clipper consistently controls the FDR and achieves high power on all four semi-synthetic datasets. In contrast, DESeq2 and edgeR cannot consistently control the FDR except for the yeast semi-synthetic dataset generated under simulation strategy 2. Given the fact that DESeq2 and edgeR do not consistently perform well on the three other semi-synthetic datasets, we hypothesize that their parametric distributional assumptions, if violated on real data, hinder valid FDR control, in line with our motivation for developing Clipper. By examining whether true non-DEGs’ p-values calculated by DESeq2 or edgeR follow the theoretical Uniform[0, 1] distribution, we find that the answer is no for many non-DEGs, as indicated by the small p-values (one per non-DEG) of uniformity tests (Additional File 1: Fig. S18); this issue is more serious for DESeq2, consistent with the worse FDR control of DESeq2 (Fig. 4a and Additional File 1: Figs. S15a–S17a). Furthermore, we observe that adding IHW to edgeR and DESeq2 has negligible effects on the four semi-synthetic datasets.

**Figure 4.**
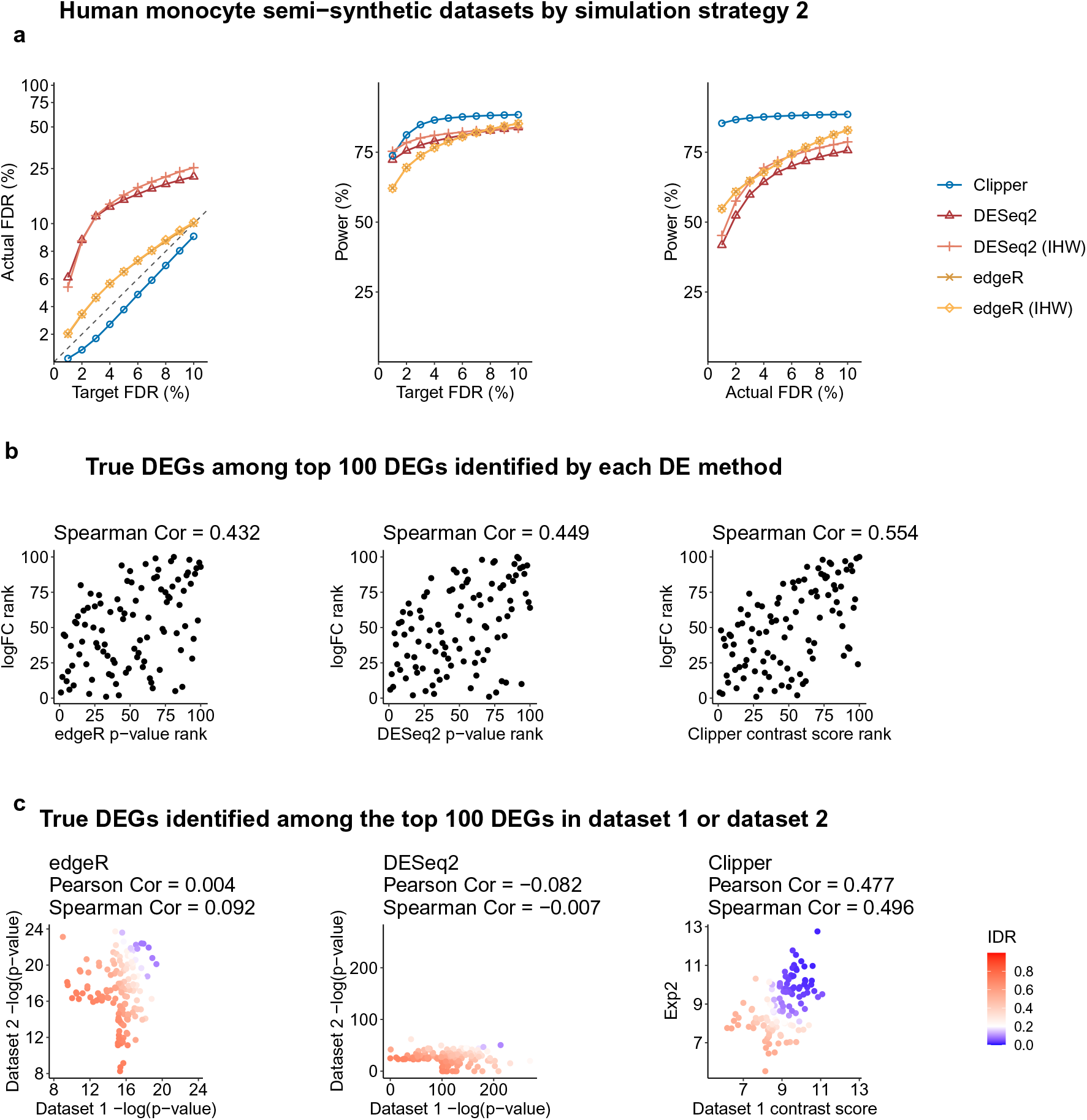
Comparison of Clipper and two popular DEG identification methods—edgeR and DESeq2—in DEG analysis on semi-synthetic bulk RNA-seq data (generated from human monocyte real data using simulation strategy 2 in Additional File 1: Section S6.3). **(a)** FDR control, power given the same target FDR, and power given the same actual FDR. **(b)** Ranking consistency of the true DEGs among the top 100 DEGs identified by each method. The consistency is defined between the genes’ ranking based on edgeR/DESeq2’s p-values or Clipper’s contrast scores and their ranking based on true expression fold changes. **(c)** Reproducibility between two semi-synthetic datasets as technical replicates. Three reproducibility criteria are used: the IDR, the Pearson correlation, and the Spearman correlation. Each criterion is calculated for edgeR/DESeq2’s p-values or Clipper’s contrast scores on the two semi-synthetic datasets. Among the three methods, only Clipper controls the FDR, and Clipper achieves the highest power, the best gene ranking consistency, and the best reproducibility.

To further explain why DESeq2 fails to control the FDR, we examined the p-value distributions of 16 non-DEGs that were most frequently identified (from the 100 semi-synthetic datasets generated from the human monocyte dataset using simulation strategy 1) by DESeq2 at the target FDR threshold *q* = 0.05. Our results in Additional File 1: Fig. S19 show that the 16 non-DEGs’ p-values are non-uniformly distributed with a mode close to 0. Such unusual enrichment of overly small p-values makes these non-DEGs mistakenly called as discoveries by DESeq2.

In addition, we compared the DEG ranking by Clipper, edgeR, and DESeq2 in two ways. First, for true DEGs, we compared their ranking by each method with their true ranking based on true expression fold changes (from large to small, as in semi-synthetic data generation in Additional File 1: Section S6.3). Specifically, we ranked true DEGs using Clipper’s contrast scores (from large to small), edgeR’s p-values (from small to large), or DESeq2’s p-values (from small to large). Our results in Fig. 4b and Additional File 1: Figs. S15b–S17b show that Clipper’s contrast scores exhibit the most consistent ranking with the ranking based on true fold changes. Second, to compare the power of Clipper, edgeR, and DESeq2 based on their DEG rankings instead of nominal p-values, we calculated their power under the actual FDRs, which only depend on gene rankings (for the definition of actual FDR, see Additional File 1: Section S6.3). Fig. 4a and Additional File 1: Figs. S15a–S17a show that, when Clipper, edgeR, and DESeq2 have the same actual FDR, Clipper consistently outperforms edgeR and DESeq2 in terms of power, i.e., Clipper has the largest number of true DEGs in its top ranked genes.

We also compared the reproducibility of Clipper, edgeR, and DESeq2 in the presence of sampling randomness. Specifically, we used two semi-synthetic datasets (generated independently from the same procedure in Additional File 1: Section S6.3) as technical replicates and computed Clipper’s contrast scores, edgeR’s p-values, and DESeq’s p-values on each dataset. For each method, we evaluated its reproducibility between the two semi-synthetic datasets by computing three criteria—the irreproducibility discovery rate (IDR) [54], the Pearson correlation, and the Spearman correlation—using its contrast scores or − log_10_-transformed p-values. Fig. 4c and Additional File 1: Figs. S15c–S17c show that Clipper’s contrast scores have higher reproducibility by all three criteria compared to edgeR’s and DESeq2’s p-values.

Finally, we compared Clipper with DESeq2 and edgeR on the real RNA-seq data of classical and non-classical human monocytes [51]. In this dataset, gene expression changes are expected to be associated with the immune response process. We input three classical and three non-classical samples into Clipper, DESeq2, and edgeR for DEG identification. Fig. 5a shows that edgeR identifies the fewest DEGs, while DESeq2 identifies the most DEGs, followed by Clipper. Notably, most DEGs identified by DESeq2 are not identified by Clipper or edgeR. To investigate whether DESeq2 makes too many false discoveries and whether Clipper finds biologically meaningful DEGs missed by DESeq2 or edgeR, we performed functional analysis on the set of DEGs identified by each method. We first performed the gene ontology (GO) analysis on the three sets of identified DEGs using the R package `clusterProfiler` [55]. Fig. 5b (“Total”) shows that more GO terms are enriched (with enrichment q-values ≤ 0.01) in the DEGs identified by Clipper than in the DEGs identified by DESeq2 or edgeR. For the GO terms enriched in all three sets of identified DEGs, Fig. 5c shows that they are all related to the immune response and thus biologically meaningful. Notably, these biologically meaningful GO terms have more significant enrichment in Clipper’s identified DEGs than in edgeR and DESeq2’s identified DEGs. We further performed GO analysis on the DEGs uniquely identified by one method in pairwise comparisons of Clipper vs. DESeq2 and Clipper vs. edgeR. Fig. 5b and Additional File 1: Fig. S20 show that multiple immune-related GO terms are enriched in Clipper-specific DEGs, while no GO terms are enriched in edgeR-specific or DESeq2-specific DEGs. In addition, we examined the DEGs that were identified by Clipper only but missed by both edgeR and DESeq2. Fig. 5d and Additional File 2 show that these genes include multiple key immune-related genes, including *CD36*, *DUSP2*, and *TNFAIP3*. We further performed pathway analysis on these genes and the DEGs that were identified by DEseq2 only but missed by both edgeR and Clipper, using the R package `limma` [10]. Additional File 1: Fig. S21a shows that the DEGs that were only identified by Clipper have significant enrichment for immune-related pathways including phagosome, a key function of monocytes and macrophages. On the contrary, Additional File 1: Fig. S21b shows that fewer immune-related pathways are enriched in DEGs that were only identified by DESeq2. Altogether, these results confirm the capacity of Clipper in real-data DEG analysis, and they are consistent with our simulation results that edgeR lacks power, while DESeq2 fails to control the FDR.

**Figure 5.**
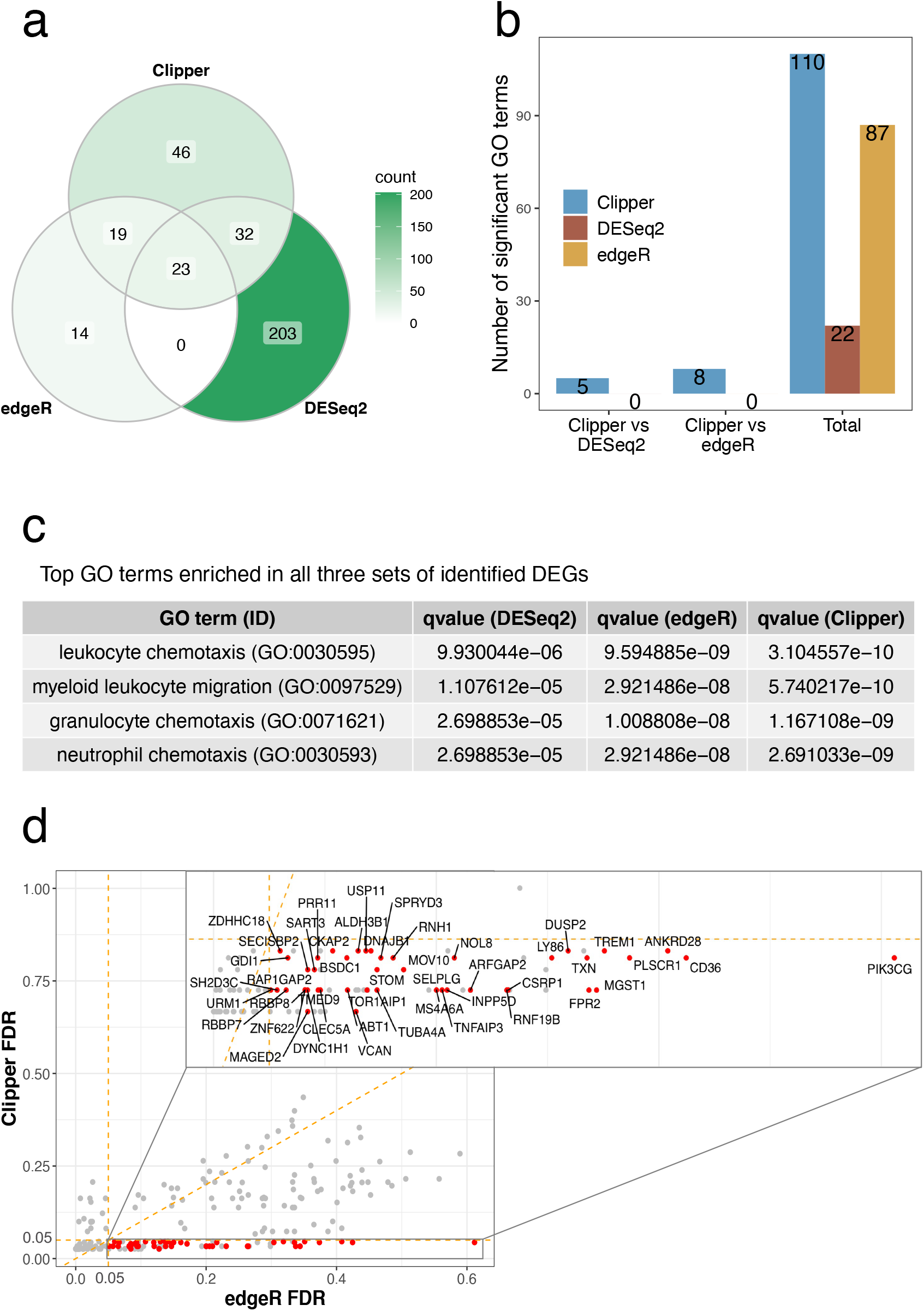
Application of Clipper, DESeq2, and edgeR to identifying DEGs from the classical and non-classical human monocyte dataset. **(a)** A Venn diagram showing the overlaps of the identified DEGs (at the FDR threshold *q* = 5%) by the three DE methods. **(b)** Numbers of GO terms enriched (with enrichment q-values < 0.01) in the DEGs found by Clipper, DESeq2 and edgeR (column 3), or in the DEGs specifically identified by Clipper or DESeq2/edgeR in the pairwise comparison between Clipper and DESeq2 (column 1) or between Clipper and edgeR (column 2). More GO terms are enriched in the DEGs identified by Clipper than in those identified by edgeR or DESeq2. **(c)** Enrichment q-values of four GO terms that are found enriched (with enrichment q-values < 0.01) in all three sets of identified DEGs, one set per method. All the four terms are most enriched in the DEGs identified by Clipper. **(d)** A scatterplot of the claimed FDR of Clipper against that of edgeR for all the DEGs identified by Clipper, edgeR or DESeq2. The 46 DEGs identified by Clipper only at 5% FDR are highlighted in red.

#### DEG identification from single-cell RNA-seq data (differential analysis II)

Single-cell RNA sequencing (scRNA-seq) technologies have revolutionized biomedical sciences by enabling genome-wide profiling of gene expression levels at an unprecedented single-cell resolution. DEG analysis is widely applied to scRNA-seq data for discovering genes whose expression levels change between two conditions or between two cell types. Compared with bulk RNA-seq data, scRNA-seq data have many more “replicates” (i.e., cells, whose number is often in hundreds) under each condition or within each cell type.

We compared Clipper (Additional File 1: Section S7.4) with edgeR [4], MAST [56], Monocle3 [57], the two-sample *t* test, and the Wilcoxon rank-sum test (Additional File 1: Section S5.4), five methods that are either popular or reported to have comparatively top performance from a previous benchmark study [58]. To verify the FDR control, we used scDesign2, a flexible probabilistic simulator, to generate scRNA-seq count data with known true DEGs [59]. scDesign2 offers three key advantages that enable the generation of realistic semi-synthetic scRNA-seq count data: (1) it captures distinct marginal distributions of different genes; (2) it preserves gene-gene correlations; (3) it adapts to various scRNA-seq protocols. Using scDesign2, we generated two semi-synthetic scRNA-seq datasets from two real scRNA-seq datasets of peripheral blood mononuclear cells (PBMCs) [60]: one using 10x Genomics [61] and the other using Drop-seq [62]. Each semi-synthetic dataset contains two cell types, CD4+ T cells and cytotoxic T cells, which we treated as two conditions (Additional File 1: Section S6.4). Having true DEGs known, the semi-synthetic datasets allow us to evaluate the FDRs and power of Clipper and the other five methods for a range of target FDR thresholds: *q* = 1%, 2%, … , 10%. Fig. 3c and Additional File 1: Fig. S22 show that on both 10x Genomics and Drop-seq semi-synthetic datasets, Clipper consistently controls the FDR and remains the most powerful (except for *q* = 1% and 2%) among all the methods that achieve FDR control. These results demonstrate Clipper’s robust performance in scRNA-seq DEG analysis.

#### DIR analysis of Hi-C data (differential analysis III)

Hi-C experiments are widely used to investigate spatial organizations of chromosomes and to map chromatin interactions across the genome. A Hi-C dataset is often processed and summarized into an interaction matrix, whose rows and columns represent manually binned chromosomal regions and whose (*i, j*)-th entry represents the measured contact intensity between the *i*-th and *j*-th binned regions. The DIR analysis aims to identify pairs of genomic regions whose contact intensities differ between conditions. Same as DEG analysis, DIR analysis also uses the FDR as a decision criterion [11–13].

We compared Clipper with three popular DIR identification methods: diffHic [13], FIND [12], and multiHiCcompare [11] (Additional File 1: Section S5.5). Specifically, we applied Clipper to DIR identification by treating pairs of genomic regions as features and their contact intensities as measurements. To verify the FDR control of Clipper (Additional File 1: Section S7.5), diffHic, FIND, and multiHiCcompare, we generated realistic semi-synthetic data from real interaction matrices of ENCODE cell line GM12878 [44] with true spiked-in DIRs to evaluate the FDR and power (Additional File 1: Section S6.5). We examined the actual FDR and power in a range of target FDR thresholds: *q* = 1%, 2%, … , 10%. Fig. 3d shows that Clipper and diffHic are the only two methods that consistently control the FDR, while multiHiCcompare and FIND fail by a large margin. In terms of power, Clipper outperforms diffHic except for *q* = 1% and 2%, even though Clipper has not been optimized for Hi-C data analysis. This result demonstrates Clipper’s general applicability and strong potential for DIR analysis.

## Discussion

In this paper, we proposed a new statistical framework, Clipper, for identifying interesting features with FDR control from high-throughput data. Clipper avoids the use of p-values and makes FDR control more reliable and flexible. We used comprehensive simulation studies to verify the FDR control by Clipper under various settings. We demonstrate that Clipper outperforms existing generic FDR control methods by having higher power and greater robustness to model misspecification. We further applied Clipper to four popular bioinformatics analyses: peak calling from ChIP-seq data, peptide identification from MS data, DEG identification from RNA-seq data, and DIR identification from Hi-C data. Our results indicate that Clipper provides a powerful add-on to existing bioinformatics tools to improve the reliability of FDR control and thus the reproducibility of scientific discoveries.

Clipper’s FDR control procedures (BC and GZ procedures in Methods) are motivated by the Barber-Candès (BC)’s knockoff paper [35] and the Gimenez-Zou (GZ)’s multiple knockoff paper [63], but we do not need to construct knockoffs in enrichment analysis when two conditions have the same number of replicates; the reason is that the replicates under the background condition serve as natural negative controls. For differential analysis and enrichment analysis with unequal numbers of replicates, in order to guarantee the theoretical assumptions for FDR control, Clipper uses permutations instead of the complicated knockoff construction because Clipper only examines features marginally and does not concern about features’ joint distribution.

We validated the FDR control by Clipper using extensive and concrete simulations, including both model-based and real-data-based data generation with ground truths, which are widely used to validate newly developed computational frameworks [64]. In contrast, in most bioinformatics method papers, the FDR control was merely mentioned but rarely validated. Many of them assumed that using the BH procedure on p-values would lead to valid FDR control; however, the reality is often otherwise because p-values would be invalid when model assumptions were violated or the p-value calculation was problematic. Here we voice the importance of validating FDR control in bioinformatics method development, and we use this work as a demonstration. We believe that Clipper provides a powerful booster to this movement. As a p-value-free alternative to the classic p-value-based BH procedure, Clipper relies less on model assumptions and is thus more robust to model misspecifications, making it an appealing choice for FDR control in diverse high-throughput biomedical data analyses.

Clipper is a flexible framework that is easily generalizable to identify a variety of interesting features. The core component of Clipper summarizes each feature’s measurements under each condition into an informative statistic (e.g., the sample mean); then Clipper combines each feature’s informative statistics under two conditions into a contrast score to enable FDR control. The current implementation of Clipper only uses the sample mean as the informative statistic to identify the interesting features that have distinct expected values under two conditions. However, by modifying the informative statistic, we can generalize Clipper to identify the features that are interesting in other aspects, e.g., having different variances between two conditions. Regarding the contrast score, Clipper currently makes careful choices between two contrast scores, minus and maximum, based on the number of replicates and the analysis task (enrichment or differential).

Notably, Clipper achieves FDR control and high power using those two simple contrast scores, which are calculated for individual features without borrowing information from other features. However, Clipper does leverage the power of multiple testing by setting a contrast score threshold based on all features’ contrast scores. This is a likely reason why Clipper achieves good power even with simple contrast scores. An advantage of Clipper is that it allows other definitions of contrast scores, such as the two-sample *t* statistic that considers within-condition variances. Empirical evidence (Additional File 1: Figs. S13 and S14) shows that the Clipper variant using the two-sample *t* statistic is underpowered by the default Clipper, which uses the minus summary statistic (difference of two conditions’ sample means) as the contrast score in the 3vs3 enrichment analysis or as the degree of interestingness in the 3vs3 differential analysis (see Methods). Here is our current interpretation of this seemingly counter-intuitive result.

- First, both the minus statistic and the *t* statistic satisfy Clipper’s theoretical conditions (Lemmas 1 and 3 in Additional File 1: Section S2), which guarantee the FDR control by the BC and GZ procedures; this is confirmed in Additional File 1: Figs. S13 and S14. Hence, from the FDR control perspective, Clipper does not require the adjustment for within-condition variances by using a *t* statistic.
- Second, Clipper is different from the two-sample *t* test or the regression-based *t* test, where the *t* statistic was purposely derived as a pivotal statistic so that its null distribution (the *t* distribution) does not depend on unknown parameters. Since Clipper does not require a null distribution for each feature, the advantage of the *t* statistic being pivotal no longer matters.
- Third, the minus statistic only requires estimates of two conditions’ mean parameters, while the *t* statistic additionally requires estimates of the two conditions’ variances. Hence, when the sample sizes (i.e., the numbers of replicates) are small, the two more parameters that need estimation in the *t* statistic might contribute to the observed power loss of the Clipper *t* statistic variant. Indeed, the power difference between the two statistics diminishes as the sample sizes increase from 3vs3 in Additional File 1: Figs. S13–S14 to 10vs10 in Additional File 1: Fig. S10 (where we compared the default Clipper with BH-pair-parametric, which is based on the two-sample *t* test and is highly similar to the Clipper *t* statistic variant).
- Fourth, we observe empirically that a contrast score would have better power if its distribution (based on its values of all features) has a larger range and a heavier right tail (in the positive domain). Compared to the minus statistic, the *t* statistic has a smaller range and a lighter right tail due to its adjustment for features’ within-condition variances (Additional File 1: Fig. S23). This observation is consistent with the power difference of the two statistics.

Beyond our current interpretation, however, we admit that future studies are needed to explore alternative contrast scores and their power with respect to data characteristics and analysis tasks. Furthermore, we may generalize Clipper to be robust against sample batch effects by constructing the contrast score as a regression-based test statistic that has batch effects removed.

Our current version of Clipper allows the identification of interesting features between two conditions. However, there is a growing need to generalize our framework to identify features across more than two conditions. For example, temporal analysis of scRNA-seq data aims to identify genes whose expression levels change along cell pseudotime [31]. To tailor Clipper for such analysis, we could define a new contrast score that differentiates the genes with stationary expression (uninteresting features) from the other genes with varying expression (interesting features). Further studies are needed to explore the possibility of extending Clipper to the regression framework so that Clipper can accommodate data of multiple conditions or even continuous conditions, as well as adjusting for confounding covariates.

We have demonstrated the broad application potential of Clipper in various bioinformatics data analyses. Specifically, when used as an add-on to established, popular bioinformatics methods such as MACS2 for peak calling and SEQUEST for peptide identification, Clipper guaranteed the desired FDR control and in some cases boosted the power. However, many more careful thoughts are needed to escalate Clipper into standalone bioinformatics methods for specific data analyses, for which data processing and characteristics (e.g., peak lengths, GC contents, proportions of zeros, and batch effects) must be appropriately accounted for before Clipper is used for the FDR control [58, 65]. We expect that the Clipper framework will propel future development of bioinformatics methods by providing a flexible p-value-free approach to control the FDR, thus improving the reliability of scientific discoveries.

After finishing this manuscript, we were informed of the work by He et al. [66], which is highly similar to the part of Clipper for differential analysis, as both work use permutation for generating negative controls and the GZ procedure for thresholding (test statistics in He et al. and contrast scores in Clipper). However, the test statistics used in He et al. are the two-sample *t* statistic and the two-sample Wilcoxon statistic, both of which are different from the minus and maximum contrast scores used in Clipper. While we leave the optimization of contrast scores to future work, we observed that the minus contrast score outpowers the two-sample *t* statistic in our analysis (Additional File 1: Figs. S13 and S14), and we hypothesize that the two-sample Wilcoxon statistic, though being a valid contrast score for differential analysis, requires a large sample size to achieve good power. For this reason, we did not consider it as a contrast score in the current Clipper implementation, whose focus is on small-sample-size high-throughout biological data.

## Conclusion

In high-throughput biological data analysis, which aims to identify interesting features by comparing two conditions, existing bioinformatics tools control the FDR based on p-values. However, obtaining valid p-values relies on either reasonable assumptions of data distribution or large numbers of replicates under both conditions—two requirements that are often unmet in biological studies. To address this issue, we propose Clipper, a general statistical framework for FDR control without relying on p-values or specific data distributions. Clipper is applicable to identifying both enriched and differential features from high-throughput biological data of diverse types. In comprehensive simulation and real-data benchmarking, Clipper outperforms existing generic FDR control methods and specific bioinformatics tools designed for various tasks, including peak calling from ChIP-seq data, differentially expressed gene identification from bulk or single-cell RNA-seq data, differentially interacting chromatin region identification from Hi-C data, and peptide identification from mass spectrometry data. Our results demonstrate Clipper’s flexibility and reliability for FDR control, as well as its broad applications in high-throughput data analysis.

## Methods

### Clipper: notations and assumptions

We first introduce notations and assumptions used in Clipper. While the differential analysis treats the two conditions symmetric, the enrichment analysis requires one condition to be the experimental condition (i.e., the condition of interest) and the other condition to be the background condition (i.e., the negative control). For simplicity, we use the same set of notations for both analyses. For two random vectors ***X*** = (*X*_1_, … , *X*_*m*_)^⊤^ and ***Y*** = (*Y*_1_, … , *Y*_*n*_)^⊤^, we write ***X*** ⊥ ***Y*** if *X*_*i*_ is independent of *Y*_*j*_ for all *i* = 1, … , *m* and *j* = 1, … , *n*. To avoid confusion, we use card(*A*) to denote the cardinality of a set *A* and |*c*| to denote the absolute value of a scalar *c*. We define *a* ∨ *b* := max(*a, b*).

Clipper only requires two inputs: the target FDR threshold *q* ∈ (0, 1) and the input data. Regarding the input data, we use *d* to denote the number of features with measurements under two conditions, and we use *m* and *n* to denote the numbers of replicates under the two conditions. For each feature *j* = 1, … , *d*, we use 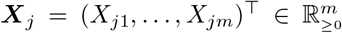 and 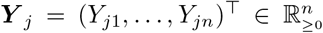 to denote its measurements under the two conditions, where ℝ_≥0_ denotes the set of non-negative real numbers. We assume that all measurements are non-negative, as is the case with most high-throughput experiments. (If this assumption does not hold, transformations can be applied to make data satisfy this assumption.)

Clipper has the following assumptions on the joint distribution of ***X***_1_, … , ***X***_*d*_, ***Y*** _1_, … , ***Y*** _*d*_. For *j* = 1, … , *d*, Clipper assumes that *X*_*j*1_, … , *X*_*jm*_ are identically distributed, so are *Y*_*j*1_, … , *Y_*jn*_*. Let 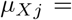 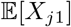 and 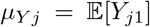 denote the expected measurement of feature *j* under the two conditions, respectively. Then conditioning on 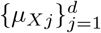 and 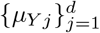,

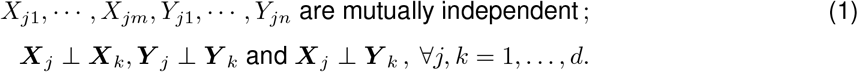

An enrichment analysis aims to identify interesting features with *μ*_*Xj*_ > *μ*_*Yj*_ (with ***X***_*j*_ and ***Y*** _*j*_ defined as the measurements under the experimental and background conditions, respectively), while a differential analysis aims to call interesting features with *μ*_*Xj*_ ≠ *μ*_*Yj*_. We define 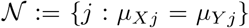 as the set of uninteresting features and denote 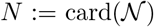. In both analyses, Clipper further assumes that an uninteresting feature *j* satisfies

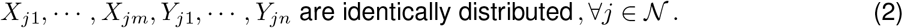

Clipper consists of two main steps: construction and thresholding of contrast scores. First, Clipper computes contrast scores, one per feature, as summary statistics that reflect the extent to which features are interesting. Second, Clipper establishes a contrast-score cutoff and calls as discoveries the features whose contrast scores exceed the cutoff.

To construct contrast scores, Clipper uses two summary statistics 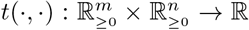 to extract data information regarding whether a feature is interesting or not:

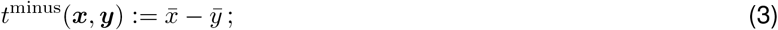

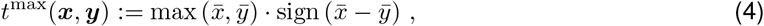

where 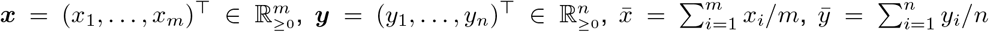, and sign(·) : ℝ → {−1, 0, 1} with sign(*x*) = 1 if *x* > 0, sign(*x*) = −1 if *x* < 0, and sign(*x*) = 0 otherwise.

Notably, other summary statistics can also be used to construct contrast scores. For example, an alternative summary statistic is the *t* statistic from the two-sample *t* test:

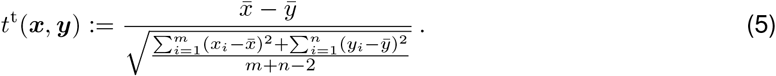

Then we introduce how Clipper works in three analysis tasks: the enrichment analysis with equal numbers of replicates under two conditions (*m* = *n*), the enrichment analysis with different numbers of replicates under two conditions (*m* ≠ *n*), and the differential analysis (when *m* + *n* > 2).

### Clipper: enrichment analysis with equal numbers of replicates (*m* = *n*)

Under the enrichment analysis, we assume that 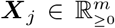 and 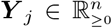 are the measurements of feature *j*, *j* = 1, … , *d*, under the experimental and background conditions with *m* and *n* replicates, respectively. We start with the simple case when *m* = *n*. Clipper defines a contrast score *C*_*j*_ of feature *j* in one of two ways:

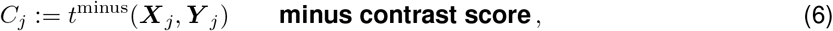

or

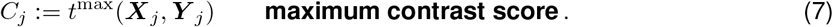

Fig. 6a shows a cartoon illustration of contrast scores when *m* = *n* = 1. Accordingly, a large positive value of *C*_*j*_ bears evidence that *μ*_*Xj*_ > *μ*_*Yj*_. Motivated by Barber and Candès [35], Clipper uses the following procedure to control the FDR under the target level *q* ∈ (0, 1).

**Figure 6.**
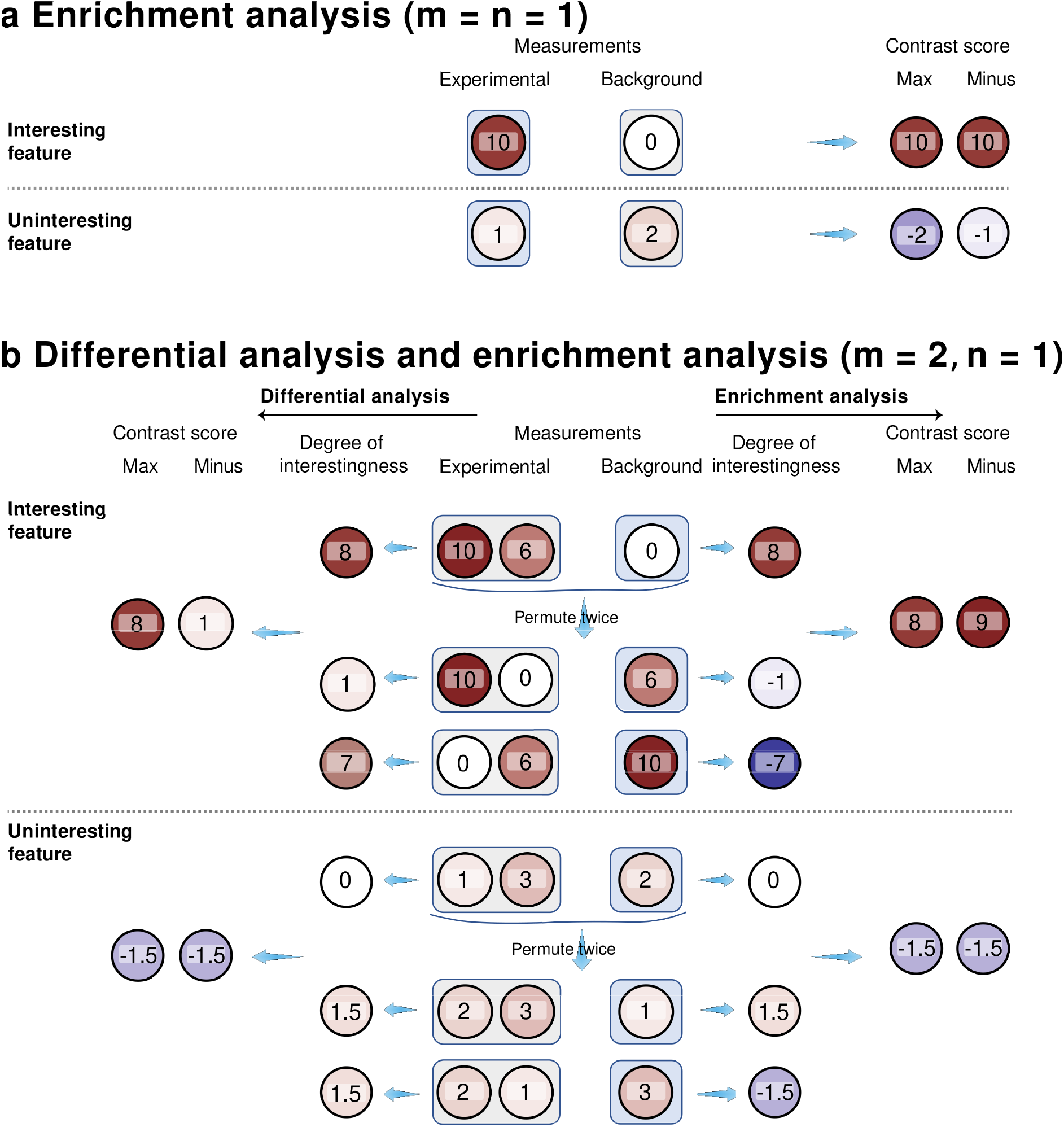
Illustration of the construction of contrast scores. **(a)** 1vs1 enrichment analysis; **(b)** 2vs1 differential analysis (left) or enrichment analysis (right). In each panel, an interesting feature (top) and an uninteresting feature (bottom) are plotted for contrast; both features have measurements under the experimental and background conditions. In (a), each feature’s measurements are summarized into a maximum (max) contrast score or a minus contrast score. In (b), each feature’s measurements are permuted across the two conditions, resulting in two sets of permuted measurements. Then for each feature, we calculate its degrees of interestingness—as the difference that equals the average of experimental measurements minus the average of background measurements (in enrichment analysis; right), or the absolute value of the difference (in differential analysis; left)—from its original measurements and permuted measurements, respectively. Finally, we summarize each feature’s degrees of interestingness into a maximum (max) contrast score or a minus contrast score.

#### 1 Definition (Barber-Candès (BC) procedure for thresholding contrast scores [35])

*Given contrast scores* 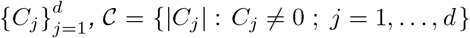 *is defined as the set of non-zero absolute values of C*_*j*_’s. *The BC procedure finds a contrast-score cutoff T*^BC^ *based on the target FDR threshold q* ∈ (0, 1) *as*

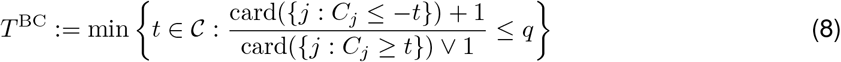

*and outputs* {*j* : *C*_*j*_ ≥ *T*^BC^} *as discoveries.*

### Clipper: enrichment analysis with any numbers of replicates *m* and *n*

When *m* ≠ *n*, Clipper constructs contrast scores via permutation of replicates across conditions. The idea is that, after permutation, every feature becomes uninteresting and can serve as its own negative control.

#### 2 Definition (Permutation)

*We define σ as permutation, i.e., a bijection from the set* {1, … , *m* + *n*} *onto itself, and we rewrite the data* ***X***_1_, … , ***X***_*d*_, ***Y*** _1_, … , ***Y*** _*d*_ *into a matrix* **W**:

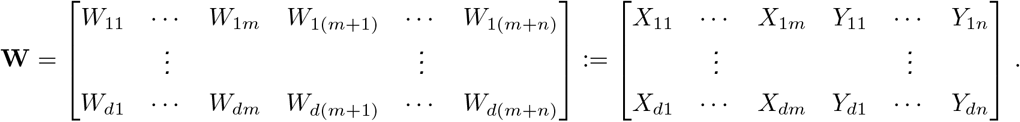

*We then apply σ to permute the columns of* **W** *and obtain*

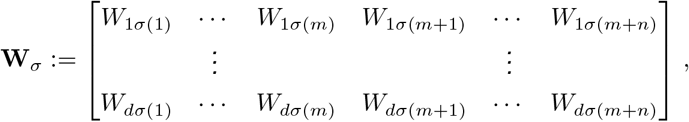

*from which we obtain the permuted measurements* 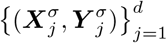, *where*

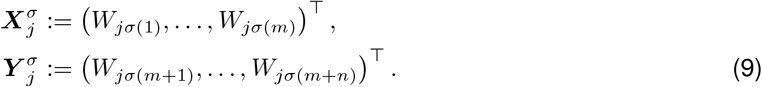

In the enrichment analysis, if two permutations *σ* and *σ*′ satisfy that

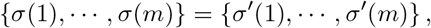

then we define *σ* and *σ*′ to be in one equivalence class. That is, permutations in the same equivalence class lead to the same division of *m* + *n* replicates (from the two conditions) into two groups with sizes *m* and *n*. In total, there are 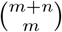 equivalence classes of permutations.

We define *σ*_0_ as the identity permutation such that *σ*_0_(*i*) = *i* for all *i* ∈ {1, … , *m* + *n*}. In addition, Clipper randomly samples *h* equivalence classes *σ*_1_, … , *σ*_*h*_ with equal probabilities without replacement from the other 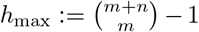 equivalence classes (after excluding the equivalence class containing *σ*_0_). Note that *h*_max_ is the maximum value *h* can take.

Clipper then obtains 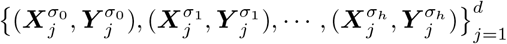, where 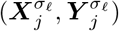 are the per-muted measurements based on *σ*_*l*_, = 0, 1, … , *h*. Then Clipper computes 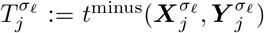 to indicate the degree of “interestingness” of feature *j* reflected by 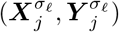. Note that Clipper chooses *t*^minus^ instead of *t*^max^ because empirical evidence shows that *t*^minus^ leads to better power. Sorting 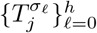 gives

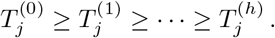

Then Clipper defines the contrast score of feature *j*, *j* = 1, … , *d*, in one of two ways:

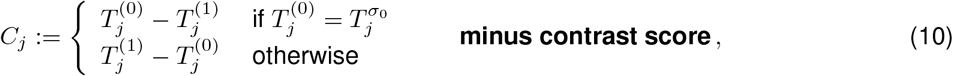

or

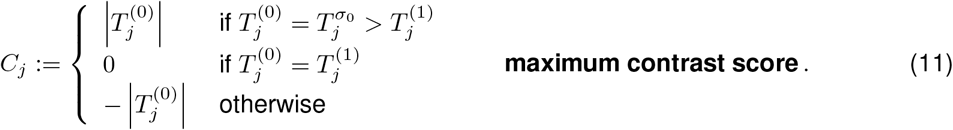

The intuition behind the contrast scores is that, if *C*_*j*_ < 0, then 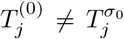, which means that at least one of 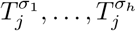 (calculated after random permutation) is greater than 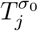 calculated from the original data (identity permutation), suggesting that feature *j* is likely an uninteresting feature in enrichment analysis. Fig. 6b (right) shows a cartoon illustration of contrast scores when *m* = 2 and *n* = 1. Motivated by Gimenez and Zou [63], we propose the following procedure for Clipper to control the FDR under the target level *q* ∈ (0, 1).

#### 3 Definition (Gimenez-Zou (GZ) procedure for thresholding contrast scores [63])

*Given h* ∈ {1, … , *h*_max_} *and contrast scores* 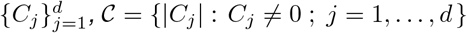 *is defined as the set of non-zero absolute values of *C*_*j*_’s. The GZ procedure finds a contrast-score cutoff T*^GZ^ *based on the target FDR threshold q* ∈ (0, 1) *as:*

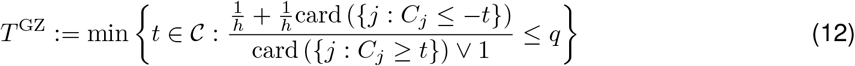

*and outputs* {*j* : *C*_*j*_ ≥ *T*^GZ^} *as discoveries.*

### Clipper: differential analysis with *m* + *n* > 2

For differential analysis, Clipper also uses permutation to construct contrast scores. When *m* ≠ *n*, the equivalence classes of permutations are defined the same as for the enrichment analysis with *m* ≠ *n*. When *m* = *n*, there is a slight change in the definition of equivalence classes of permutations: if *σ* and *σ*′ satisfy that

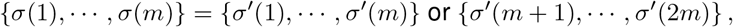

then we say that *σ* and *σ*′ are in one equivalence class. In total, there are 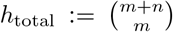 (when *m* ≠ *n*) or 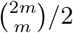 (when *m* = *n*) equivalence classes of permutations. Hence, to have more than one equivalence class, we cannot perform differential analysis with *m* = *n* = 1; in other words, the total number of replicates *m* + *n* must be at least 3.

Then Clipper randomly samples *σ*_1_, … , *σ*_*h*_ with equal probabilities without replacement from the *h*_max_ := *h*_total_ −1 equivalence classes that exclude the class containing *σ*_0_, i.e., the identity permutation. Note that *h*_max_ is the maximum value *h* can take. Next, Clipper computes 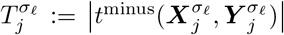, where 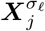 and 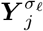 are the permuted data defined in (9), and it defines *C*_*j*_ as the contrast score of feature *j*, *j* = 1, … , *d*, in the same ways as in (10) or (11). Fig. 6b (left) shows a cartoon illustration of contrast scores when *m* = 2 and *n* = 1.

Same as in the enrichment analysis with *m* ≠ *n*, Clipper also uses the GZ procedure [63] to set a cutoff on contrast scores to control the FDR under the target level *q* ∈ (0, 1).

### Clipper: discussion on contrast scores

Granted, when we use permutations to construct contrast scores in the GZ procedure, we can convert contrast scores into permutation-based p-values (see Additional File 1: Section S1.1.2). However, when the numbers of replicates are small, the number of possible permutations is small, so permutation-based p-values would have a low resolution (e.g., when *m* = 2 and *n* = 1, the number of non-identity permutations is only 2). Hence, applying the BH procedure to the permutation-based p-values would result in almost no power. Although Yekutieli and Benjamini proposed another thresholding procedure for permutation-based p-values [67], it still requires the number of permutations to be large to obtain a reliable FDR control. Furthermore, if we apply the SeqStep+ procedure by Barber and Candés [35] to permutation-based p-values, it would be equivalent to our application of the GZ procedure to contrast scores (Additional File 1: Section S1.1.2).

For both differential and enrichment analyses, the two contrast scores (minus and maximum) can both control the FDR. Based on the power comparison results in Additional File 1: Section S3 and Additional File 1: Figs. S24–S28, Clipper has the following default choice of contrast score: for the enrichment analysis with *m* = *n*, Clipper uses the BC procedure with the minus contrast score; for the enrichment analysis with *m* ≠ n or the differential analysis, Clipper uses the GZ procedure with the maximum contrast score.

### Generic FDR control methods

In our simulation analysis, we compared Clipper against generic FDR control methods including p-value-based methods and local-fdr-based methods. Briefly, each p-value-based method is a combination of a p-value calculation approach and a p-value thresholding procedure. We use either the “paired” or “pooled” approach (see next paragraph) to calculate p-values of features and then threshold the p-values using the BH procedure (Additional File 1: Definition S1) or Storey’s qvalue procedure (Additional File 1: Definition S2) to make discoveries (Additional File 1: Section S1.1). As a result, we have four p-value-based methods: BH-pair, BH-pool, qvalue-pair, and qvalue-pool (Fig. 1b).

Regarding the existing p-value calculation approaches in bioinformatics tools, we categorize them as “paired” or “pooled.” The paired approach has been widely used to detect DEGs and protein-binding sites [1, 2, 4, 5]. It examines one feature at a time and compares the feature’s measurements between two conditions using a statistical test. In contrast, the pooled approach is popular in proteomics for identifying peptide sequences from MS data [68]. For every feature, it defines a test statistic and estimates a null distribution by pooling all features’ observed test statistic values under the background condition. Finally, it calculates a p-value for every feature based on the feature’s observed test statistic under the experimental condition and the null distribution.

In parallel to p-value-based methods, local-fdr-based methods estimate local fdrs of features and then threshold the local fdrs using the locfdr procedure (Additional File 1: Definition S5) to make discoveries. The estimation of local fdrs takes one of two approaches: (1) empirical null, which is estimated parametrically from the test statistic values that are likely drawn from the null distribution, and (2) swapping null, which is constructed by swapping measurements between experimental and background conditions. The resulting two local-fdr-based-methods are referred to as locfdr-emp and locfdr-swap (Figs. 1b and 2). Additional File 1: Section S1 provides a detailed explanation of these generic methods and how we implemented them in this work.

Specific to the p-value-based methods, for the paired approach, besides the ideal implementation that uses the correct model to calculate p-values (BH-pair-correct and qvalue-pair-correct), we also consider common mis-implementations. The first mis-implementations is misspecification of the distribution (BH-pair-mis and qvalue-pair-mis). An example is the detection of protein-binding sites from ChIP-seq data. A common assumption is that ChIP-seq read counts in a genomic region (i.e., a feature) follow the Poisson distribution [1, 2], which implies that the counts have the variance equal to the mean. However, if only two replicates are available, it is impossible to check whether this Poisson distribution is reasonably specified. The second mis-implementation is the misspecification of a two-sample test as a one-sample test (BH-pair-2as1 and qvalue-pair-2as1), which ignores the sampling randomness of replicates under one condition. This issue is implicit but widespread in bioinformatics methods [1, 69].

To summarize, we compared Clipper against the following implementations of generic FDR control methods:

- **BH-pool** or **qvalue-pool**: p-values calculated by the <u>pool</u>ed approach and thresholded by the <u>BH</u> or <u>qvalue</u> procedure.
- **BH-pair-correct** or **qvalue-pair-correct**: p-values calculated by the <u>pair</u>ed approach with the <u>correct</u> model specification and thresholded by the <u>BH</u> or <u>qvalue</u> procedure.
- **BH-pair-mis** or **qvalue-pair-mis**: p-values calculated by the <u>pair</u>ed approach with a <u>mis</u>specified model and thresholded by the <u>BH</u> or <u>qvalue</u> procedure.
- **BH-pair-2as1** or **qvalue-pair-2as1**: p-values calculated by the <u>pair</u>ed approach that misformulates a two-sample test as a one-sample test (<u>2as1</u>) and thresholded by the <u>BH</u> or <u>qvalue</u> procedure.
- **locfdr-emp**: local fdrs calculated by the <u>emp</u>irical null approach and thresholded by the <u>locfdr</u> procedure.
- **locfdr-swap**: local fdrs calculated by the <u>swap</u>ping approach and thresholded by the <u>locfdr</u> procedure.

### Software packages used in this study

- **p.adjust** `R` function (in `R` package `stats` v 4.0.2 with default arguments) [14]: used for BH-pool, BH-pair-correct, BH-pair-mis, and BH-pair-2as1.
- **qvalue** `R` package (v 2.20.0 with default arguments) [71]: used for qvalue-pool, qvalue-pair-correct, qvalue-pair-mis, and qvalue-pair-2as1.
- **locfdr** `R` package (v 1.1-8 with default arguments) [72]: used for locfdr-emp.
- **MACS2** software package (v 2.2.6 with default settings) [1]: available at https://github.com/macs3-project/MACS/releases/tag/v2.2.6.
- **ChIPulate** software package [45]: available at https://github.com/vishakad/chipulate.
- **HOMER** software package (findPeaks v 3.1.9.2 with default settings) [2]: available at https://www.bcgsc.ca/platform/bioinfo/software/findpeaks/releases/3.1.9.2/findpeaks3-1-9-2-tar.gz.
- **SEQUEST** in `Proteome Discoverer` (v 2.3.0.523 with default settings) [3]: commercial software by ThermoScientific.
- **edgeR** `R` package (v 3.30.0 with default arguments) [4]: available at https://www.bioconductor.org/packages/release/bioc/html/edgeR.html.
- **DESeq2** `R` package (v 1.28.1 with default arguments) [5]: available at http://bioconductor.org/packages/release/bioc/html/DESeq2.html.
- **limma** `R` package (v 3.44.3 with default arguments) [10]: available at https://www.bioconductor.org/packages/release/bioc/html/limma.html.
- **MAST** `R` package (v 1.14.0 with default arguments) [56]: available at https://www.bioconductor.org/packages/release/bioc/html/MAST.html.
- **monocle3** `R` package (v 0.2.3.0 with default arguments) [57]: available at https://github.com/cole-trapnell-lab/monocle3.
- **MultiHiCcompare** `R` package (v 1.6.0 with default arguments) [11]: available at https://bioconductor.org/packages/release/bioc/html/multiHiCcompare.html.
- **diffHic** `R` package (v 1.20.0 with default arguments) [13]: available at https://www.bioconductor.org/packages/release/bioc/html/diffHic.html.
- **FIND** `R` package (v 0.99 with default arguments) [12]: available at https://bitbucket.org/nadhir/find/src/master/.

## Availability of Data and Materials

- The **Clipper** `R` package is available at https://github.com/JSB-UCLA/Clipper/ [73].
- The code and processed data for reproducing the figures are available at https://zenodo.org/record/5115468 (DOI number 10.5281/zenodo.5115468) [74].
- A video introduction of Clipper is available at https://youtu.be/-GXyHiJMpLo.
- Real datasets:
  – The H3K4me3 ChIP-seq dataset with one experimental sample (GEO accession number GSM733708) and two control samples (GEO accession number GSM733742) from the cell line GM12878 is available at ftp://hgdownload.cse.ucsc.edu/goldenPath/hg19/encodeDCC/wgEncodeBroadHistone/, with the experimental sample `wgEncodeBroadHistoneGm12878H3k4me3StdAlnRep1.bam` and the two control samples `wgEncodeBroadHistoneGm12878ControlStdAlnRep1.bam` `wgEncodeBroadHistoneGm12878ControlStdAlnRep2.bam`. The processed dataset is available at https://zenodo.org/record/5115468.
  – The Pfu mass spectrometry proteomics data have been deposited to the ProteomeXchange Consortium via the PRIDE [70] partner repository with the dataset identifier PXD028558 (https://www.ebi.ac.uk/pride/archive/projects/PXD028558). The processed MS benchmark dataset is available as data/pfu/Archaea Sequest DECOYS 100 FDR.xlsx for the decoy PSMs and data/pfu/Archaea Sequest Targets 100 FDR.xlsx for the target PSMs at https://zenodo.org/record/5202768.
  – The human monocyte RNA-seq dataset (SRA accession number SRP082682) is available at https://www.ncbi.nlm.nih.gov/Traces/study/?acc=srp082682. The dataset includes 17 samples of classical monocytes and 17 samples of non-classical monocytes, and it is converted to a sample-by-gene count matrix by `R` package GenomicFeatures (v 1.40.1). The processed count matrix is available at https://zenodo.org/record/5115468.
  – The Hi-C dataset from the cell line GM12878 (GEO accession number GSE63525) is available at https://www.ncbi.nlm.nih.gov/geo/query/acc.cgi?acc=GSE63525. The count matrix is under the filename GSE63525 GM12878 primary intrachromosomal contact matrices.tar.gz, and the matrix corresponding to Chromosome 1 and bin width 1MB is used. The processed dataset is available at https://zenodo.org/record/5115468.
  – The PBMC single-cell RNA-seq dataset (GEO accession number GSE132044) is available at https://www.ncbi.nlm.nih.gov/geo/query/acc.cgi?acc=GSE132044. The processed dataset is available at https://zenodo.org/record/5115468.

## Acknowledgements

The authors would like to thank Dr. Yu-Cheng Yang for his suggestions on the figures and R package. The authors would also like to thank Mr. Nikos Ignatiadis, Dr. Lihua Lei, and Dr. Rina Barber for their insightful comments after we presented this work at the International Seminar on Selective Inference (https://www.selectiveinferenceseminar.com/past-talks). The authors also appreciate the comments and feedback from Mr. Tianyi Sun, Ms. Kexin Li, and other members of the Junction of Statistics and Biology at UCLA (http://jsb.ucla.edu).

## Author Contribution

J.J.L., X.G., and Y.E.C. developed the methodology. X.G. performed the simulation analysis and the DEG analysis on semi-synthetic single-cell RNA-seq data. X.G. did the peak calling analysis, advised by W.L.. Y.E.C. did the DEG analysis on semi-synthetic bulk RNA-seq data, the DIR analysis, and the peptide identification analysis. The real data used in peptide identification analysis was generated by M.M., K.W., and A.M., supported by L.D.W.. X.G. and Y.E.C. performed DEG analysis on real bulk RNA-seq data, the results of which were interpreted by X.G., D.S., and N.W.. X.G. and Y.E.C. developed the R package. Y.E.C. wrote the mathematical proof. X.G., Y.E.C., and J.J.L. wrote the manuscript. The authors read and approved the final manuscript.

## Ethics Approval

Not applicable.

## Additional Files

### Additional file 1

Supplementary materials. It includes a review of generic FDR control methods (Section S1 [75]), the detailed Clipper methodology (Section S2), the comparison of Clipper variant algorithms (Section S3), data generation and detailed implementation of the paired approach (a p-value calculation approach) in simulation studies (Section S4), bioinformatic methods with FDR control functionality (Section S5 [76, 77]), benchmark data generation in omics data applications (Section S6 [78–82]), implementation of Clipper in omics data applications (Section S7), proofs (Section S8), and Figures S1–S29.

### Additional file 2

Supplementary table. Biological functions of the DEGs (between classical and non-classical human monocytes) that were identified by Clipper only but missed by both edgeR and DESeq2 from the human monocyte RNA-seq dataset (SRA accession number SRP082682).

