## Additional file 1 for "Clipper: p-value-free FDR control on high-throughput data from two conditions"

### Supplementary Material

### S1 Review of generic FDR control methods

To facilitate our discussion, we introduce the notations for data. For feature $j = 1, \ldots, d$, we use $\boldsymbol{X}_j = (X_{j1}, \ldots, X_{jm})^\top \in \mathbb{R}^m_{\geq 0}$ and $\boldsymbol{Y}_j = (Y_{j1}, \ldots, Y_{jn})^\top \in \mathbb{R}^n_{\geq 0}$ to denote its measurements under the experimental and background conditions, respectively. We assume that $X_{j1}, \ldots, X_{jm}$ are identically distributed, so are $Y_{j1}, \ldots, Y_{jn}$. Let $\mu_{Xj} = \mathbb{E}[X_{j1}]$ and $\mu_{Yj} = \mathbb{E}[Y_{j1}]$ denote the expected measurement of feature $j$ under the two conditions, respectively. Then we denote by $\bar{X}_j$ the sample average of $X_{j1}, \cdots, X_{jm}$ and by $\bar{Y}_j$ the sample average of $Y_{j1}, \cdots, Y_{jn}$.

#### S1.1 P-value-based methods

Here we describe the details of p-value-based FDR control methods, including BH-pair, BH-pool, qvalue-pair, and qvalue-pool. Each of these four methods first computes p-values using either the pooled approach or the paired approach, and it then relies on the BH procedure [1] or Storey's qvalue procedure [2] for FDR control. In short, every p-value-based method is a combination of a p-value calculation approach and a p-value thresholding procedure. Below we introduce two p-value calculation approaches (paired and pooled) and two p-value thresholding procedures (BH and Storey's qvalue).

##### S1.1.1 P-value calculation approaches

**The paired approach.** The paired approach examines one feature at a time and compares its measurements between two conditions. Besides the ideal implementation, i.e., the *correct paired approach* that uses the correct model to calculate p-values, we also include commonly-used flawed implementations that either misspecify the distribution, i.e., the *misspecified paired approach*, or misformulate the two-sample test as a one-sample test, i.e., the *2as1 paired approach*.

Here we use the negative binomial distribution as an example to demonstrate the ideas of the correct, misspecified, and 2as1 paried approaches. Suppose that for each feature $j$, its measurements under each condition follow a negative binomial distribution, and the two distributions under the two conditions have the same dispersion; that is, $X_{j1}, \cdots, X_{jm} \overset{\text{i.i.d.}}{\sim} \text{NB}(\mu_{Xj}, \theta_j)$ ; $Y_{j1}, \cdots, Y_{jn} \overset{\text{i.i.d.}}{\sim} \text{NB}(\mu_{Yj}, \theta_j)$, where $\theta_j$ is the dispersion parameter such that the variance $\text{Var}(X_{ji}) = \mu_{Xj} + \theta_j \mu^2_{Xj}$.

- The correct paired approach assumes that the two negative binomial distributions have the same dispersion parameter $\theta_j$, and it uses the two-sample test for the null hypothesis $H_0 : \mu_{Xj} = \mu_{Yj}$ against the alternative hypothesis $H_1 : \mu_{Xj} > \mu_{Yj}$ (enrichment analysis) or $H_1 : \mu_{Xj} \neq \mu_{Yj}$ (differential analysis).
- The misspecified paired approach misspecifies the negative binomial distribution as Poisson, and it uses the two-sample test for the null hypothesis $H_0 : \mu_{Xj} = \mu_{Yj}$ against the alternative hypothesis $H_1 : \mu_{Xj} > \mu_{Yj}$ (enrichment analysis) or $H_1 : \mu_{Xj} \neq \mu_{Yj}$ (differential analysis).
- The 2as1 paired approach bluntly assumes $\mu_{Yj} = \bar{Y}_j$, and it performs the one-sample test based on $X_{j1}, \ldots, X_{jm}$ for the null hypotheses $H_0 : \mu_{Xj} = \bar{Y}_j$ against the alternative hypothesis $H_1 : \mu_{Xj} > \bar{Y}_j$ (enrichment analysis) or $H_1 : \mu_{Xj} \neq \bar{Y}_j$ (differential analysis).

**The pooled approach.** The pooled approach pools all features' average measurements under the background condition $\{\bar{Y}_j\}_{j=1}^d$ to form a null distribution, and it calculates a p-value for each feature $j$

by comparing $\bar{X}_j$ to the null distribution. Specifically, in enrichment analysis, the p-value of feature $j$ is computed as:

$$p_j = \frac{\mathrm{card}\left(\{k : \bar{Y}_k \geq \bar{X}_j\}\right)}{d} .$$

In differential analysis, the p-value of feature $j$ is computed as:

$$p_j = 2 \cdot \min\left(\frac{\mathrm{card}\left(\{k : \bar{Y}_k \geq \bar{X}_j\}\right)}{d}, \frac{\mathrm{card}\left(\{k : \bar{Y}_k \leq \bar{X}_j\}\right)}{d}\right) .$$

##### S1.1.2 P-value thresholding procedures for FDR control

**Definition S1 (BH procedure for thresholding p-values [1])** *The features' p-values $p_1, \ldots, p_d$ are sorted in an ascending order $p_{(1)} \leq p_{(2)} \leq \ldots \leq p_{(d)}$. Given the target FDR threshold $q$, the Benjamini–Hochberg (BH) procedure finds a p-value cutoff $T^{\mathrm{BH}}$ as*

$$T^{\mathrm{BH}} := p_{(k)}, \text{ where } k = \max\left\{j = 1, \ldots, d : p_{(j)} \leq \frac{j}{d} q\right\} . \tag{S1}$$

*Then BH outputs $\{j : p_j \leq T^{\mathrm{BH}}\}$ as discoveries.*

**Definition S2 (Storey's qvalue procedure for thresholding p-values [2])** *The features' p-values $p_1, \ldots, p_d$ are sorted in an ascending order $p_{(1)} \leq p_{(2)} \leq \ldots \leq p_{(d)}$. Let $\hat{\pi}_0$ denote an estimate of the probability $P($the $i$-th feature is uninteresting$)$ (see Storey [2] for details). Storey's qvalue procedure defines the q-value for $p_{(d)}$ as*

$$\hat{q}(p_{(d)}) := \frac{\hat{\pi}_0 \cdot d \cdot p_{(d)}}{\mathrm{card}\left(\{k : p_k \leq p_{(d)}\}\right)} = \hat{\pi}_0 \cdot p_{(d)} .$$

*Then for $j = d-1, d-2, \ldots, 1$, the q-value for $p_{(j)}$ is defined as:*

$$\hat{q}(p_{(j)}) := \min\left(\hat{q}(p_{(j+1)}), \frac{\hat{\pi}_0 \cdot d \cdot p_{(j)}}{\mathrm{card}\left(\{k : p_k \leq p_{(j)}\}\right)}\right) .$$

*Then Storey's qvalue procedure outputs $\{j : \hat{q}(p_j) \leq q\}$ as discoveries.*

We use function `qvalue` from `R` package `qvalue` (v 2.20.0; with default estimate $\hat{\pi}_0$) to calculate q-values.

**Definition S3 (SeqStep+ procedure for thresholding p-values [3])** *Define $H_0^j$ as the null hypothesis for feature $j$ and $p_j$ as the p-value for $H_0^j$, $j = 1, \ldots, d$. Order the null hypotheses $H_0^1, \ldots, H_0^d$ from the most to the least promising (here more promising means more likely to be interesting) and denote the resulting null hypotheses and p-values as $H_0^{(1)}, \ldots, H_0^{(d)}$ and $p_{(1)}, \ldots, p_{(d)}$. Given any target FDR threshold $q$, a pre-specified constant $s \in (0,1)$, and a subset $\mathcal{K} \subseteq \{1, \ldots, d\}$, the SeqStep+ procedure finds a cutoff $\hat{j}$ as*

$$\hat{j} := \max\left\{j \in \mathcal{K} : \frac{1 + \mathrm{card}\left(\{k \in \mathcal{K}, k \leq j : p_{(k)} > s)\}\right.}{\mathrm{card}\left(\{k \in \mathcal{K}, k \leq j : p_{(k)} \leq s)\} \vee 1\right.} \leq \frac{1-s}{s} q\right\} \tag{S2}$$

*Then SeqStep+ rejects $\left\{H_0^{(j)} : p_{(j)} \leq s,\ j \leq \hat{j},\ j \in \mathcal{K}\right\}$. If the orders of the null hypotheses are independent of the p-values, the SeqStep+ procedure ensures FDR control.*

The GZ procedure (Definition 3) used in Clipper is a special case of the SeqStep+ procedure with $s = 1/(h+1)$. Recall that given the number of non-identical permutations $h \in \{1, \cdots, h_{\max}\}$ and contrast scores $\{C_j\}_{j=1}^d$, the GZ procedure sorts $\{|C_j|\}_{j=1}^d$ in a decreasing order:

$$|C_{(1)}| \geq |C_{(2)}| \geq \cdots \geq |C_{(d)}| \,. \tag{S3}$$

To see the connection between the GZ procedure and SeqStep+, we consider the null hypothesis for the $j$-th ordered feature, $j = 1, \ldots, d$, as $H_0^{(j)} : \mu_{X(j)} = \mu_{Y(j)}$ and define the corresponding p-value $p_{(j)} := \frac{r(T_{(j)}^{\sigma_0})}{h+1}$, where $r(T_{(j)}^{\sigma_0})$ is the rank of $T_{(j)}^{\sigma_0}$ in $\{T_{(j)}^{\sigma_0}, \cdots, T_{(j)}^{\sigma_h}\}$ in a descending order. We also define $\mathcal{K} := \{j = 1, \ldots, d : C_j \neq 0\}$ as the subset of features with non-zero $C_j$'s. Finally, we input the p-values, null hypothesis orders in (S3), $s = 1/(h+1)$, $q$ and $\mathcal{K}$ into the SeqStep+ procedure, and we obtain the GZ procedure.

The BC procedure (Definition 1) is a further special case with $h = 1$, $p_{(j)} := \left(\mathbb{1}(C_{(j)} > 0) + 1\right)/2$, and $\mathcal{K} := \{j = 1, \ldots, d : C_j \neq 0\}$.

##### S1.2 Local-fdr-based methods

The FDR is statistical criterion that ensures the reliability of discoveries as a whole. In contrast, the local fdr focuses on the reliability of each discovery. The definition of the local fdr relies on some pre-computed summary statistics $z_j$ for feature $j$, $j = 1, \ldots, d$. In the calculation of local fdr, $\{z_1, \ldots, z_d\}$ are assumed to be realizations of an abstract random variable $Z$ that represents any feature. Let $p_0$ or $p_1$ denote the prior probability that any feature is uninteresting or interesting, with $p_0 + p_1 = 1$. Let $f_0(z) := \mathbb{P}(Z = z \,|\, \text{uninteresting})$ or $f_1(z) := \mathbb{P}(Z = z \,|\, \text{interesting})$ denote the conditional probability density of $Z$ at $z$ given that $Z$ represents an uninteresting or interesting feature. Thus by Bayes' theorem, the posterior probability of any feature being uninteresting given its summary statistic $Z = z$ is

$$\mathbb{P}(\text{uninteresting} \mid Z = z) = p_0 f_0(z)/f(z) \,, \tag{S4}$$

where $f(z) := p_0 f_0(z) + p_1 f_1(z)$ is the marginal probability density of $Z$. Accordingly, the local fdr of feature $j$ is defined as follows.

**Definition S4 (Local fdr [4])** *Given notations defined above, the local fdr of feature $j$ is defined as*

$$\textit{local-fdr}_j := f_0(z_j)/f(z_j) \,.$$

*Because $p_0 \leq 1$, local-fdr$_j$ is an upper bound of the posterior probability of feature $j$ being uninteresting given its summary statistic $z_j$, defined in (S4).*

Note that another definition of the local fdr is the posterior probability $\mathbb{P}(\text{uninteresting} \mid z)$ in (S4) [5]. Although this other definition is more reasonable, we do not use it but choose Definition S4 because the estimation of $p_0$ is ususally difficult. Another reason is that uninteresting features are the dominant majority in high-throughput biological data, so $p_0$ is often close to $1$.

We define local-fdr-based methods as a type of FDR control methods by thresholding local fdrs of features under the target FDR threshold $q$. Although the local fdr is different from FDR, it has been shown that thresholding the local fdrs at $q$ will approximately control the FDR under $q$ [4]. This makes local-fdr-based methods competitors against Clipper and p-value-based methods.

Every local-fdr-based method is a combination of a local fdr calculation approach and a local fdr thresholding procedure. Below we introduce two local fdr calculation approaches (empirical null and swapping) and one local fdr thresholding procedure. After the combination, we have two local-fdr-based methods: locfdr-emp and locfdr-swap.

##### S1.2.1 Local fdr calculation approaches

With $z_1, \ldots, z_d$, the calculation of local fdr defined in Definition S4 requires the estimation of $f_0$ and $f$, two probability densities. $f$ is estimated by nonparametric density estimation, and $f_0$ is estimated by either the empirical null approach [4] or the swapping approach, which shuffles replicates between conditions [5]. With the estimated $\hat{f}$ and $\hat{f}_0$, the estimated local fdr of feature $j$ is

$$\widehat{\text{local-fdr}}_j := \hat{f}_0(z_j)/\hat{f}(z_j)\,. \tag{S5}$$

**The empirical null approach.** This approach assumes a parametric distribution, typically the Gaussian distribution, to estimate $f_0$. Then with the density estimate $\hat{f}$, the local fdr is estimated for each feature $j$. The implementation of this approach depends on the numbers of replicates.

- In 1vs1 enrichment and differential analyses, we define $z_j$ as

$$z_j := \frac{D_j}{\sqrt{\frac{1}{d}\sum_{j=1}^{d}\left(D_j - \bar{D}\right)^2}}\,,$$

  where $D_j = X_{j1} - Y_{j1}$ and $\bar{D} = \sum_{j=1}^{d} D_j/d$.

- In 2vs1 enrichment and differential analyses, we define $z_j$ as

$$z_j := \frac{\bar{X}_j - Y_{j1}}{\sqrt{\frac{s_{Xj}^2}{2}}}\,,$$

  where $s_{Xj}^2 = \sum_{i=1}^{2}(X_{ji} - \bar{X}_j)^2$.

- In $m$vs$n$ enrichment and differential analyses with $m, n \geq 2$, we define $z_j$ as the two-sample $t$-statistic with unequal variances:

$$z_j := \frac{\bar{X}_j - \bar{Y}_j}{\sqrt{\frac{s_{Xj}^2}{m} + \frac{s_{Yj}^2}{n}}}\,,$$

  where $s_{Xj}^2 = \frac{1}{m-1}\sum_{i=1}^{m}(X_{ji} - \bar{X}_j)^2$ and $s_{Yj}^2 = \frac{1}{n-1}\sum_{i=1}^{n}(Y_{ji} - \bar{Y}_j)^2$ are the sample variances of feature $j$ under the experimental and background conditions.

Then $\{\widehat{\text{locfdr}}_j\}_{j=1}^d$ are estimated from $\{z_j\}_{j=1}^d$ by function `locfdr` in `R` package `locfdr` (v 1.1-8; with default arguments).

**The swapping approach.** This approach swaps $\lceil m/2 \rceil$ replicates under the experimental condition with $\lceil n/2 \rceil$ replicates under the background condition. Then it calculates the summary statistic for each feature on the swapped data, obtaining $z_1', \ldots, z_d'$. Finally, it estimates $f_0$ and $f$ by applying kernel density estimation to $z_1', \ldots, z_d'$ and $z_1, \ldots, z_d$, respectively (by function `kde` in `R` package `ks`). With $\hat{f}_0$ and $\hat{f}$, $\{\widehat{\text{locfdr}}_j\}_{j=1}^d$ are calculated by Definition S4.

The implementation of this approach depends on the numbers of replicates. Below are three special cases included in this work.

- In 1vs1 enrichment and differential analyses, the swapping approach is inapplicable because interesting features would not become uninteresting after the swapping.

- In 2vs1 enrichment and differential analyses, we define $z_j$ and $z'_j$ as

$$z_j = \frac{X_{j1} + X_{j2}}{2} - Y_{j1}\,,$$
$$z'_j = \frac{X_{j1} + Y_{j1}}{2} - X_{j2}\,.$$

- In $3$vs$3$ enrichment and differential analyses with, we define $z_j$ and $z'_j$ as

$$z_j = \frac{X_{j1} + X_{j2}}{2} - \frac{Y_{j1} + Y_{j2}}{2}\,,$$
$$z'_j = \frac{X_{j1} + Y_{j1}}{2} - \frac{X_{j2} + Y_{j2}}{2}\,.$$

Then we apply kernel density estimation to $\{z_j\}_{j=1}^d$ and $\{z'_j\}_{j=1}^d$ to obtain $\hat{f}$ and $\hat{f}_0$, respectively. By (S5), we calculate $\{\widehat{\text{locfdr}}_j\}_{j=1}^d$.

###### S1.2.2 The local fdr thresholding procedure

**Definition S5 (locfdr procedure)** *Given the local fdr estimates* $\{\widehat{\textit{local-fdr}}_j\}_{j=1}^d$ *and the target FDR threshold* $q$*, the locfdr procedure outputs* $\{j = 1, \ldots, d : \widehat{\textit{local-fdr}}_j \leq q\}$ *as discoveries.*

#### S2 The Clipper methodology

Clipper is a flexible framework that reliably controls the FDR without using p-values in high-throughput data analysis with two conditions. Clipper has two functionalities: (I) enrichment analysis, which identifies the "interesting" features that have higher expected measurements (i.e., true signals) under the experimental condition than the background, a.k.a. negative control condition (if the goal is to identify the interesting features with smaller expected measurements under the experimental condition, enrichment analysis can be applied after the values are negated); (II) differential analysis, which identifies the interesting features that have different expected measurements between the two conditions. For both functionalities, uninteresting features are defined as those that have equal expected measurements under the two conditions.

Clipper only relies on two fundamental statistical assumptions of biological data analysis: (1) measurement errors (i.e., differences between measurements and their expectations, with the expectations including biological signals and batch effects) are independent across all features and experiments; (2) every uninteresting feature has measurement errors identically distributed across all experiments. These two assumptions are used in almost all bioinformatics tools and commonly referred to as the "measurement model" in statistical genomics [6].

In the following subsections, we will first introduce notations and assumptions used in Clipper. Then we will detail how Clipper works and discuss its theoretical guarantee in three analysis tasks: the enrichment analysis with equal numbers of replicates under two conditions ($m = n$), the enrichment analysis with different numbers of replicates under two conditions ($m \neq n$), and the differential analysis (when $m + n > 2$)..

##### S2.1 Notations and assumptions

To facilitate our discussion, we first introduce the following mathematical notations. For two random vectors $\boldsymbol{X} = (X_1, \ldots, X_m)^\top$ and $\boldsymbol{Y} = (Y_1, \ldots, Y_n)^\top$, or two sets of random variables $\mathcal{X} = \{X_1, \ldots, X_m\}$

and $\mathcal{Y} = \{Y_1, \ldots, Y_n\}$, we write $\boldsymbol{X} \perp \boldsymbol{Y}$ or $\mathcal{X} \perp \mathcal{Y}$ if $X_i$ is independent of $Y_j$ for all $i = 1, \ldots, m$ and $j = 1, \ldots, n$. To avoid confusion, we use $\mathrm{card}(A)$ to denote the cardinality of a set $A$ and $|c|$ to denote the absolute value of a scalar $c$. We define $a \vee b := \max(a, b)$.

Clipper only requires two inputs: the target FDR threshold $q \in (0, 1)$ and the input data. Regarding the input data, we use $d$ to denote the number of features with measurements under two conditions, and we use $m$ and $n$ to denote the numbers of replicates under the two conditions. For each feature $j = 1, \ldots, d$, we use $\boldsymbol{X}_j = (X_{j1}, \ldots, X_{jm})^\top \in \mathbb{R}^m_{\geq 0}$ and $\boldsymbol{Y}_j = (Y_{j1}, \ldots, Y_{jn})^\top \in \mathbb{R}^n_{\geq 0}$ to denote its measurements under the two conditions, where $\mathbb{R}_{\geq 0}$ denotes the set of non-negative real numbers. We assume that all measurements are non-negative, as in the case of most high-throughput experiments. (If this assumption does not hold, transformations can be applied to make data satisfy this assumption.)

Clipper has the following assumptions on the joint distribution of $\boldsymbol{X}_1, \ldots, \boldsymbol{X}_d, \boldsymbol{Y}_1, \ldots, \boldsymbol{Y}_d$. For $j = 1, \ldots, d$, Clipper assumes that $X_{j1}, \ldots, X_{jm}$ are identically distributed, so are $Y_{j1}, \ldots, Y_{jn}$. Let $\mu_{Xj} = \mathbb{E}[X_{j1}]$ and $\mu_{Yj} = \mathbb{E}[Y_{j1}]$ denote the expected measurement of feature $j$ under the two conditions, respectively. Then conditioning on $\{\mu_{Xj}\}_{j=1}^d$ and $\{\mu_{Yj}\}_{j=1}^d$,

$$\begin{aligned} X_{j1}, \cdots, X_{jm}, Y_{j1}, \cdots, Y_{jn} \text{ are mutually independent}; \\ \boldsymbol{X}_j \perp \boldsymbol{X}_k, \boldsymbol{Y}_j \perp \boldsymbol{Y}_k \text{ and } \boldsymbol{X}_j \perp \boldsymbol{Y}_k\,, \ \forall j, k = 1, \ldots, d. \end{aligned} \tag{S6}$$

An enrichment analysis aims to identify interesting features with $\mu_{Xj} > \mu_{Yj}$ (with $\boldsymbol{X}_j$ and $\boldsymbol{Y}_j$ defined as the measurements under the experimental and background conditions, respectively), while a differential analysis aims to call interesting features with $\mu_{Xj} \neq \mu_{Yj}$. We define $\mathcal{N} := \{j : \mu_{Xj} = \mu_{Yj}\}$ as the set of uninteresting features and denote $N := \mathrm{card}(\mathcal{N})$. In both analyses, Clipper further assumes that an uninteresting feature $j$ satisfies

$$X_{j1}, \cdots, X_{jm}, Y_{j1}, \cdots, Y_{jn} \text{ are identically distributed}\,, \forall j \in \mathcal{N}\,. \tag{S7}$$

Clipper consists of two main steps: construction and thresholding of contrast scores. First, Clipper computes contrast scores, one per feature, as summary statistics that reflect the extent to which features are interesting. Second, Clipper establishes a contrast-score cutoff and calls as discoveries the features whose contrast scores exceed the cutoff.

To construct contrast scores, Clipper uses two summary statistics $t(\cdot, \cdot) : \mathbb{R}^m_{\geq 0} \times \mathbb{R}^n_{\geq 0} \to \mathbb{R}$ to extract data information regarding whether a feature is interesting or not:

$$t^{\text{minus}}(\boldsymbol{x}, \boldsymbol{y}) := \bar{x} - \bar{y}\,; \tag{S8}$$

$$t^{\max}(\boldsymbol{x}, \boldsymbol{y}) := \max(\bar{x}, \bar{y}) \cdot \mathrm{sign}(\bar{x} - \bar{y})\,, \tag{S9}$$

where $\boldsymbol{x} = (x_1, \ldots, x_m)^\top \in \mathbb{R}^m_{\geq 0}$, $\boldsymbol{y} = (y_1, \ldots, y_n)^\top \in \mathbb{R}^n_{\geq 0}$, $\bar{x} = \sum_{i=1}^m x_i/m$, $\bar{y} = \sum_{i=1}^n y_i/n$, and $\mathrm{sign}(\cdot) : \mathbb{R} \to \{-1, 0, 1\}$ with $\mathrm{sign}(x) = 1$ if $x > 0$, $\mathrm{sign}(x) = -1$ if $x < 0$, and $\mathrm{sign}(x) = 0$ otherwise.

Notably, other summary statistics can also be used to construct contrast scores. For example, an alternative summary statistic is the $t$ statistic from the two-sample $t$ test:

$$t^{\mathrm{t}}(\boldsymbol{x}, \boldsymbol{y}) := \frac{\bar{x} - \bar{y}}{\sqrt{\frac{\sum_{i=1}^m (x_i - \bar{x})^2 + \sum_{i=1}^n (y_i - \bar{y})^2}{m+n-2}}}\,.$$

#### S2.2 Enrichment analysis with equal numbers of replicates ($m = n$)

Under the enrichment analysis, we assume that $\boldsymbol{X}_j \in \mathbb{R}^m_{\geq 0}$ and $\boldsymbol{Y}_j \in \mathbb{R}^n_{\geq 0}$ are the measurements of feature $j$, $j = 1, \ldots, d$, under the experimental and background conditions with $m$ and $n$ replicates, respectively. We start with the simple case when $m = n$. Clipper defines a contrast score $C_j$ of feature $j$ in one of two ways:

$$C_j := t^{\text{minus}}(\boldsymbol{X}_j, \boldsymbol{Y}_j) \qquad \textbf{minus contrast score}\,, \tag{S10}$$

or

$$C_j := t^{\max}(\boldsymbol{X}_j, \boldsymbol{Y}_j) \qquad \textbf{maximum contrast score}\,. \tag{S11}$$

Accordingly, a large positive value of $C_j$ bears evidence that $\mu_{Xj} > \mu_{Yj}$. Motivated by Barber and Candès [3] and Arias-Castro and Chen [7], Clipper proposes the following BC procedure to control the FDR under the target level $q \in (0, 1)$.

**Definition S6 (Barber-Candès (BC) procedure for thresholding contrast scores [3])** *Given contrast scores* $\{C_j\}_{j=1}^d$, $\mathcal{C} = \{|C_j| : C_j \neq 0\,;\ j = 1, \ldots, d\}$ *is defined as the set of non-zero absolute values of* $C_j$*'s. The BC procedure finds a contrast-score cutoff* $T^{\mathrm{BC}}$ *based on the target FDR threshold* $q \in (0, 1)$ *as*

$$T^{\mathrm{BC}} := \min\left\{t \in \mathcal{C} : \frac{\operatorname{card}(\{j : C_j \leq -t\}) + 1}{\operatorname{card}(\{j : C_j \geq t\}) \vee 1} \leq q\right\} \tag{S12}$$

*and outputs* $\{j : C_j \geq T^{\mathrm{BC}}\}$ *as discoveries.*

**Theorem 1** *Suppose that the input data satisfy the Clipper assumptions* (S6)–(S7) *and* $m = n$*. Then for any* $q \in (0, 1)$ *and either definition of constrast scores in* (S10) *or* (S11)*, the contrast-score cutoff* $T^{\mathrm{BC}}$ *found by the BC procedure guarantees that the discoveries have the FDR under* $q$*:*

$$\mathrm{FDR} = \mathbb{E}\left[\frac{\operatorname{card}\left(\{j \in \mathcal{N} : C_j \geq T^{\mathrm{BC}}\}\right)}{\operatorname{card}(\{j : C_j \geq T^{\mathrm{BC}}\}) \vee 1}\right] \leq q\,,$$

*where* $\mathcal{N} = \{j : \mu_{Xj} = \mu_{Yj}\}$ *denotes the set of uninteresting features.*

The proof of Theorem 1 (Supp. Section S8) requires two key ingredients: Lemma 1, which states important properties of contrast scores, and Lemma 2 from [8], which states a property of a Bernoulli process with independent but not necessarily identically distributed random variables. The cutoff $T^{\mathrm{BC}}$ can be viewed as a stopping time of a Bernoulli process.

**Lemma 1** *Suppose that the input data that satisfy the Clipper assumptions* (S6)–(S7) *and* $m = n$*, and that Clipper constructs contrast scores* $\{C_j\}_{j=1}^d$ *based on* (S10) *or* (S11)*. Denote* $S_j = \operatorname{sign}(C_j) \in \{-1, 0, 1\}$*. Then* $\{S_j\}_{j=1}^d$ *satisfy the following properties:*

*(a)* $S_1, \ldots, S_d$ *are mutually independent ;*

*(b)* $\mathbb{P}(S_j = 1) = \mathbb{P}(S_j = -1)$ *for all* $j \in \mathcal{N}$*;*

*(c)* $\{S_j\}_{j \in \mathcal{N}} \perp \mathcal{C}$*.*

Notably, Lemma 1(a) can be relaxed as $\mathbb{P}(S_j = 1) \leq \mathbb{P}(S_j = -1)$ for all $j \in \mathcal{N}$. Then Lemma 2 still holds, and so does Theorem 1, making Clipper still have theoretical FDR control.

**Lemma 2** *Suppose that* $Z_1, \ldots, Z_d$ *are independent with* $Z_j \sim \mathrm{Bernoulli}(\rho_j)$*, and* $\min_j \rho_j \geq \rho > 0$*. Let* $J$ *be a stopping time in reverse time with respect to the filtration* $\{\mathcal{F}_j\}$*, where*

$$\mathcal{F}_j = \sigma\left(\{(Z_1 + \cdots + Z_j),\, Z_{j+1}, \cdots, Z_d\}\right)\,, \tag{S13}$$

*with $\sigma(\cdot)$ denoting a $\sigma$-algebra. Then*

$$\mathbb{E}\left[\frac{1+J}{1+Z_1+\cdots+Z_J}\right] \leq \rho^{-1}.$$

Here we give a brief intuition about how Lemma 2 bridges Lemma 1 and Theorem 1 for FDR control. First we note that the false discovery proportion (FDP), whose expectation is the FDR, satisfies

$$\begin{aligned}
\mathrm{FDP} &:= \frac{\operatorname{card}\left(\{j \in \mathcal{N} : C_j \geq T^{\mathrm{BC}}\}\right)}{\operatorname{card}\left(\{j : C_j \geq T^{\mathrm{BC}}\}\right) \vee 1} && \text{(S14)}\\
&= \frac{\operatorname{card}\left(\{j \in \mathcal{N} : C_j \geq T^{\mathrm{BC}}\}\right)}{\operatorname{card}\left(\{j \in \mathcal{N} : C_j \leq -T^{\mathrm{BC}}\}\right)+1} \cdot \frac{\operatorname{card}\left(\{j \in \mathcal{N} : C_j \leq -T^{\mathrm{BC}}\}\right)+1}{\operatorname{card}\left(\{j : C_j \geq T^{\mathrm{BC}}\}\right) \vee 1} && \text{(S15)}\\
&\leq \frac{\operatorname{card}\left(\{j \in \mathcal{N} : C_j \geq T^{\mathrm{BC}}\}\right)}{\operatorname{card}\left(\{j \in \mathcal{N} : C_j \leq -T^{\mathrm{BC}}\}\right)+1} \cdot \frac{\operatorname{card}\left(\{j : C_j \leq -T^{\mathrm{BC}}\}\right)+1}{\operatorname{card}\left(\{j : C_j \geq T^{\mathrm{BC}}\}\right) \vee 1} && \text{(S16)}\\
&\leq \frac{\operatorname{card}\left(\{j \in \mathcal{N} : C_j \geq T^{\mathrm{BC}}\}\right)}{\operatorname{card}\left(\{j \in \mathcal{N} : C_j \leq -T^{\mathrm{BC}}\}\right)+1} \cdot q\,, && \text{(S17)}
\end{aligned}$$

where the last inequality follows from the definition of $T^{\mathrm{BC}}$ (S12).

By its definition, if $T^{\mathrm{BC}}$ exists, it is positive. This implies that Clipper would never call the features with $C_j = 0$ as discoveries. Here we sketch the idea of proving Theorem 1 by considering a simplified case where $\mathcal{C}$ is fixed instead of being random; that is, we assume the features with non-zero contrast scores to be known. Then, without loss of generality, we assume $\mathcal{C} = \{1, \ldots, d\}$. Then we order the absolute values of uninteresting features' contrast scores, i.e., elements in $\{|C_j| : j \in \mathcal{N}\}$, from the largest to the smallest, denoted by $|C_{(1)}| \geq |C_{(2)}| \geq \cdots \geq |C_{(N)}|$. Let $J = \sum_{j\in\mathcal{N}} \mathbb{1}\left(|C_j| \geq T^{\mathrm{BC}}\right)$, the number of uninteresting features whose contrast scores have absolute values no less than $T^{\mathrm{BC}}$. When $J > 0$, $|C_{(1)}| \geq \cdots \geq |C_{(J)}| \geq T^{\mathrm{BC}}$. Define $Z_k = \mathbb{1}\left(C_{(k)} < 0\right)$, $k = 1, \ldots, N$. Then for each order $k$, the following holds

$$\begin{aligned}
C_{(k)} \geq T^{\mathrm{BC}} &\iff \left|C_{(k)}\right| \geq T^{\mathrm{BC}} \text{ and } C_{(k)} > 0 \iff k \leq J \text{ and } Z_k = 0\,;\\
C_{(k)} \leq -T^{\mathrm{BC}} &\iff \left|C_{(k)}\right| \geq T^{\mathrm{BC}} \text{ and } C_{(k)} < 0 \iff k \leq J \text{ and } Z_k = 1\,.
\end{aligned}$$

Then the upper bound of FDP becomes

$$\begin{aligned}
\frac{\operatorname{card}(\{j \in \mathcal{N} : C_j \geq T^{\mathrm{BC}}\})}{\operatorname{card}(\{j \in \mathcal{N} : C_j \leq -T^{\mathrm{BC}}\})+1} \cdot q &= \frac{\sum_{k=1}^{N} \mathbb{1}\left(C_{(k)} \geq T^{\mathrm{BC}}\right)}{1+\sum_{k=1}^{N} \mathbb{1}\left(C_{(k)} \leq -T^{\mathrm{BC}}\right)} \cdot q\\
&= \frac{\sum_{k=1}^{J} \mathbb{1}\left(C_{(k)} \geq T^{\mathrm{BC}}\right)}{1+\sum_{k=1}^{J} \mathbb{1}\left(C_{(k)} \leq -T^{\mathrm{BC}}\right)} \cdot q\\
&= \frac{(1-Z_1)+\cdots+(1-Z_J)}{1+Z_1+\cdots+Z_J} \cdot q\\
&= \left(\frac{1+J}{1+Z_1+\cdots+Z_J} - 1\right) \cdot q\,.
\end{aligned}$$

By Lemma 1(a)–(b), $Z_k \overset{\text{i.i.d.}}{\sim} \text{Bernoulli}(0.5)$, which together with Lemma 1(c) satisfy the condition of Lemma 2 and make $\rho = 0.5$. Then by Lemma 2, we have

$$\mathrm{FDR} = \mathbb{E}[\mathrm{FDP}] \leq \mathbb{E}\left[\frac{1+J}{1+Z_1+\cdots+Z_J} - 1\right] \cdot q \leq (\rho^{-1}-1)\cdot q = q\,,$$

which is the statement of Theorem 1. The complete proof of Theorem 1 is in Supp. Section S8.

###### S2.2.1 An optional, heuristic fix if the BC procedure makes no discoveries

Although the BC procedure has theoretical guarantee of FDR control, it lacks power when the number of replicates $m = n$, the target FDR threshold $q$, and the number of features $d$ are all small (e.g., $m = n = 1$, $q = 0.01$ and $d = 1000$ in Fig. S24). As a result, the BC procedure may lead to no discoveries. In that case, Clipper implements a heuristic fix—an approximate p-value Benjamini-Hochberg (aBH) procedure—to increase the power. The aBH procedure constructs an empirical null distribution of contrast scores by additionally assuming that uninteresting features' contrast scores follow a symmetric distribution around zero; it then computes approximate p-values of features based on the empirical null distribution, and finally it uses the BH procedure [1] to threshold the approximate p-values.

**Definition S7 (The aBH procedure)** *Given contrast scores $\{C_j\}_{j=1}^d$, an empirical null distribution is defined on $\mathcal{E} := \{C_j : C_j < 0; j = 1, \ldots, d\} \cup \{-C_j : C_j < 0; j = 1, \ldots, d\}$. The aBH procedure defines the approximate p-value of feature $j$ as*

$$p_j := \frac{\sum_{c \in \mathcal{E}} \mathbb{1}(c \geq C_j)}{\text{card}(\mathcal{E}) \vee 1}\,.$$

*Then it applies the BH procedure with the target FDR threshold $q$ to $\{p_j\}_{j=1}^d$ to call discoveries.*

##### S2.3 Enrichment analysis with any numbers of replicates $m$ and $n$

When $m \neq n$, the BC procedure cannot guarantee FDR control because Lemma 1 no longer holds. To control the FDR in a more general setting ($m = n$ or $m \neq n$), Clipper constructs contrast scores via permutation of replicates across conditions. The idea is that, after permutation, every feature becomes uninteresting and can serve as its own negative control.

**Definition S8 (Permutation)** *We define $\sigma$ as permutation, i.e., a bijection from the set $\{1, \cdots, m+n\}$ onto itself, and we rewrite the data $\boldsymbol{X}_1, \ldots, \boldsymbol{X}_d, \boldsymbol{Y}_1, \ldots, \boldsymbol{Y}_d$ into a matrix $\mathbf{W}$:*

$$\mathbf{W} = \begin{bmatrix} W_{11} & \cdots & W_{1m} & W_{1(m+1)} & \cdots & W_{1(m+n)} \\ & \vdots & & & \vdots & \\ W_{d1} & \cdots & W_{dm} & W_{d(m+1)} & \cdots & W_{d(m+n)} \end{bmatrix} := \begin{bmatrix} X_{11} & \cdots & X_{1m} & Y_{11} & \cdots & Y_{1n} \\ & \vdots & & & \vdots & \\ X_{d1} & \cdots & X_{dm} & Y_{d1} & \cdots & Y_{dn} \end{bmatrix}.$$

*We then apply $\sigma$ to permute the columns of $\mathbf{W}$ and obtain*

$$\mathbf{W}_\sigma := \begin{bmatrix} W_{1\sigma(1)} & \cdots & W_{1\sigma(m)} & W_{1\sigma(m+1)} & \cdots & W_{1\sigma(m+n)} \\ & \vdots & & & \vdots & \\ W_{d\sigma(1)} & \cdots & W_{d\sigma(m)} & W_{d\sigma(m+1)} & \cdots & W_{d\sigma(m+n)} \end{bmatrix},$$

*from which we obtain the permuted measurements $\left\{(\boldsymbol{X}_j^\sigma, \boldsymbol{Y}_j^\sigma)\right\}_{j=1}^d$, where*

$$\begin{aligned} \boldsymbol{X}_j^\sigma &:= \left(W_{j\sigma(1)}, \ldots, W_{j\sigma(m)}\right)^\top, \\ \boldsymbol{Y}_j^\sigma &:= \left(W_{j\sigma(m+1)}, \ldots, W_{j\sigma(m+n)}\right)^\top. \end{aligned} \tag{S18}$$

In the enrichment analysis, if two permutations $\sigma$ and $\sigma'$ satisfy that

$$\{\sigma(1), \cdots, \sigma(m)\} = \{\sigma'(1), \cdots, \sigma'(m)\},$$

then we define $\sigma$ and $\sigma'$ to be in one equivalence class. That is, permutations in the same equivalence class lead to the same division of $m+n$ replicates (from the two conditions) into two groups with sizes $m$ and $n$. In total, there are $\binom{m+n}{m}$ equivalence classes of permutations.

We define $\sigma_0$ as the identity permutation such that $\sigma_0(i) = i$ for all $i \in \{1, \cdots, m+n\}$. In addition, Clipper randomly samples $h$ equivalence classes $\sigma_1, \ldots, \sigma_h$ with equal probabilities without replacement from the other $h_{\max} := \binom{m+n}{m} - 1$ equivalence classes (after excluding the equivalence class containing $\sigma_0$). Note that $h_{\max}$ is the maximum value $h$ can take.

Clipper then obtains $\left\{(\boldsymbol{X}_j^{\sigma_0}, \boldsymbol{Y}_j^{\sigma_0}), (\boldsymbol{X}_j^{\sigma_1}, \boldsymbol{Y}_j^{\sigma_1}), \cdots, (\boldsymbol{X}_j^{\sigma_h}, \boldsymbol{Y}_j^{\sigma_h})\right\}_{j=1}^d$, where $(\boldsymbol{X}_j^{\sigma_\ell}, \boldsymbol{Y}_j^{\sigma_\ell})$ are the permuted measurements based on $\sigma_\ell$, $\ell = 0, \ldots, h$. Then Clipper computes $T_j^{\sigma_\ell} := t^{\text{minus}}(\boldsymbol{X}_j^{\sigma_\ell}, \boldsymbol{Y}_j^{\sigma_\ell})$ to indicate the degree of "interestingness" of feature $j$ reflected by $(\boldsymbol{X}_j^{\sigma_\ell}, \boldsymbol{Y}_j^{\sigma_\ell})$. Note that Clipper chooses $t^{\text{minus}}$ instead of $t^{\max}$ because empirical evidence shows that $t^{\text{minus}}$ leads to better power. Sorting $\{T_j^{\sigma_\ell}\}_{\ell=0}^h$ gives

$$T_j^{(0)} \geq T_j^{(1)} \geq \cdots \geq T_j^{(h)} .$$

Then Clipper defines the contrast score of feature $j$, $j = 1, \ldots, d$, in one of two ways:

$$C_j := \begin{cases} T_j^{(0)} - T_j^{(1)} & \text{if } T_j^{(0)} = T_j^{\sigma_0} \\ T_j^{(1)} - T_j^{(0)} & \text{otherwise} \end{cases} \qquad \textbf{minus contrast score}, \tag{S19}$$

or

$$C_j := \begin{cases} \left|T_j^{(0)}\right| & \text{if } T_j^{(0)} = T_j^{\sigma_0} > T_j^{(1)} \\ 0 & \text{if } T_j^{(0)} = T_j^{(1)} \\ -\left|T_j^{(0)}\right| & \text{otherwise} \end{cases} \qquad \textbf{maximum contrast score}. \tag{S20}$$

The intuition behind the contrast scores is that, if $C_j < 0$, then $\mathbb{1}(T_j^{(0)} = T_j^{\sigma_0}) = 0$, which means that at least one of $T_j^{\sigma_1}, \ldots, T_j^{\sigma_h}$ (after random permutation) is greater than $T_j^{\sigma_0}$ calculated from the original data (identity permutation), suggesting that feature $j$ is likely an uninteresting feature in enrichment analysis. Motivated by Gimenez and Zou [9], we propose the following procedure for Clipper to control the FDR under the target level $q \in (0, 1)$.

**Definition S9 (Gimenez-Zou (GZ) procedure for thresholding contrast scores [9])** *Given $h \in \{1, \cdots, h_{\max}\}$ and contrast scores $\{C_j\}_{j=1}^d$, $\mathcal{C} = \{|C_j| : C_j \neq 0\ ;\ j = 1, \ldots, d\}$ is defined as the set of non-zero absolute values of $C_j$'s. The GZ procedure finds a contrast-score cutoff $T^{\text{GZ}}$ based on the target FDR threshold $q \in (0, 1)$ as:*

$$T^{\text{GZ}} := \min \left\{ t \in \mathcal{C} : \frac{\frac{1}{h} + \frac{1}{h}\text{card}\left(\{j : C_j \leq -t\}\right)}{\text{card}\left(\{j : C_j \geq t\}\right) \vee 1} \leq q \right\} \tag{S21}$$

*and outputs $\left\{j : C_j \geq T^{\text{GZ}}\right\}$ as discoveries.*

**Theorem 2** *Suppose that the input data that satisfy the Clipper assumptions (S6)–(S7). Then for any $q \in (0, 1)$ and either definition of contrast scores in (S19) or (S20), the contrast-score cutoff $T^{\text{GZ}}$ found by the GZ procedure (S21) guarantees that the discoveries have the FDR under $q$:*

$$\text{FDR} = \mathbb{E}\left[ \frac{\text{card}\left(\left\{j \in \mathcal{N} : C_j \geq T^{\text{GZ}}\right\}\right)}{\text{card}\left(\{j : C_j \geq T^{\text{GZ}}\}\right) \vee 1} \right] \leq q ,$$

*where $\mathcal{N}$ denotes the set of uninteresting features.*

The proof of Theorem 2 (Supp. Section S8) is similar to that of Theorem 1 and requires two key ingredients: Lemma 2, which is also used in the proof of Theorem 1, and Lemma 3, which is similar to Lemma 1 and is about the properties of signs of $\{C_j\}_{j=1}^d$. The cutoff $T^{\mathrm{GZ}}$ can also be viewed as a stopping time of a Bernoulli process.

**Lemma 3** *For input data that satisfy the Clipper assumptions* (S6) *and* (S7)*, Clipper constructs contrast scores* $\{C_j\}_{j=1}^d$ *based on* (S20) *or* (S19)*. Denote* $S_j = \mathrm{sign}(C_j) \in \{-1, 0, 1\}$*. Then* $\{S_j\}_{j=1}^d$ *and* $\{C_j\}_{j=1}^d$ *satisfy the following properties:*

*(a)* $S_1, \ldots, S_d$ *are mutually independent ;*

*(b)* $\mathbb{P}(S_j = 1) \le \frac{1}{h+1}$ *for all* $j \in \mathcal{N}$*;*

*(c)* $\{S_j\}_{j \in \mathcal{N}} \perp \mathcal{C}$.

We note that the GZ procedure is also applicable to the enrichment analysis with equal numbers of replicates, i.e., $m = n$ (Section S2.2). We will compare the GZ procedure against the BC procedure in our results.

#### S2.4 Differential analysis with $m + n > 2$

For differential analysis, Clipper also uses permutation to construct contrast scores. When $m \neq n$, the equivalence classes of permutations are defined the same as for the enrichment analysis with $m \neq n$. When $m = n$, there is a slight change in the definition of equivalence classes of permutations: if $\sigma$ and $\sigma'$ satisfy that

$$\{\sigma(1), \cdots, \sigma(m)\} = \{\sigma'(1), \cdots, \sigma'(m)\} \text{ or } \{\sigma'(m+1), \cdots, \sigma'(2m)\},$$

then we say that $\sigma$ and $\sigma'$ are in one equivalence class. In total, there are $h_{\mathrm{total}} := \binom{m+n}{m}$ (when $m \neq n$) or $\binom{2m}{m}/2$ (when $m = n$) equivalence classes of permutations. Hence, to have more than one equivalence class, we cannot perform differential analysis with $m = n = 1$; in other words, the total number of replicates $m + n$ must be at least $3$.

Then Clipper randomly samples $\sigma_1, \ldots, \sigma_h$ with equal probabilities without replacement from the $h_{\max} := h_{\mathrm{total}} - 1$ equivalence classes that exclude the class containing $\sigma_0$, i.e., the identity permutation. Note that $h_{\max}$ is the maximum value $h$ can take. Next, Clipper computes $T_j^{\sigma_\ell} := \left| t^{\mathrm{minus}}(\boldsymbol{X}_j^{\sigma_\ell}, \boldsymbol{Y}_j^{\sigma_\ell}) \right|$, where $\boldsymbol{X}_j^{\sigma_\ell}$ and $\boldsymbol{Y}_j^{\sigma_\ell}$ are the permuted data defined in (S18), and it defines $C_j$ as the contrast score of feature $j$, $j = 1, \ldots, d$, in the same ways as in (S19) or (S20).

Same as in the enrichment analysis with $m \neq n$, Clipper also uses the GZ procedure [9] to set a cutoff on contrast scores to control the FDR under the target level $q \in (0, 1)$, following Theorem 2.

#### S2.5 Clipper variant algorithms

For nomenclature, we assign the following names to Clipper variant algorithms, each of which combines a contrast score definition with a thresholding procedure.

- **Clipper-minus-BC**: minus contrast score $C_j = t^{\mathrm{minus}}(\boldsymbol{X}_j, \boldsymbol{Y}_j)$ (S10) and BC procedure (Definition S6);
- **Clipper-minus-aBH**: minus contrast score $C_j = t^{\mathrm{minus}}(\boldsymbol{X}_j, \boldsymbol{Y}_j)$ and aBH procedure (Definition S7);
- **Clipper-minus-GZ**: minus contrast score $\tau_j = T_j^{(0)} - T_j^{(1)}$ (S19) and GZ procedure (Definition S9);
- **Clipper-max-BC**: maximum contrast score $C_j = t^{\max}(\boldsymbol{X}_j, \boldsymbol{Y}_j)$ (S11) and BC procedure;

- **Clipper-max-aBH**: maximum contrast score $C_j = t^{\max}(\boldsymbol{X}_j, \boldsymbol{Y}_j)$ and aBH procedure;
- **Clipper-max-GZ**: maximum contrast score $\tau_j = T_j^{(0)}$ (S20) and GZ procedure.

##### S2.6 R package "Clipper"

In the R package `Clipper`, the default implementation is as follows. Based on the power comparison results in Section S3 and Figs. S24, S25, S26, and S27, Clipper uses Clipper-minus-BC as the default algorithm for the enrichment analysis with equal numbers of replicates; when there are no discoveries, Clipper suggests users to increase the target FDR threshold $q$ or to use the Clipper-minus-aBH algorithm with the current $q$. For the enrichment analysis with different numbers of replicates under two conditions or the differential analysis, Clipper uses the Clipper-max-GZ algorithm by default.

#### S3 Comparison of Clipper variant algorithms

We compared Clipper variant algorithms applicable to each experimental design. Based on the comparison results, we selected a variant algorithm as the default Clipper implementation for each design.

- **1vs1 enrichment analysis.** Under each of the $12$ settings, we compared Clipper-minus-BC, Clipper-minus-aBH, Clipper-max-BC, and Clipper-max-aBH (Section S2.5), the only four Clipper variant algorithms applicable to 1vs1 enrichment analysis. The results in Fig. S24 show that, regardless of the contrast scores being minus or maximum (max), the BC procedure always guarantees the FDR control under a range of target FDR thresholds $q \in \{1\%, 2\%, \cdots, 10\%\}$. Notably, in terms of power, the two contrast scores consistently have different advantages under the two background scenarios: Clipper-max-BC has higher power under the homogeneous background, while Clipper-minus-BC is more powerful under the heterogeneous background. Considering that the heterogeneous scenario is prevalent in high-throughput biological data, the minus contrast score is preferred. As the power of Clipper-minus-BC drops when $q$ is too small ($q \leq 3\%$) and $d$ is not too large ($d = 1000$), we consider the aBH procedure as an alternative to control the FDR. The results show that Clipper-minus-aBH is indeed more powerful when Clipper-minus-BC lacks power; however, Clipper-minus-aBH cannot guarantee the exact FDR control as Clipper-minus-BC does. Therefore, Clipper uses **Clipper-minus-BC** by default in 1vs1 enrichment analysis, and it recommends users to increase $q$ when too few discoveries are made; if users reject this option, then Clipper would use Clipper-minus-aBH to increase power for the current $q$.
- **2vs1 enrichment analysis.** Under each of the $6$ settings, we compared Clipper-minus-GZ and Clipper-max-GZ (Section S2.5), the only two Clipper variant algorithms applicable to 2vs1 enrichment analysis. For either algorithm, we further compared two numbers of permutation equivalence classes: $h = 1$ or $2$, where the latter is $h_{\max} = \binom{3}{1} - 1$—the maximum number of equivalence classes that do not include the identity permutation. Note that $h$ is a required input parameter for the GZ procedure. The results in Fig. S25 show that, regardless of $h$ and the contrast score definition—maximum (max) or minus, the GZ procedure always guarantees the FDR control under all target FDR thresholds $q \in \{1\%, 2\%, \cdots, 10\%\}$. In terms of power, Clipper-max-GZ($h = 1$) is consistently more powerful than the other three Clipper variants under all settings. Therefore, Clipper uses **Clipper-max-GZ($h = 1$)** by default in enrichment analysis with unequal numbers of replicates under two conditions.
- **3vs3 enrichment analysis.** Under each of the $12$ settings, we compared five Clipper variant algorithms: Clipper-minus-BC, Clipper-minus-aBH, Clipper-max-BC, Clipper-max-aBH, and

Clipper-max-GZ (Section S2.5). Fig. S26 shows the comparison of the first four variants: regardless of the contrast scores being minus or maximum (max), the BC procedure simultaneously guarantees the FDR control and achieves good power under a range of target FDR thresholds $q \in \{1\%, 2\%, \cdots, 10\%\}$. Similar to the results in the 1vs1 enrichment analysis, Clipper-max-BC has higher power under the homogeneous background, while Clipper-minus-BC is more powerful under the heterogeneous background. By the same reasoning—the prevalent heterogeneous scenarios in high-throughput biological data—we prefer the minus contrast score. Unlike the 1vs1 enrichment analysis, here Clipper-minus-BC is consistently as powerful as Clipper-minus-aBH, even when $q$ is small, but Clipper-minus-aBH cannot guarantee the exact FDR control. Therefore, Clipper-minus-BC achieves the overall best performance among the first four Clipper variants. Given that the GZ procedure is also applicable to this setting, we further compared Clipper-minus-BC with Clipper-max-GZ($h = 1$), the most powerful Clipper variant with the GZ procedure and the default Clipper implementation in the 2vs1 enrichment and differential analyses and the 3vs3 differential analysis. The results in Fig. S28 show that while both **Clipper-minus-BC** and Clipper-max-GZ($h = 1$) control the FDR, the former is more powerful. Hence, we will use Clipper-minus-BC as the default when both conditions have more than one and the same number of replicates.

Under the simulation settings from Gaussian distributions, we also compared Clipper-minus-BC with another Clipper variant using the BC procedure and the $t$ statistic as the contrast score (Clipper-t), where the $t$ statistic is from the two-sample $t$ test. Fig. S13 shows that, although Clipper-t always guarantees the FDR control under a range of target FDR thresholds $q \in \{1\%, 2\%, \cdots, 10\%\}$, it has lower power compared to Clipper-minus-BC, our default Clipper for enrichment analysis with equal numbers of replicates. Based on this result, we did not consider the $t$ statistic as an alternative contrast score for Clipper.

- **2vs1 differential analysis.** Similar to 2vs1 enrichment analysis, under each of the $6$ settings, we compared Clipper-minus-GZ and Clipper-max-GZ (Section S2.5) with $h = 1$ or $2$. The results in Fig. S25 show that, regardless of $h$ and the contrast score definition—maximum (max) or minus, the GZ procedure always guarantees the FDR control under a range of target FDR thresholds $q \in \{1\%, 2\%, \cdots, 10\%\}$. Notably, in terms of power, Clipper-minus-GZ($h = 2$) is the most powerful when when $q$ is very small ($q \leq 2\%$) under Poisson and negative binomial settings, while Clipper-max-GZ($h = 1$) is the most powerful otherwise. Considering that Clipper-max-GZ($h = 1$) outperforms the other three Clipper variants in most cases, Clipper uses **Clipper-max-GZ($h = 1$)** by default in 2vs1 differential analysis, and it recommends users to use Clipper-minus-GZ($h = 2$) when too few discoveries are made.

- **3vs3 differential analysis.** Under each of the $12$ settings, we compared Clipper-minus-GZ, and Clipper-max-GZ (Section S2.5) with $h = 1$, $3$ or $9$, where $h = 9$ is $h_{\max} = \binom{6}{3}/2 - 1$—the maximum number of equivalence classes that do not include the identity permutation. The results in Fig. S27 show that, regardless of $h$ and the contrast score definition—maximum (max) or minus, the GZ procedure always guarantees the FDR control under a range of target FDR thresholds $q \in \{1\%, 2\%, \cdots, 10\%\}$. In terms of power, Clipper-max-GZ($h = 1$) is consistently more powerful than the other Clipper variant algorithms under all settings. Therefore, Clipper uses **Clipper-max-GZ($h = 1$)** by default in 3vs3 differential analysis.

  Under the simulation settings from Gaussian distributions, we also compared Clipper-max-GZ with another Clipper variant using the GZ procedure and the $t$ statistic to calculate the degree of interestingness (Clipper-t), where the $t$ statistic is from the two-sample $t$ test. Fig. S14 shows that, although Clipper-t always guarantees the FDR control under a range of target FDR thresholds

$q \in \{1\%, 2\%, \cdots, 10\%\}$, it has lower power compared to Clipper-max-GZ, our default Clipper for differential analysis. Based on this result, we did not consider the $t$ statistic as an alternative contrast scores for Clipper.

In summary, whenever Clipper-minus-BC is applicable (enrichment analysis with equal number of replicates under two conditions), it is chosen as the default Clipper implementation; otherwise, Clipper-max-GZ($h = 1$) is the default.

### S4 Data generation and detailed implementation of the paired approach (a p-value calculation approach) in simulation studies

We describe how we simulated data and how we implemented the paired approach in different simulation settings: 1vs1 enrichment analysis, 2vs1 enrichment analysis, 3vs3 enrichment analysis, 2vs1 differential analysis, and 3vs3 differential analysis, combined with three distribution families (Gaussian, Poisson, and negative binomial) and two background scenarios (homogeneous and heterogeneous). Under some settings, we considered different numbers of features and the existence of outliers.

In each simulation setting, we generated $200$ simulated datasets, computed an FDP and an empirical power on each dataset, and averaged the $200$ FDPs and $200$ empirical powers to approximate the FDR and power, repsectively. For notation simplicity, we use $N(\mu, \sigma^2)$ to denote the Gaussian distribution with mean $\mu$ and variance $\sigma^2$, $\mathrm{Pois}(\lambda)$ to denote the Poisson distribution with mean $\lambda$, and $\mathrm{NB}(\mu, \theta)$ to denote the negative binomial distribution with mean $\mu$ and dispersion $\theta$ (such that its variance equals $\mu + \theta\mu^2$).

For each design and analysis, we compared the default Clipper implementation with other generic FDR control methods. Specifically, seven generic methods (BH-pool, qvalue-pool, BH-pair-mis, qvalue-pair-mis, BH-pair-2as1, qvalue-pair-2as1, and locfdr-emp) are included in all designs and analyses. The two methods relying on correct model specification, BH-pair-correct and qvalue-pair-correct, are only included in the 3vs3 enrichment and differential analyses, because it is almost impossible to correctly specify a model with fewer than three replicates per condition. The permutation-based method, locfdr-swap, is excluded from the 1vs1 enrichment analysis because it requires at least one condition to have more than one replicate.

In addition to the above designs and analyses, we also compared the default Clipper implementation with BH-pair methods that use parametric or non-parametric tests to calculate p-values when the numbers of replicates are $10$ under both conditions for enrichment analysis, i.e., 10vs10 enrichment analysis.

#### S4.1 1vs1 enrichment analysis

We simulated data with $d = 1000$ and $10{,}000$ features under two background scenarios and three distributional families—a total of $12$ settings. In each setting, $10\%$ of the features are interesting ($\mu_{Xj} > \mu_{Yj}$), and the rest are uninteresting (with $\mu_{Xj} = \mu_{Yj}$). Recall that $\mathcal{N}$ denotes the set of uninteresting features.

##### Gaussian distribution

We simulated data from Gaussian using the following procedure:

- Under the homogeneous background scenario, we set $\mu_{Yj} = 0$ for all $d$ features. For uninteresting features, we set $\mu_{Xj} = \mu_{Yj} = 0$ for $j \in \mathcal{N}$. For interesting features, we generated $\{\mu_{Xj}\}_{j \notin \mathcal{N}}$ i.i.d. from $N(5, 1)$.

- Under the heterogeneous background scenario, we generated $\{\mu_{Yj}\}_{j=1}^{d}$ i.i.d. from $N(0, 2^2)$. For uninteresting features, we set $\mu_{Xj} = \mu_{Yj}$ for $j \in \mathcal{N}$. For interesting features, we generated $\{\mu_{Xj}\}_{j \notin \mathcal{N}}$ i.i.d. from $N(5, 1)$.
- We independently generated $X_{j1}$ from $N(\mu_{Xj}, 1)$ and $Y_{j1}$ from $N(\mu_{Yj}, 1)$, $j = 1, \ldots, d$.

To implement the misspecified paired approach (as in BH-pair-mis and qvalue-pair-mis), we assumed that the null distribution of $X_{j1} - Y_{j1}$, $j = 1, \ldots, d$ is $N(0, \hat{\sigma}^2)$, where

$$\hat{\sigma}^2 = \frac{1}{d-1}\sum_{j=1}^{d}\left(X_{j1} - \frac{1}{d}\sum_{j=1}^{d} X_{j1}\right)^2 + \frac{1}{d-1}\sum_{j=1}^{d}\left(Y_{j1} - \frac{1}{d}\sum_{j=1}^{d} Y_{j1}\right)^2 .$$

This is a misspecified model that assumes that $\mu_{Xj}$'s are all equal and so are $\mu_{Yj}$'s. Then we computed the p-value of feature $j = 1, \ldots, d$ as the right tail probability of $X_{j1} - Y_{j1}$ in $N(0, \hat{\sigma}^2)$, i.e., $1 - \Phi\left(\frac{X_{j1} - Y_{j1}}{\hat{\sigma}}\right)$, where $\Phi$ is the cumulative distribution function of $N(0, 1)$.

To implement the 2as1 paired approach (as in BH-pair-2as1 and qvalue-pair-2as1), we treated $N(Y_{j1}, 1)$ conditioning on the observed $Y_{j1}$ as the null distribution of $X_{j1}$. Then we calculated the p-value of feature $j = 1, \ldots, d$ as the right tail probability of $X_{j1}$ in $N(Y_{j1}, 1)$, i.e., $1 - \Phi\left(X_{j1} - Y_{j1}\right)$.

**Poisson distribution**

We simulated data from Poisson using the following procedure:

- Under the homogeneous background scenario, we set $\mu_{Yj} = 20$ for all $d$ features. For uninteresting features, we set $\mu_{Xj} = \mu_{Yj} = 20$ for $j \in \mathcal{N}$. For interesting features, we generated $\{\mu_{Xj}\}_{j \notin \mathcal{N}}$ i.i.d. from $\mathrm{Pois}(40)$.
- Under the heterogeneous background scenario, we generated $\{\mu_{Yj}\}_{j=1}^{d}$ i.i.d. from $\mathrm{Pois}(20)$. For uninteresting features, we set $\mu_{Xj} = \mu_{Yj}$ for $j \in \mathcal{N}$. For interesting features, we generated $\{\mu_{Xj}\}_{j \notin \mathcal{N}}$ i.i.d. from $\mathrm{Pois}(40)$.
- We independently generated $X_{j1}$ from $\mathrm{Pois}(\mu_{Xj})$ and $Y_{j1}$ from $\mathrm{Pois}(\mu_{Yj})$, $j = 1, \ldots, d$.

To implement the misspecified paired approach (as in BH-pair-mis and qvalue-pair-mis), we first defined a log-transformation $f(x) = \log(x + 0.01)$, which we applied to $X_{j1}$ and $Y_{j1}$, $j = 1, \ldots, d$. We assumed that the null distribution of $f(X_{j1}) - f(Y_{j1})$, $j = 1, \ldots, d$ is $N(0, \hat{\sigma}^2)$, where

$$\hat{\sigma}^2 = \frac{1}{d-1}\sum_{j=1}^{d}\left(f(X_{j1}) - \frac{1}{d}\sum_{j=1}^{d} f(X_{j1})\right)^2 + \frac{1}{d-1}\sum_{j=1}^{d}\left(f(Y_{j1}) - \frac{1}{d}\sum_{j=1}^{d} f(Y_{j1})\right)^2 .$$

This model misspecifies the Poisson distribution as the log-normal distribution.

Then we computed the p-value of feature $j = 1, \ldots, d$ as the right tail probability of $f(X_{j1}) - f(Y_{j1})$ in $N(0, \hat{\sigma}^2)$, i.e., $1 - \Phi\left(\frac{f(X_{j1}) - f(Y_{j1})}{\hat{\sigma}}\right)$, where $\Phi$ is the cumulative distribution function of $N(0, 1)$.

To implement the 2as1 paired approach (as in BH-pair-2as1 and qvalue-pair-2as1), we treated $\mathrm{Pois}(Y_{j1})$ conditioning on the observed $Y_{j1}$ as the null distribution of $X_{j1}$. Then we calculated the p-value of feature $j = 1, \ldots, d$ as the right tail probability of $X_{j1}$ in $\mathrm{Pois}(Y_{j1})$, i.e., $\mathbb{P}(Z \geq X_{j1})$ where $Z \sim \mathrm{Pois}(Y_{j1})$.

**Negative binomial distribution**

We simulated data from negative binomial using the following procedure:

- Under the homogeneous background scenario, we set $\mu_{Yj} = 20$ for all $d$ features. For uninteresting features, we set $\mu_{Xj} = \mu_{Yj} = 20$ for $j \in \mathcal{N}$. For interesting features, we generated $\{\mu_{Xj}\}_{j \notin \mathcal{N}}$ i.i.d. from $\mathrm{NB}(45, 45^{-1})$.

- Under the heterogeneous background scenario, we generated $\{\mu_{Yj}\}_{j=1}^{d}$ i.i.d. from $\mathrm{NB}(20, 20^{-1})$. For uninteresting features, we set $\mu_{Xj} = \mu_{Yj}$ for $j \in \mathcal{N}$. For interesting features, we generated $\{\mu_{Xj}\}_{j \notin \mathcal{N}}$ i.i.d. from $\mathrm{NB}(45, 45^{-1})$.

- We independently generated $X_{j1}$ from $\mathrm{NB}(\mu_{Xj}, \mu_{Xj}^{-1})$ and $Y_{j1}$ from $\mathrm{NB}(\mu_{Yj}, \mu_{Yj}^{-1})$, $j = 1, \ldots, d$.

To implement the misspecified paired approach (as in BH-pair-mis and qvalue-pair-mis), we assumed that for each uninteresting feature $j$, $Y_{j1}$ and $X_{j1}$ follow the same Poisson distribution. We calculated the p-value of feature $j$ from a two-sample Poisson test for the null hypothesis $H_0 : \mu_{Xj} = \mu_{Yj}$ against the alternative hypothesis $H_1 : \mu_{Xj} > \mu_{Yj}$ using function `poisson.test` in R package `stats`.

To implement the 2as1 paired approach (as in BH-pair-2as1 and qvalue-pair-2as1), we treated $\mathrm{NB}(Y_{j1}, Y_{j1}^{-1})$ conditioning on the observed $Y_{j1}$ as the null distribution of $X_{j1}$. Then we calculated the p-value of feature $j = 1, \ldots, d$ as the right tail probability of $X_{j1}$ in $\mathrm{NB}(Y_{j1}, Y_{j1}^{-1})$.

#### S4.2 2vs1 enrichment analysis

We simulated data with $d = 10{,}000$ features under two background scenarios and three distributional families—a total of $6$ settings. In each setting, $10\%$ of the features are interesting ($\mu_{Xj} > \mu_{Yj}$) and the rest are uninteresting (with $\mu_{Xj} = \mu_{Yj}$). Recall that $\mathcal{N}$ denotes the set of uninteresting features.

##### Gaussian distribution

We simulated data from Gaussian using the following procedure:

- Under the homogeneous background scenario, we set $\mu_{Yj} = 0$ for all $d$ features. For uninteresting features, we set $\mu_{Xj} = \mu_{Yj} = 0$ for $j \in \mathcal{N}$. For interesting features, we generated $\{\mu_{Xj}\}_{j \notin \mathcal{N}}$ i.i.d. from $N(5, 1)$.

- Under the heterogeneous background scenario, we generated $\{\mu_{Yj}\}_{j \in \mathcal{N}}$ i.i.d. from $N(0, 2^2)$ and set $\mu_{Xj} = \mu_{Yj}$ for $j \in \mathcal{N}$. We next generated $\{\mu_{Yj}\}_{j \notin \mathcal{N}}$ i.i.d. from $N(0, 2^2)$ and $\{\mu_{Xj}\}_{j \notin \mathcal{N}}$ i.i.d. from $N(5, 1)$.

- We independently generated $X_{j1}$ and $X_{j2}$ from $N(\mu_{Xj}, 1)$ and $Y_{j1}$ from $N(\mu_{Yj}, 1)$, $j = 1, \ldots, d$.

To implement the misspecified paired approach (as in BH-pair-mis and qvalue-pair-mis), we assumed that the null distribution of $\frac{1}{2}(X_{j1} + X_{j2}) - Y_{j1}$, $j = 1, \ldots, d$, is $N(0, \hat{\sigma}^2)$, where

$$\hat{\sigma}^2 = \frac{1}{2(2d-1)} \sum_{j=1}^{d} \sum_{i=1}^{2} \left( X_{ji} - \frac{1}{2d} \sum_{j=1}^{d} \sum_{i=1}^{2} X_{ji} \right)^2 + \frac{1}{d-1} \sum_{j=1}^{d} \left( Y_{j1} - \frac{1}{d} \sum_{j=1}^{d} Y_{j1} \right)^2 .$$

This is a misspecified model that assumes $\mu_{Xj}$'s are all equal and so are $\mu_{Yj}$'s. Then we computed the p-value of feature $j = 1, \ldots, d$ as the right tail probability of $\frac{1}{2}(X_{j1} + X_{j2}) - Y_{j1}$ in $N(0, \hat{\sigma}^2)$, i.e., $1 - \Phi\left( \frac{\frac{1}{2}(X_{j1}+X_{j2}) - Y_{j1}}{\hat{\sigma}} \right)$, where $\Phi$ is the cumulative distribution function of $N(0, 1)$.

To implement the 2as1 paired approach (as in BH-pair-2as1 and qvalue-pair-2as1), we treated $N(Y_{j1}, 1/2)$ conditioning on the observed $Y_{j1}$ as the null distribution of $\frac{1}{2}(X_{j1} + X_{j2})$. Then we calculated the p-value of feature $j = 1, \ldots, d$ as the right tail probability of $\frac{1}{2}(X_{j1} + X_{j2})$ in $N(Y_{j1}, 1/2)$, i.e., $1 - \Phi\left( \frac{\frac{1}{2}(X_{j1}+X_{j2}) - Y_{j1}}{1/\sqrt{2}} \right)$.

##### Poisson distribution

We simulated data from Poisson using the following procedure:

- Under the homogeneous background scenario, we set $\mu_{Yj} = 20$ for all $d$ features. For uninteresting features, we set $\mu_{Xj} = \mu_{Yj} = 20$ for $j \in \mathcal{N}$. For interesting features, we generated $\{\mu_{Xj}\}_{j \notin \mathcal{N}}$ i.i.d. from $\mathrm{Pois}(40)$.

- Under the heterogeneous background scenario, we generated $\{\mu_{Yj}\}_{j=1}^{d}$ i.i.d. from $\mathrm{Pois}(20)$. For uninteresting features, we set $\mu_{Xj} = \mu_{Yj}$ for $j \in \mathcal{N}$. For interesting features, we generated $\{\mu_{Xj}\}_{j \notin \mathcal{N}}$ i.i.d. from $\mathrm{Pois}(40)$.

- We independently generated $X_{j1}$ and $X_{j2}$ from $\mathrm{Pois}(\mu_{Xj})$ and $Y_{j1}$ from $\mathrm{Pois}(\mu_{Yj})$, $j = 1, \ldots, d$.

To implement the misspecified paired approach (as in BH-pair-mis and qvalue-pair-mis), we first defined a log-transformation $f(x) = \log(x + 0.01)$, which we applied to $X_{j1}$ and $Y_{j1}$, $j = 1, \ldots, d$. We assumed that the null distribution of $f(X_{j1}) + f(X_{j2}) - 2f(Y_{j1})$, $j = 1, \ldots, d$ is $N(0, \hat{\sigma}^2)$, where

$$\hat{\sigma}^2 = \frac{6}{d-1} \sum_{j=1}^{d} \left( f(Y_{j1}) - \frac{1}{d} \sum_{j=1}^{d} f(Y_{j1}) \right)^2 .$$

This model misspecifies the Poisson distribution as the log-normal distribution.

Then we computed the p-value of feature $j = 1, \ldots, d$ as the right tail probability of $f(X_{j1}) + f(X_{j2}) - 2f(Y_{j1})$ in $N(0, \hat{\sigma}^2)$, i.e., $1 - \Phi\left(\frac{f(X_{j1}) + f(X_{j2}) - 2f(Y_{j1})}{\hat{\sigma}}\right)$, where $\Phi$ is the cumulative distribution function of $N(0, 1)$.

To implement the 2as1 paired approach (as in BH-pair-2as1 and qvalue-pair-2as1), we assumed that for each uninteresting feature $j$, $X_{j1}$ and $X_{j2}$ independently follow $\mathrm{Pois}(Y_{j1})$ conditioning on the observed $Y_{j1}$. Then we calculated the p-value of feature $j = 1, \ldots, d$ by performing a one-sample Poisson test using the R function `poisson.test` for the null hypothesis $H_0 : \mu_{X_j} = Y_{j1}$ against the alternative hypothesis $H_1 : \mu_{X_j} > Y_{j1}$.

##### Negative binomial distribution

We simulated data from negative binomial using the following procedure:

- Under the homogeneous background scenario, we set $\mu_{Yj} = 20$ for all $d$ features. For uninteresting features, we set $\mu_{Xj} = \mu_{Yj} = 20$ for $j \in \mathcal{N}$. For interesting features, we generated $\{\mu_{Xj}\}_{j \notin \mathcal{N}}$ i.i.d. from $\mathrm{NB}(45, 45^{-1})$.

- Under the heterogeneous background scenario, we generated $\{\mu_{Yj}\}_{j=1}^{d}$ i.i.d. from $\mathrm{NB}(20, 20^{-1})$. For uninteresting features, we set $\mu_{Xj} = \mu_{Yj}$ for $j \in \mathcal{N}$. For interesting features, we generated $\{\mu_{Xj}\}_{j \notin \mathcal{N}}$ i.i.d. from $\mathrm{NB}(45, 45^{-1})$.

- We independently generated $X_{j1}$ and $X_{j2}$ from $\mathrm{NB}(\mu_{Xj}, \mu_{Xj}^{-1})$ and $Y_{j1}$ from $\mathrm{NB}(\mu_{Yj}, \mu_{Yj}^{-1})$, $j = 1, \ldots, d$.

To implement the misspecified paired approach (as in BH-pair-mis and qvalue-pair-mis), we assumed that for each uninteresting feature $j$, $X_{ji}$, $i = 1, 2$ and $Y_{j1}$ follow the same Poisson distribution. We calculated the p-value of feature $j$ from a two-sample Poisson test for the null hypothesis $H_0 : \mu_{Xj} = \mu_{Yj}$ against the alternative hypothesis $H_1 : \mu_{Xj} > \mu_{Yj}$ using the function `poisson.test` in R package `stats`.

To implement the 2as1 paired approach (as in BH-pair-2as1 and qvalue-pair-2as1), we treated $\mathrm{NB}(2Y_{j1}, (2Y_{j1})^{-1})$ conditioning on the observed $Y_{j1}$ as the null distribution of $X_{j1} + X_{j2}$. Then we calculated the p-value of feature $j = 1, \ldots, d$ as the right tail probability of $X_{j1} + X_{j2}$ in $\mathrm{NB}(2Y_{j1}, (2Y_{j1})^{-1})$.

#### S4.3 3vs3 enrichment analysis

We simulated data with and without outliers under two background scenarios and three distributional families—a total of $12$ settings. In each setting, we generated $d = 10{,}000$ features, among which $10\%$ are interesting (with $\mu_{Xj} > \mu_{Yj}$) and the rest are uninteresting (with $\mu_{Xj} = \mu_{Yj}$). For the results in Fig. S11, we simulated data without outliers under two background scenarios and three distributional families using two more proportions of interesting features: $20\%$ and $40\%$. The data generation under the Gaussian, Poisson, and negative binomial distributions is the same as the settings with $10\%$ interesting features.

Under the settings with outliers, we generated $\{O_{ji}^X : j = 1, \ldots, d; i = 1, \ldots, 3\}$ and $\{O_{ji}^Y : j = 1, \ldots, d; i = 1, \ldots, 3\}$ i.i.d. from $\mathrm{Bernoulli}(0.1)$, where $O_{ji}^X = 1$ or $O_{ji}^Y = 1$ indicates $X_{ji}$ or $Y_{ji}$ is an outlier, respectively. Under settings without outliers, $O_{ji}^X = O_{ij}^Y = 0$ for all $j = 1, \ldots, d; i = 1, \ldots, 3$.

**Gaussian distribution**

- Under the homogeneous background scenario, we set $\mu_{Yj} = 0$ for all $d$ features. For uninteresting features, we set $\mu_{Xj} = \mu_{Yj} = 0$ for $j \in \mathcal{N}$. For interesting features, we generated $\{\mu_{Xj}\}_{j \notin \mathcal{N}}$ i.i.d. from $N(5, 1)$.

- Under the heterogeneous background scenario, we generated $\{\mu_{Yj}\}_{j=1}^d$ i.i.d. from $N(0, 2^2)$. For uninteresting features, we set $\mu_{Xj} = \mu_{Yj}$ for $j \in \mathcal{N}$. For interesting features, we generated $\{\mu_{Xj}\}_{j \notin \mathcal{N}}$ i.i.d. from $N(5, 1)$.

- We independently generated $X_{ji}$ from $N(\mu_{Xj}, 1)$ if $O_{ji}^X = 0$ or from the top $1\%$ percentile of $N(\mu_{Xj}, 1)$ if $O_{ji}^X = 1$, $j = 1, \ldots, d$; $i = 1, \ldots, 3$. Similarly, we independently generated $Y_{ji}$ from $N(\mu_{Yj}, 1)$ if $O_{ji}^Y = 0$ or from the top $1\%$ percentile of $N(\mu_{Yj}, 1)$ if $O_{ji}^Y = 1$, $j = 1, \ldots, d$; $i = 1, \ldots, 3$.

- For the results in Additional File 1: Fig. S13, under the heterogeneous background scenario, we generated $\{\mu_{Yj}\}_{j=1}^d$ i.i.d. from $N(0, 2^2)$, and $\{s_j\}_{j=1}^d$ i.i.d. from a uniform distribution $U(0.5, 2)$. For uninteresting features, we set $\mu_{Xj} = \mu_{Yj}$ for $j \in \mathcal{N}$. For interesting features, we generated $\{\mu_{Xj}\}_{j \notin \mathcal{N}}$ i.i.d. from $N(5, 1)$. We then independently generated $X_{ji}$ from $N(\mu_{Xj}, s_j^2)$ if $O_{ji}^X = 0$ or from the top $1\%$ percentile of $N(\mu_{Xj}, s_j^2)$ if $O_{ji}^X = 1$, $j = 1, \ldots, d$; $i = 1, \ldots, 3$. Similarly, we independently generated $Y_{ji}$ from $N(\mu_{Yj}, s_j^2)$ if $O_{ji}^Y = 0$ or from the top $1\%$ percentile of $N(\mu_{Yj}, s_j^2)$ if $O_{ji}^Y = 1$, $j = 1, \ldots, d$; $i = 1, \ldots, 3$.

- For the results in Additional File 1: Fig. S12, we generated correlated features. We first selected $10$ groups of features ($2$ groups of interesting features and $8$ groups of uninteresting features), with each group containing $200$ features. For each group $k$, we used $k_1, \ldots, k_{200}$ to denote the indices of the $200$ features within that group and generated $\{X_{k_l i}\}_{l=1}^{200}$ from a multivariate Gaussian distribution $N(\boldsymbol{\mu}_k, \boldsymbol{\Sigma}_k)$, where $\boldsymbol{\mu}_k = (\mu_{Xk_1}, \ldots, \mu_{Xk_{200}})$ and $\boldsymbol{\Sigma}_k$ is a matrix with diagonal entries as $1$ and other entries as a fixed correlation. In our simulation, the fixed correlation took two values: $0.2$ and $0.4$.

To implement the correct paired approach (as in BH-pair-correct and qvalue-pair-correct), we calculated the p-value of feature $j$ from a two-sample t-test with equal variance for the null hypothesis $H_0 : \mu_{Xj} = \mu_{Yj}$ against the alternative hypothesis $H_1 : \mu_{Xj} > \mu_{Yj}$.

To implement the misspecified paired approach (as in BH-pair-mis and qvalue-pair-mis), we calculated the p-value of feature $j$ from a two-sample t-test with unequal variance for the null hypothesis $H_0 : \mu_{X_j} = \mu_{Y_j}$ against the alternative hypothesis $H_1 : \mu_{X_j} > \mu_{Y_j}$ .

To implement the 2as1 paired approach (as in BH-pair-2as1 and qvalue-pair-2as1), we assumed that for each uninteresting feature $j$, $X_{ji}$, $i = 1, \ldots, 3$ are i.i.d. Gaussian with mean $\bar{Y}_j$ conditioning on the observed $\bar{Y}_j$ and unknown variance. We calculated the p-value of feature $j$ using a one-sample t-test for the null hypothesis $H_0 : \mu_{X_j} = \bar{Y}_j$ against the alternative hypothesis $H_1 : \mu_{X_j} > \bar{Y}_j$.

**Poisson distribution**

We simulated data from Poisson using the following procedure:

- Under the homogeneous background scenario, we set $\mu_{Yj} = 20$ for all $d$ features. For uninteresting features, we set $\mu_{Xj} = \mu_{Yj} = 20$ for $j \in \mathcal{N}$. For interesting features, we generated $\{\mu_{Xj}\}_{j \notin \mathcal{N}}$ i.i.d. from $\text{Pois}(40)$.
- Under the heterogeneous background scenario, we generated $\{\mu_{Yj}\}_{j=1}^{d}$ i.i.d. from $\text{Pois}(20)$. For uninteresting features, we set $\mu_{Xj} = \mu_{Yj}$ for $j \in \mathcal{N}$. For interesting features, we generated $\{\mu_{Xj}\}_{j \notin \mathcal{N}}$ i.i.d. from $\text{Pois}(40)$.
- We independently generated $X_{ji}$ from $\text{Pois}(\mu_{Xj})$ if $O_{ji}^X = 0$ or from the top $1\%$ percentile of $\text{Pois}(\mu_{Xj})$ if $O_{ji}^X = 1$, $j = 1, \ldots, d$, $i = 1, \ldots, 3$. Similarly, we independently generated $Y_{ji}$ from $\text{Pois}(\mu_{Yj})$ if $O_{ji}^Y = 0$ or from the top $1\%$ percentile of $\text{Pois}(\mu_{Yj})$ if $O_{ji}^Y = 1$, $j = 1, \ldots, d; 1, \ldots, 3$.

To implement the correct paired approach (as in BH-pair-correct and qvalue-pair-correct), we calculated the p-value of feature $j$ by performing a two-sample Poisson test for the null hypothesis $H_0 : \mu_{Xj} = \mu_{Yj}$ against the alternative hypothesis $H_1 : \mu_{Xj} > \mu_{Yj}$ using the function `poisson.test` in R package `stats`.

To implement the misspecified paired approach (as in BH-pair-mis and qvalue-pair-mis), we first defined a log-transformation $f(x) = \log(x + 0.01)$, which we applied to $X_{ji}$ and $Y_{ji}$, $j = 1, \ldots, d$; $i = 1, \ldots, 3$. We assumed that for each uninteresting feature $j$, $\{f(X_{ji})\}_{i=1}^3$ and $\{f(Y_{ji})\}_{i=1}^3$ follow Gaussian distributions with mean $\mu_{f(Xj)}$ and $\mu_{f(Yj)}$, respectively. Then we computed the p-value of feature $j$ using a two-sample equal variance t-test for the null hypothesis $H_0 : \mu_{f(Xj)} = \mu_{f(Yj)}$ against the alternative hypothesis $H_1 : \mu_{f(Xj)} > \mu_{f(Yj)}$.

To implement the 2as1 paired approach (as in BH-pair-2as1 and qvalue-pair-2as1), we assumed that for each uninteresting feature $j$, $\{X_{ji}\}_{i=1}^3$ follow $\text{Pois}(\bar{Y}_j)$ conditioning on the observed $\bar{Y}_j$. We calculated the p-value of feature $j$ by performing a one-sample Poisson test for the null hypothesis $H_0 : \mu_{Xj} = \bar{Y}_j$ against the alternative hypothesis $H_1 : \mu_{Xj} > \bar{Y}_j$ using R function `poisson.test` from package `stats`.

**Negative binomial distribution**

We simulated data from negative binomial using the following procedure:

- Under the homogeneous background scenario, we set $\mu_{Yj} = 20$ for all $d$ features. For uninteresting features, we set $\mu_{Xj} = \mu_{Yj} = 20$ for $j \in \mathcal{N}$. For interesting features, we generated $\{\mu_{Xj}\}_{j \notin \mathcal{N}}$ i.i.d. from $\text{NB}(45, 45^{-1})$.
- Under the heterogeneous background scenario, we generated $\{\mu_{Yj}\}_{j=1}^{d}$ i.i.d. from $\text{NB}(20, 20^{-1})$. For uninteresting features, we set $\mu_{Xj} = \mu_{Yj}$ for $j \in \mathcal{N}$. For interesting features, we generated $\{\mu_{Xj}\}_{j \notin \mathcal{N}}$ i.i.d. from $\text{NB}(45, 45^{-1})$.

- We independently generated $X_{ji}$ from $\mathrm{NB}(\mu_{Xj}, \mu_{Xj}^{-1})$ if $O_{ji}^X = 0$ or from the top $1\%$ percentile of $\mathrm{NB}(\mu_{Xj}, \mu_{Xj}^{-1})$ if $O_{ji}^X = 1$, $j = 1, \ldots, d$, $i = 1, \ldots, 3$. Similarly, we independently generated $Y_{ji}$ from $\mathrm{NB}(\mu_{Yj}, \mu_{Yj}^{-1})$ if $O_{ji}^Y = 0$ or from the top $1\%$ percentile of $\mathrm{NB}(\mu_{Yj}, \mu_{Yj}^{-1})$ if $O_{ji}^Y = 1$, $j = 1, \ldots, d$, $i = 1, \ldots, 3$.

To implement the correct paired approach (as in BH-pair-correct and qvalue-pair-correct), we performed a two-sample negative binomial test for the null hypothesis $H_0 : \mu_{Xj} = \mu_{Yj}$ against the alternative $H_1 : \mu_{Xj} > \mu_{Yj}$ using $T_j := \sum_{i=1}^3 X_{ji} - \sum_{i=1}^3 Y_{ji}$ as the test statistic. We computed the p-value of feature $j$ as the right tail probability

$$\mathbb{P}(T_j \geq t_j) = \sum_{k_1=0}^{\infty} \sum_{k_2=k_1+t_j}^{\infty} \mathbb{P}\left(\sum_{i=1}^3 X_{ji} \geq k_2\right) \mathbb{P}\left(\sum_{i=1}^3 Y_{ji} = k_1\right),$$

where $t_j$ is the realization of $T_j$, $\mathbb{P}(\sum_{i=1}^3 X_{ji} \geq k_2)$ and $\mathbb{P}(\sum_{i=1}^3 Y_{ji} = k_1)$ can be estimated from the null distribution of $X_{ji}$ and $Y_{ji}$, $j = 1, \ldots, d$; $i = 1, \ldots, 3$. As $\sum_{i=1}^3 X_{ji}$ and $\sum_{i=1}^3 Y_{ji}$ follow the same distribution under null, we estimated $\mu_{Xj}$ and $\mu_{Yj}$ as $\hat{\mu}_{Xj} = \hat{\mu}_{Yj} := (\sum_{i=1}^3 X_{ji} + \sum_{i=1}^3 Y_{ji})/6$. Then, we calculated $\mathbb{P}(\sum_{i=1}^3 X_{ji} \geq k_2)$ and $\mathbb{P}(\sum_{i=1}^3 Y_{ji} = k_1)$ using the estimated distribution of $X_{ji}$ and $Y_{ji}$ as $\mathrm{NB}(\hat{\mu}_{Xj}, (\hat{\mu}_{Xj})^{-1})$ and $\mathrm{NB}(\hat{\mu}_{Yj}, (\hat{\mu}_{Yj})^{-1})$, respectively, $j = 1, \ldots, d$; $i = 1, \ldots, 3$.

To implement the misspecified paired approach (as in BH-pair-mis and qvalue-pair-mis), we assumed that for each uninteresting feature $j$, $\{X_{ji}\}_{j=1}^3$ and $\{Y_{ji}\}_{j=1}^3$ follow the same Poisson distribution. We calculated the p-value of feature $j$ from a two-sample Poisson test for the null hypothesis $H_0 : \mu_{Xj} = \mu_{Yj}$ against the alternative hypothesis $H_1 : \mu_{Xj} > \mu_{Yj}$ using function `poisson.test` in R package `stats`.

To implement the 2as1 paired approach (as in BH-pair-2as1 and qvalue-pair-2as1), we treated $\mathrm{NB}(\sum_{i=1}^3 Y_{ji}, (\sum_{i=1}^3 Y_{ji})^{-1})$ conditioning on the observed $\sum_{i=1}^3 Y_{ji}$ as the null distribution of $\sum_{i=1}^3 X_{ji}$. Then we calculated the p-value of feature $j = 1, \ldots, d$ as the right tail probability of $\sum_{i=1}^3 X_{ji}$ in $\mathrm{NB}(\sum_{i=1}^3 Y_{ji}, (\sum_{i=1}^3 Y_{ji})^{-1})$.

#### S4.4 10vs10 enrichment analysis

We simulated data without outliers under heterogeneous background scenario and three distributional families—a total of $3$ settings. In each setting, we generated $d = 10{,}000$ features, among which $10\%$ are interesting (with $\mu_{Xj} > \mu_{Yj}$) and the rest are uninteresting (with $\mu_{Xj} = \mu_{Yj}$).

The data generation under the Gaussian, Poisson, and negative binomial distributions is the same as in the 3vs3 enrichment analysis (Section S4.3) except that we set the number of replicates to $10$ under each condition, and we did not generate outliers.

The correct paired approaches in BH-pair-parametric are the same as the corresponding BH-pair-correct in the 3vs3 enrichment analysis (Section S4.3) except that, under the negative binomial distribution, the test statistic $T_j$ and its null distribution should have the number of replicates changed from $3$ to $10$. The misspecified and 2as1 paired approaches (BH-pair-mis and BH-pair-2as1) are also the same as the corresponding approaches in the 3vs3 enrichment analysis (Section S4.3).

To implement the non-parametric paired approaches, we calculated the p-value of feature $j$ from the one-sided two-sample Wilcoxon rank-sum test (using `R` function `wilcox.test` in package `stats`) in BH-pair-Wilcoxon and from the one-sided two-sample permutation test (using `R` function `oneway_test` in package `coin`) in BH-pair-permutation.

#### S4.5 2vs1 differential analysis

We simulated data with $d = 10{,}000$ features under two background scenarios and three distributional families—a total of $6$ settings. In each setting, we set $10\%$ features as "up-regulated" with $\mu_{Xj} > \mu_{Yj}$ and another $10\%$ features as "down-regulated" with $\mu_{Xj} < \mu_{Yj}$.

###### Gaussian distribution

We simulated data from Gaussian using the following procedure:

- Under the homogeneous background scenario, we set $\mu_{Yj} = 0$ for all $d$ features. For uninteresting features, we set $\mu_{Xj} = \mu_{Yj} = 0$ for $j \in \mathcal{N}$. For up-regulated features, we generated $\mu_{Xj}$ i.i.d. from $N(5, 1)$. For down-regulated features, generated $\mu_{Xj}$ i.i.d. from $N(-5, 1)$ .
- Under the heterogeneous background scenario, we generated $\{\mu_{Yj}\}_{j=1}^{d}$ i.i.d. from $N(0, 2^2)$. For uninteresting features, we set $\mu_{Xj} = \mu_{Yj}$ for $j \in \mathcal{N}$. For up-regulated features, we generated $\mu_{Xj}$ i.i.d. from $N(5, 1)$. For down-regulated features, generated $\mu_{Xj}$ i.i.d. from $N(-5, 1)$ .
- We independently generated $X_{j1}$ and $X_{j2}$ from $N(\mu_{Xj}, 1)$ and $Y_{j1}$ from $N(\mu_{Yj}, 1)$, $j = 1, \ldots, d$.

To implement the misspecified paired approach (as in BH-pair-mis and qvalue-pair-mis), we assumed that the null distribution of $\frac{1}{2}(X_{j1} + X_{j2}) - Y_{j1}$, $j = 1, \ldots, d$, is $N(0, \hat{\sigma}^2)$, where

$$\hat{\sigma}^2 = \frac{1}{2(2d-1)} \sum_{j=1}^{d} \sum_{i=1}^{2} \left( X_{ji} - \frac{1}{2d} \sum_{j=1}^{d} \sum_{i=1}^{2} X_{ji} \right)^2 + \frac{1}{d-1} \sum_{j=1}^{d} \left( Y_{j1} - \frac{1}{d} \sum_{j=1}^{d} Y_{j1} \right)^2 .$$

This is a misspecified model assuming that $\mu_{Xj}$'s are all equal and so are $\mu_{Yj}$'s. Then we computed the p-value of feature $j = 1, \ldots, d$ as the two-sided tail probability of $\frac{1}{2}(X_{j1} + X_{j2}) - Y_{j1}$ in $N(0, \hat{\sigma}^2)$, i.e., $2 \cdot \min\left(1 - \Phi\left(\frac{\frac{1}{2}(X_{j1}+X_{j2})-Y_{j1}}{\hat{\sigma}}\right), \Phi\left(\frac{\frac{1}{2}(X_{j1}+X_{j2})-Y_{j1}}{\hat{\sigma}}\right)\right)$, where $\Phi$ is the cumulative distribution function of $N(0, 1)$.

To implement the 2as1 paired approach (as in BH-pair-2as1 and qvalue-pair-2as1), we treated $N(Y_{j1}, 1)$ conditioning on the observed $Y_{j1}$ as the null distribution of $X_{j1}$. Then we calculated the p-value of feature $j = 1, \ldots, d$ as the two-sided tail probability of $\frac{1}{2}(X_{j1} + X_{j2})$ in $N(Y_{j1}, 1/2)$, i.e., $2 \cdot \min\left(1 - \Phi\left(\frac{\frac{1}{2}(X_{j1}+X_{j2})-Y_{j1}}{1/\sqrt{2}}\right), \Phi\left(\frac{\frac{1}{2}(X_{j1}+X_{j2})-Y_{j1}}{1/\sqrt{2}}\right)\right)$.

###### Poisson distribution

We simulated data from Poisson using the following procedure:

- Under the homogeneous background scenario, we set $\mu_{Yj} = 20$ for all $d$ features. For uninteresting features, we set $\mu_{Xj} = \mu_{Yj} = 20$ for $j \in \mathcal{N}$. For up-regulated features, we generated $\mu_{Xj}$ i.i.d. from $\text{Pois}(60)$. For down-regulated features, we generated $\mu_{Xj}$ i.i.d. from $\text{Pois}(5)$ .
- Under the heterogeneous background scenario, we generated $\{\mu_{Yj}\}_{j=1}^{d}$ i.i.d. from $\text{Pois}(20)$. For uninteresting features, we set $\mu_{Xj} = \mu_{Yj}$ for $j \in \mathcal{N}$. For up-regulated features, we generated $\mu_{Xj}$ i.i.d. from $\text{Pois}(60)$. For down-regulated features, we generated $\mu_{Xj}$ i.i.d. from $\text{Pois}(5)$ .
- We independently generated $X_{j1}$ and $X_{j2}$ from $\text{Pois}(\mu_{Xj})$ and $Y_{j1}$ from $\text{Pois}(\mu_{Yj})$, $j = 1, \ldots, d$.

To implement the misspecified paired approach (as in BH-pair-mis and qvalue-pair-mis), we first defined a log-transformation $f(x) = \log(x + 0.01)$, which we applied to $X_{j1}$ and $Y_{j1}$, $j = 1, \ldots, d$. We assumed that the null distribution of $f(X_{j1}) + f(X_{j2}) - 2f(Y_{j1})$, $j = 1, \dots, d$ is $N(0, \hat{\sigma}^2)$, where

$$\hat{\sigma}^2 = \frac{6}{d-1} \sum_{j=1}^{d} \left( f(Y_{j1}) - \frac{1}{d} \sum_{j=1}^{d} f(Y_{j1}) \right)^2 .$$

This model misspecifies the Poisson distribution as the log-normal distribution. Then we computed the p-value of feature $j = 1, \dots, d$ as the two-sided tail probability of $f(X_{j1}) + f(X_{j2}) - 2f(Y_{j1})$ in $N(0, \hat{\sigma}^2)$, i.e., $2 \cdot \min \left( 1 - \Phi\left( \frac{f(X_{j1})+f(X_{j2})-2f(Y_{j1})}{\hat{\sigma}} \right), \Phi\left( \frac{f(X_{j1})+f(X_{j2})-2f(Y_{j1})}{\hat{\sigma}} \right) \right)$, where $\Phi$ is the cumulative distribution function of $N(0,1)$.

To implement the 2as1 paired approach (as in BH-pair-2as1 and qvalue-pair-2as1), we assumed that for each uninteresting feature $j$, $X_{j1}$ and $X_{j2}$ independently follow $\mathrm{Pois}(Y_{j1})$ conditioning on the observed $Y_{j1}$. Then we calculated the p-value of feature $j = 1, \dots, d$ by performing a one-sample Poisson test using the R function `poisson.test` for the null hypothesis $H_0 : \mu_{X_j} = Y_{j1}$ against the alternative hypothesis $H_1 : \mu_{X_j} \neq Y_{j1}$.

###### Negative binomial distribution

We simulated data from negative binomial using the following procedure:

- Under the homogeneous background scenario, we set $\mu_{Yj} = 30$ for all $d$ features. For uninteresting features, we set $\mu_{Xj} = \mu_{Yj} = 30$ for $j \in \mathcal{N}$. For up-regulated features, we generated $\mu_{Xj}$ i.i.d. from $\mathrm{NB}(70, 70^{-1})$. For down-regulated features, we generated $\mu_{Xj}$ i.i.d. from $\mathrm{NB}(7, 7^{-1})$.

- Under the heterogeneous background scenario, we generated $\{\mu_{Yj}\}_{j=1}^{d}$ i.i.d. from $\mathrm{NB}(30, 30^{-1})$. For uninteresting features, we set $\mu_{Xj} = \mu_{Yj}$ for $j \in \mathcal{N}$. For up-regulated features, we generated $\mu_{Xj}$ i.i.d. from $\mathrm{NB}(70, 70^{-1})$. For down-regulated features, we generated $\mu_{Xj}$ i.i.d. from $\mathrm{NB}(7, 7^{-1})$.

- We independently generated $X_{j1}$ and $X_{j2}$ from $\mathrm{NB}(\mu_{Xj}, \mu_{Xj}^{-1})$ and $Y_{j1}$ from $\mathrm{NB}(\mu_{Yj}, \mu_{Yj}^{-1})$, $j = 1, \dots, d$.

To implement the misspecified paired approach (as in BH-pair-mis and qvalue-pair-mis), we assumed that for each uninteresting feature $j$, $X_{j1}$, $X_{j2}$, and $Y_{j1}$ follow the same Poisson distribution. We calculated the p-value of feature $j$ from a two-sample Poisson test for the null hypothesis $H_0 : \mu_{Xj} = \mu_{Yj}$ against the alternative hypothesis $H_1 : \mu_{Xj} \neq \mu_{Yj}$ using the function `poisson.test` in R package `stats`.

To implement the 2as1 paired approach (as in BH-pair-2as1 and qvalue-pair-2as1), we treated $\mathrm{NB}(2Y_{j1}, (2Y_{j1})^{-1})$ conditioning on the observed $Y_{j1}$ as the null distribution of $X_{j1}+X_{j2}$. Then we calculated the p-value of feature $j = 1, \dots, d$ as the two-sided tail probability of $X_{j1}+X_{j2}$ in $\mathrm{NB}(2Y_{j1}, (2Y_{j1})^{-1})$, i.e., twice the smaller of the left-tail and right-tail probabilities.

#### S4.6 3vs3 differential analysis

We simulated data with or without outliers under two background scenarios and three distributional families—a total of $12$ settings. In each setting, we generated $d = 10{,}000$ features, among which $10\%$ features were "up-regulated features" with $\mu_{Xj} > \mu_{Yj}$ and another $10\%$ were "down-regulated features" with $\mu_{Xj} < \mu_{Yj}$.

Under the settings with outliers, we generated $\{O_{ji}^X : j = 1, \dots, d; i = 1, \dots, 3\}$ and $\{O_{ji}^Y : j = 1, \dots, d; i = 1, \dots, 3\}$ i.i.d. from $\mathrm{Bernoulli}(0.1)$, where $O_{ji}^X = 1$ or $O_{ji}^Y = 1$ indicates $X_{ji}$ or $Y_{ji}$ is an outlier, respectively. Under settings without outliers, $O_{ji}^X = O_{ij}^Y = 0$ for all $j = 1, \dots, d; i = 1, \dots, 3$.

**Gaussian distribution**

- Under the homogeneous background scenario, we set $\mu_{Yj} = 0$ for all $d$ features. For uninteresting features, we set $\mu_{Xj} = \mu_{Yj} = 0$ for $j \in \mathcal{N}$. For up-regulated features, we generated $\mu_{Xj}$ i.i.d. from $N(5, 1)$. For down-regulated features, generated $\mu_{Xj}$ i.i.d. from $N(-5, 1)$.

- Under the heterogeneous background scenario, we generated $\{\mu_{Yj}\}_{j=1}^{d}$ i.i.d. from $N(0, 2^2)$. For uninteresting features, we set $\mu_{Xj} = \mu_{Yj}$ for $j \in \mathcal{N}$. For up-regulated features, we generated $\mu_{Xj}$ i.i.d. from $N(5, 1)$. For down-regulated features, generated $\mu_{Xj}$ i.i.d. from $N(-5, 1)$.

- We independently generated $X_{ji}$ from $N(\mu_{Xj}, 1)$ if $O_{ji}^{X} = 0$ or from the top $1\%$ percentile of $N(\mu_{Xj}, 1)$ if $O_{ji}^{X} = 1$, $j = 1, \ldots, d$; $i = 1, \ldots, 3$. Similarly, we independently generated $Y_{ji}$ from $N(\mu_{Yj}, 1)$ if $O_{ji}^{Y} = 0$ or from the top $1\%$ percentile of $N(\mu_{Yj}, 1)$ if $O_{ji}^{Y} = 1$, $j = 1, \ldots, d$; $i = 1, \ldots, 3$.

- For the results in Additional File 1: Fig. S14, under the heterogeneous background scenario, we generated $\{\mu_{Yj}\}_{j=1}^{d}$ i.i.d. from $N(0, 2^2)$, and $\{s_j\}_{j=1}^{d}$ i.i.d. from a uniform distribution $U(0.5, 2)$. For uninteresting features, we set $\mu_{Xj} = \mu_{Yj}$ for $j \in \mathcal{N}$. For up-regulated features, we generated $\mu_{Xj}$ i.i.d. from $N(5, 1)$. For down-regulated features, generated $\mu_{Xj}$ i.i.d. from $N(-5, 1)$. We then independently generated $X_{ji}$ from $N(\mu_{Xj}, s_j^2)$ if $O_{ji}^{X} = 0$ or from the top $1\%$ percentile of $N(\mu_{Xj}, s_j^2)$ if $O_{ji}^{X} = 1$, $j = 1, \ldots, d$; $i = 1, \ldots, 3$. Similarly, we independently generated $Y_{ji}$ from $N(\mu_{Yj}, s_j^2)$ if $O_{ji}^{Y} = 0$ or from the top $1\%$ percentile of $N(\mu_{Yj}, s_j^2)$ if $O_{ji}^{Y} = 1$, $j = 1, \ldots, d$; $i = 1, \ldots, 3$.

To implement the correct paired approach (as in BH-pair-correct and qvalue-pair-correct), we calculated the p-value of feature $j$ from a two-sample t-test with equal variance for the null hypothesis $H_0 : \mu_{Xj} = \mu_{Yj}$ against the alternative hypothesis $H_1 : \mu_{Xj} \neq \mu_{Yj}$.

To implement the misspecified paired approach (as in BH-pair-mis and qvalue-pair-mis), we calculated the p-value of feature $j$ from a two-sample t-test with unequal variance for the null hypothesis $H_0 : \mu_{X_j} = \mu_{Y_j}$ against the alternative hypothesis $H_1 : \mu_{X_j} \neq \mu_{Y_j}$.

To implement the 2as1 paired approach (as in BH-pair-2as1 and qvalue-pair-2as1), we treated $N(\bar{Y}_j, 1)$ conditioning on observed $\bar{Y}_j$ as the null distribution of $X_{ji}$, $i = 1, \ldots, 3$. We calculated the p-value of feature $j$ using a one-sample t-test for the null hypothesis $H_0 : \mu_{X_j} = \bar{Y}_j$ against the alternative hypothesis $H_1 : \mu_{X_j} \neq \bar{Y}_j$.

**Poisson distribution**

We simulated data from Poisson using the following procedure:

- Under the homogeneous background scenario, we set $\mu_{Yj} = 20$ for all $d$ features. For uninteresting features, we set $\mu_{Xj} = \mu_{Yj} = 20$ for $j \in \mathcal{N}$. For up-regulated features, we generated $\mu_{Xj}$ i.i.d. from $\text{Pois}(40)$. For down-regulated features, we generated $\mu_{Xj}$ i.i.d. from $\text{Pois}(5)$.

- Under the heterogeneous background scenario, we generated $\{\mu_{Yj}\}_{j=1}^{d}$ i.i.d. from $\text{Pois}(20)$. For uninteresting features, we set $\mu_{Xj} = \mu_{Yj}$ for $j \in \mathcal{N}$. For up-regulated features, we generated $\mu_{Xj}$ i.i.d. from $\text{Pois}(40)$. For down-regulated features, we generated $\mu_{Xj}$ i.i.d. from $\text{Pois}(5)$.

- We independently generated $X_{ji}$ from $\text{Pois}(\mu_{Xj})$ if $O_{ji}^{X} = 0$ or from the top $1\%$ percentile of $\text{Pois}(\mu_{Xj})$ if $O_{ji}^{X} = 1$, $j = 1, \ldots, d$; $i = 1, \ldots, 3$. Similarly, we independently generated $Y_{ji}$ from $\text{Pois}(\mu_{Yj})$ if $O_{ji}^{Y} = 0$ or from the top $1\%$ percentile of $\text{Pois}(\mu_{Yj})$ if $O_{ji}^{Y} = 1$, $j = 1, \ldots, d$; $i = 1, \ldots, 3$.

To implement the correct paired approach (as in BH-pair-correct and qvalue-pair-correct), we calculated the p-value of feature $j$ by performing a two-sample Poisson test for the null hypothesis $H_0$ : $\mu_{Xj} = \mu_{Yj}$ against the alternative hypothesis $H_1 : \mu_{Xj} \neq \mu_{Yj}$ using function `poisson.test` in R package `stats`.

To implement the misspecified paired approach (as in BH-pair-mis and qvalue-pair-mis), we first defined a log-transformation $f(x) = \log(x + 0.01)$, which we applied to $X_{ji}$ and $Y_{ji}$, $j = 1, \ldots, d$; $i = 1, \ldots, 3$. We assumed that for each uninteresting feature $j$, $\{f(X_{ji})\}_{i=1}^3$ and $\{f(Y_{ji})\}_{i=1}^3$ follow Gaussian distributions with mean $\mu_{f(Xj)}$ and $\mu_{f(Yj)}$, respectively. Then we computed the p-value of feature $j$ using a two-sample equal variance t-test for the null hypothesis $H_0 : \mu_{f(Xj)} = \mu_{f(Yj)}$ against the alternative hypothesis $H_1 : \mu_{f(Xj)} \neq \mu_{f(Yj)}$.

To implement the 2as1 paired approach (as in BH-pair-2as1 and qvalue-pair-2as1), we assumed that for each uninteresting feature $j$, $\{X_{ji}\}_{i=1}^3$ follow $\text{Pois}(\bar{Y}_j)$ conditioning on the observed $\bar{Y}_j$. We calculated the p-value of feature $j$ by performing a one-sample Poisson test for the null hypothesis $H_0 : \mu_{Xj} = \bar{Y}_j$ against the alternative hypothesis $H_1 : \mu_{Xj} \neq \bar{Y}_j$ using the function `poisson.test` in R package `stats`.

##### Negative binomial distribution

We simulated data from negative binomial using the following procedure:

- Under the homogeneous background scenario, we set $\mu_{Yj} = 30$ for all $d$ features. For uninteresting features, we set $\mu_{Xj} = \mu_{Yj} = 30$ for $j \in \mathcal{N}$. For up-regulated features, we generated $\mu_{Xj}$ i.i.d. from $\text{NB}(70, 70^{-1})$. For down-regulated features, we generated $\mu_{Xj}$ i.i.d. from $\text{NB}(7, 7^{-1})$ .

- Under the heterogeneous background scenario, we generated $\{\mu_{Yj}\}_{j=1}^d$ i.i.d. from $\text{NB}(30, 30^{-1})$. For uninteresting features, we set $\mu_{Xj} = \mu_{Yj}$ for $j \in \mathcal{N}$. For up-regulated features, we generated $\mu_{Xj}$ i.i.d. from $\text{NB}(70, 70^{-1})$. For down-regulated features, we generated $\mu_{Xj}$ i.i.d. from $\text{NB}(7, 7^{-1})$ .

- We independently generated $X_{ji}$ from $\text{NB}(\mu_{Xj}, \mu_{Xj}^{-1})$ if $O_{ji}^X = 0$ or from the top $1\%$ percentile of $\text{NB}(\mu_{Xj}, \mu_{Xj}^{-1})$ if $O_{ji}^X = 1$; $j = 1, \ldots, d$, $i = 1, \ldots, 3$. Similarly, we independently generated $Y_{ji}$ from $\text{NB}(\mu_{Yj}, \mu_{Yj}^{-1})$ if $O_{ji}^Y = 0$ or from the top $1\%$ percentile of $\text{NB}(\mu_{Yj}, \mu_{Yj}^{-1})$ if $O_{ji}^Y = 1$, $j = 1, \ldots, d$; $i = 1, \ldots, 3$.

To implement the correct paired approach with unknown dispersion (as in BH-pair-correct and qvalue-pair-correct), we performed a two-sample negative binomial test for the null hypothesis $H_0$ : $\mu_{Xj} = \mu_{Yj}$ against the alternative hypothesis $H_1 : \mu_{Xj} \neq \mu_{Yj}$ using the coefficient from the negative binomial regression as the test statistic. Specifically, for each feature $j$ we performed a negative binomial regression by treating the condition labels as a categorical covariate and feature $j$'s measurements as the response. We implemented this regression analysis using function `glm.nb` in R package `MASS` and extracted the p-value of the coefficient as the p-value of feature $j$. The dispersion parameter was not pre-specified but estimated by `glm.nb`.

To implement the correct paired approach with known dispersion, we performed a similar negative binomial regression but with the pre-specified dispersion parameter $30^{-1}$ for each feature $j$. Then we computed the feature's p-value as the p-value of the coefficient of the condition covariate. We implemented this regression analysis using function `glm` in R package `stats`.

To implement the misspecified paired approach (as in BH-pair-mis and qvalue-pair-mis), we assumed that for each uninteresting feature $j$, $\{X_{ji}\}_{j=1}^3$ and $\{Y_{ji}\}_{j=1}^3$ follow the same Poisson distribution. We calculated the p-value of feature $j$ from a two-sample Poisson test for the null hypothesis

$H_0 : \mu_{Xj} = \mu_{Yj}$ against the alternative hypothesis $H_1 : \mu_{Xj} \neq \mu_{Yj}$ using function `poisson.test` in R package `stats`.

To implement the 2as1 paired approach (as in BH-pair-2as1 and qvalue-pair-2as1), we first used function `glm.nb` in R package `MASS` to estimate $\hat{\mu}_{Yj}$ and $\hat{\theta}_{Yj}$ from $\{Y_{ji}\}_{j=1}^{3}$. Then we computed the p-value of feature $j$ by treating $\text{NB}(3\hat{\mu}_{Yj}, (3\hat{\theta}_{Yj})^{-1})$ as the null distribution of $\sum_{i=1}^{3} X_{ji}$ and calculated its two-sided tail probability, i.e., twice the smaller of the left-tail and right-tail probabilities.

### S5 Bioinformatic methods with FDR control functionality

#### S5.1 Peak calling methods for ChIP-seq data

**MACS2** MACS2 [10] uses sliding windows with a fixed length across the genome and identifies peaks by using a Poisson distribution to model the read counts within each window, which has one read count per replicate. Specifically, for each region (which is combined from sliding windows), MACS2 performs a one-sample Poisson test to calculate a p-value, where the null distribution is set to be Poisson with its parameter estimated from the background. By thresholding p-values, MACS2 identifies a set of candidate peaks. It also estimates for each candidate peak a q-value by swapping the experimental sample with the background (negative control) sample, and the q-values are used for FDR control. We used MACS2 software (version 2.2.6) with its default settings.

**HOMER** We used findPeaks, a program in HOMER [11], to perform peak calling on ChIP-seq data. The p-value calculation in findPeaks is similar to that in MACS2; that is, findPeaks also uses the Poisson distribution as the null distribution of read counts in a genomic region, and it also estimates the Poisson mean from the background sample. Then findPeaks identifies peaks by setting thresholds on p-values and fold-changes (the folder change of a region is defined as the observed read count under the experimental sample divided by the estimated Poisson mean from the the background sample). We used findPeaks version 3.1.9.2.

#### S5.2 SEQUEST for peptide identification from MS data

**SEQUEST** SEQUEST uses probability-based scoring to identify PSMs from mass-spectrometry data. We ran SEQUEST in Proteome Discoverer 2.3.0.523 (ThermoScientific) with the following settings: 10 ppm precursor tolerance; 0.6 Da fragment tolerance; static modifications: methylthio (C); dynamic modifications: deamination (NQ), oxidation (M). We ran Percolator [12] in conjunction with SEQUEST with the target/decoy selection mode set to "separate." For SEQUEST, for a range of target FDR thresholds ($q \in \{1\%, 2\%, \ldots, 10\%\}$), we identified the target PSMs with SEQUEST q-values no greater than $q$ as discoveries. To prepare the input for Clipper, we set peptide and protein FDRs to $100\%$ to obtain the entire lists of target PSMs and decoy PSMs with their SEQUEST q-values.

#### S5.3 Differentially expressed gene (DEG) methods for bulk RNA-seq data

**edgeR** edgeR models each gene's read counts by using a negative binomial regression, where the condition is incorporated as an indicator covariate, and the condition's coefficient represents the gene-wise differential expression effect [13]. We used `R` package `edgeR` version 3.30.0.

**DESeq2** DESeq2 uses a similar negative binomial regression as edgeR to model each gene's read counts under two conditions. DESeq2 differs from edgeR mainly in their estimation of the dispersion parameter in the negative binomial distribution [14]. We used `R` package `DESeq2` version 1.28.1.

##### S5.4 Differentially expressed gene (DEG) methods for scRNA-seq data

**MAST** MAST models each gene's log read counts (TPM) by using a two-part generalized regression model. Each gene's expression rate was modeled using logistic regression and, conditioning on a cell expressing the gene, the gene's expression level was modeled as Gaussian [15]. We used `R` package `MAST` version 1.14.0.

**Monocle3** Monocle3 uses a generalized linear model to model each gene's normalized expression value, with other information included as covariates (time, treatment, and so on) [16]. We used `R` package `monocle3` version 0.2.3.0.

##### S5.5 Differentially interacting chromatin regions (DIR) methods for Hi-C data

**MultiHiCcompare** MultiHiCcompare relies on a non-parametric method to jointly normalize multiple Hi-C interaction matrices [17]. It uses a generalized linear model to detect DIRs. MultiHiCcompare is an extension of the `HiCcompare` package [18]. We used `R` package `multiHiCcompare` version 1.6.0.

**diffHiC** diffHiC uses the statistical framework of the `edgeR` package to model biological variability and to test for significant differences between conditions [19]. We used `R` package `diffHiC` version 1.20.0.

**FIND** FIND uses a spatial Poisson process to detect chromosomal regions that display a significant difference between two regions' contact intensity and their neighbouring contact intensities [20]. We used `R` package `FIND` version 0.99.

#### S6 Benchmark data generation in omics data applications

##### S6.1 ChIP-seq data with synthetic spike-in peaks

We used two control samples (which we refer to as Control 1 and Control 2) from H3K4me3 ChIP-seq data in Chromosome 1 of the cell line GM12878 [21].

(i) We created two semi-synthetic experimental samples by adding synthetic true peaks to Control 1. To mimic real H3K4me3 ChIP-seq data, where peaks are located predominantly in promoter regions, we added synthetic true peaks to promoter regions annotated from Ensembl BioMart (Ensemble hg 19, regulation 104) [22]. Specifically, we randomly sampled $585$ genes' promoter regions from Chromosome 1. We then used ChIPulate to simulate reads from these promoter regions (for each simulation, extraction efficiency parameter and PCR efficiency parameter were randomly sampled from a uniform distribution between $0$ to $1$; binding energy parameters were randomly sampled from a uniform distribution between $0$ and $2$; sequencing depth parameter was set to $50$). Then we added the simulated reads to Control 1. We repeated this procedure for twice to obtain two semi-synthetic experimental samples (i.e., two replicates under the experimental condition).

(ii) We repeated Step (i) for $20$ times to generate $20$ sets of semi-synthetic experimental samples. For each set of experimental samples, we paired them with Control 2, which was treated as the background sample (i.e., one replicate under the background condition). Hence, we obtained $20$ semi-synthetic ChIP-seq datasets, each containing $585$ synthetic true peaks.

(iii) After applying a peak calling method to these $20$ semi-synthetic datasets, we evaluated the method's $20$ FDPs and $20$ empirical power, which were then averaged as the method's approximate FDR and power. In the evaluation, a called peak was a true positive if it overlapped with a synthetic true peak; otherwise, it was a false positive.

#### S6.2 Real MS benchmark data

We purchased the complex proteomics standard (CPS) (part number 400510) from Agilent (Agilent, Santa Clara, CA, USA). The CPS contains soluble proteins extracted from the archaeon *Pyrococcus furiosus* (*Pfu*), which has a complete protein database; that is, all proteins from *Pfu* were catalogued into its protein database with known protein sequences. We subjected the CPS to a shotgun proteomics analysis that generated mass spectra of *Pfu*.

To generate a benchmark dataset, we first generated a reference database by concatenating the Uniprot *Pyrococcus furiosus* (*Pfu*) database, the Uniprot Human database, and two contaminant databases: the CRAPome [23] and the contaminant database from MaxQuant [24]. During the process, we purified the reference database by first performing *in silico* digestion of *Pfu* proteins and then removing human proteins that contained *Pfu* peptides from the reference database. We then input the *Pfu* mass spectra (from the CPS) and the purified reference database into SEQUEST. We considered a target PSM as true if SEQUEST reported its protein as from *Pfu* or the two contaminants; otherwise (if from Human), we considered the target PSM as false. The *in silico* digestion was performed in Python using the `pyteomics.parser` function from `pyteomics` with the following settings: Trypsin digestion, two allowed missed cleavages, minimum peptide length of six [25, 26].

#### S6.3 Bulk RNA-seq data with synthetic spike-in DEGs

We generated four sets of realistic semi-synthetic data from two real RNA-seq datasets. The first one is a human monocyte RNA-seq dataset including $17$ samples of classical monocytes and $17$ samples of non-classical monocytes [27]. Each sample contains expression levels of $d = 52{,}376$ transcripts.

The second one is a yeast RNA-seq dataset including $48$ samples of a *snf2* knockout mutant cell line and $48$ samples of negative control (without the knockout) [28]. Each sample contains expression levels of $d = 7126$ genes. We preprocessed this dataset by removing low-quality replicates (replicates $6, 13, 25, 35$ from the knockout; replicates $21, 22, 25, 28, 34, 36$ from the control) identified by the original paper Gierliński et al. [28], leaving us with $44$ replicates under the knockout condition and $42$ replicates under the negative control.

Here we describe our **simulation strategy 1**. Given either the human monocyte dataset or the yeast dataset, we performed the following steps.

(i) We first performed normalization on all samples across two conditions using the edgeR normalization method trimmed mean of M-values (TMM) [29]. We denote the resulting normalized read count matrix of classical human monocytes or yeasts without the knockout by $\mathbf{X}^1$ and the normalized read count matrix of non-classical human monocytes or yeast with the knockout by $\mathbf{X}^2$, respectively. Following the convention in bioinformatics, the columns and rows of $\mathbf{X}^1$ and $\mathbf{X}^2$ represent biological samples and genes, respectively.

(ii) To define true DEGs, we first computed the fold change of gene $j$ by FC$_j = \left[(\bar{\mathbf{X}}_j^2 + 1)/(\bar{\mathbf{X}}_j^1 + 1)\right]$ for $j = 1, \ldots, d$, where $\mathbf{X}_j^1$ and $\mathbf{X}_j^2$ denote the $j$-th row vector of $\mathbf{X}^1$ and $\mathbf{X}^2$ respectively and $\bar{\cdot}$ denotes the average of elements in a vector. We added the pseudo-count of $1$ to avoid division by $0$. We defined true DEGs as those with $|\log_2$ FC$_j| \geq 4$ for the human monocyte dataset and with $|\log_2$ FC$_j| \geq 1.5$ for the yeast dataset, resulting $191$ true human DEGs (transcripts) and $152$ true yeast DEGs.

(iii) We generated semi-synthetic data with $3$ samples under both the experimental and background conditions, a typical design in bulk RNA-seq experiments. Specifically, if gene $j$ is a true DEG, we randomly sampled without replacement $3$ values from $\mathbf{X}_j^1$ as counts under the experimental condition, and another $3$ values from $\mathbf{X}_j^2$ as counts under the background condition. If gene $j$ is not a true DEG, we randomly sampled $6$ values without replacement from $(\mathbf{X}_j^1, \mathbf{X}_j^2)$ and randomly split them into $3$ and $3$ counts under two conditions. Doing so guaranteed that a non-DEG's read counts are i.i.d. regardless of condition.

(iv) We repeated Step (iii) for $100$ times to generate $100$ semi-synthetic datasets.

Next, we describe our **simulation strategy 2**. Let us now re-use notations $\mathbf{X}^1$ to denote the original read count matrix of classical human monocytes or yeast without the knockout, and $\mathbf{X}^2$ to denote the original read count matrix of non-classical human monocytes or yeast with the knockout. Both $\mathbf{X}^1$ and $\mathbf{X}^2$ have rows as genes or transcripts and columns as biological samples. Given either the human monocyte dataset or the yeast dataset, we performed the following steps.

(i) We first identified genes whose read counts are positive in all samples under both conditions and denote the number of such genes by $d_p$. Then from these identified genes, we randomly sampled without replacement $\min(d_p, 0.3d)$ genes as true DEGs. The remaining $d - \min(d_p, 0.3d)$ genes were considered true non-DEGs.

(ii) To generate fold changes of true DEGs, we first computed the fold change of gene $j$ by FC$_j = \left[(\bar{\mathbf{X}}_j^2 + 1)/(\bar{\mathbf{X}}_j^1 + 1)\right]$ for $j = 1, \ldots, d$, where $\mathbf{X}_j^1$ and $\mathbf{X}_j^2$ denote the $j$-th row vector of $\mathbf{X}^1$ and $\mathbf{X}^2$ respectively and $\bar{\cdot}$ denotes the average of elements in a vector. Let $\mathcal{W}$ denote $\{$FC$_j$ : FC$_j \geq 16, j = 1, \ldots, d\}$ for the human monocyte dataset and $\{$FC$_j$ : FC$_j \geq 1.5, j = 1, \ldots, d\}$ for the yeast dataset. We then sorted unique elements in $\mathcal{W}$ and denoted them by $w_{(1)} < \cdots < w_{(n_u)}$, where $n_u$ is the number of unique elements in $\mathcal{W}$. To generate a fold change of a true DEG, say gene $j$, we randomly generated an integer $v$ with equal probability from $\{1, \cdots, n_u - 1\}$ and a value $p$ from Uniform$(0, 1)$. Then we calculated the fold change as $R_j = w_{(v)} + p(w_{(v+1)} - w_{(v)})$. Using this approach, generated the fold changes independently for all true DEGs.

(iii) Next, we randomly sampled $6$ replicates without replacement from $\mathbf{X}^2$ and split them into two groups of $3$ replicates. We denote the resulting matrices as $\widetilde{\mathbf{X}}^1$ and $\widetilde{\mathbf{X}}^2$, whose $j$-th rows are denoted respectively by $\widetilde{\mathbf{X}}_j^1$ and $\widetilde{\mathbf{X}}_j^2$. If gene $j$ is a true DEG, we generated $U_j$ from Bernoulli$(1/2)$. Then we set gene $j$'s expression levels under the two conditions to $R_j\widetilde{\mathbf{X}}_j^1$ and $\widetilde{\mathbf{X}}_j^2$ if $U_j = 0$ or $\widetilde{\mathbf{X}}_j^1$ and $R_j\widetilde{\mathbf{X}}_j^2$ if $U_j = 1$. If gene $j$ is not a true DEG, its expression levels under the two conditions would remain unchanged, i.e., $\widetilde{\mathbf{X}}_j^1$ and $\widetilde{\mathbf{X}}_j^2$. Such data generation strategy has no guarantee of i.i.d. read counts for non-DEGs if the samples in $\mathbf{X}^2$ have batch effects.

(iv) We repeated Step (iii) for $100$ times to generate $100$ semi-synthetic datasets.

The human monocyte RNA-seq dataset is available in the NCBI Sequence Read Archive (SRA) under accession number SRP082682 (`https://www.ncbi.nlm.nih.gov/Traces/study/?acc=srp082682`). The yeast RNA-seq data is available in the European Nucleotide Archive (ENA) archive with project ID PRJEB5348 (`https://www.ebi.ac.uk/ena/browser/view/PRJEB5348`).

##### S6.4 Single-cell RNA-seq data with synthetic spike-in DEGs

We used scDesign2, a flexible probabilistic simulator to generate realistic scRNA-seq count data with gene correlations captured [30]. Using scDesign2, we generated two sets of semi-synthetic data from two peripheral blood mononuclear cell (PBMC) real datasets [31]: one generated using the 10x Genomics protocol [32] and the other using Drop-seq [33]. Each semi-synthetic dataset contains two types of cells: CD4+ T cells, and cytotoxic T cells, which we treated as two conditions. Starting with the real data generated using either 10x Genomics or Drop-seq, we used the following steps to generate semi-synthetic scRNA-seq data.

(i) First, we fit the real data count matrices using `R` function `fit_model_scDesign2` for each cell type by specifying the underlying distribution of each gene as negative binomial. Denote the resulting marginal distributions of gene $j$ as $NB(\hat{\mu}_{j1}, \hat{\theta}_{j1})$ for CD4+ T cells and $NB(\hat{\mu}_{j2}, \hat{\theta}_{j2})$ for cytotoxic T cells, $j = 1, \ldots, d$. The gene-gene correlations with each cell type were fitted using a copula model.

(ii) Let $\mathbf{X}^{\text{cd4}}$ and $\mathbf{X}^{\text{cyto}}$ denote the read count matrices of CD4+ T cells and cytotoxic T cells. To define true DEGs, we first computed the log fold change of gene $j$ by logFC$_j = \log_2\left[(\bar{\mathbf{X}}_j^{\text{cd4}} + 1)/(\bar{\mathbf{X}}_j^{\text{cyto}} + 1)\right]$ for $j = 1, \ldots, d$, where $\mathbf{X}_j^{\text{cd4}}$ and $\mathbf{X}_j^{\text{cyto}}$ denote the $j$-th row vector of $\mathbf{X}^{\text{cd4}}$ and $\mathbf{X}^{\text{cyto}}$ respectively and $\bar{\cdot}$ denotes the average of elements in a vector. We then selected $1000$ genes with the largest absolute fold changes as true DEGs and kept the remaining ones as true non-DEGs.

(iii) We simulated the semi-synthetic datasets using `R` function `simulate_count_scDesign2`. Specifically, we set the number of synthetic cells generated by scDesign2 equal to the number of real cells for each cell type. If a gene $j$ is a true DEG, we specify its marginal distributions under the two conditions as $NB(\hat{\mu}_{j1}, \hat{\theta}_{j1})$ and $NB(\hat{\mu}_{j2}, \hat{\theta}_{j2})$ respectively. If a gene $j$ is a true non-DEG, we specify its marginal distribution under both conditions as $NB((\hat{\mu}_{j1} + \hat{\mu}_{j2})/2, (\hat{\theta}_{j1} + \hat{\theta}_{j2})/2)$. We used the fitted copula models from the two cell types to generate genes' (correlated) expression read counts.

(iv) We repeated Step (iii) for $200$ times to generate $200$ semi-synthetic datasets.

Both `fit_model_scDesign2` and `simulate_count_scDesign2` come from `R` package `scDesign2` [30]. The 10x Genomic PBMC dataset and the Drop-seq PBMC dataset are available from the Gene Expression Omnibus (GEO) with accession number GSE132044 (https://www.ncbi.nlm.nih.gov/geo/query/acc.cgi?acc=GSE132044) and the Single Cell Portal with accession numbers SCP424 (https://singlecell.broadinstitute.org/single_cell/study/SCP424/single-cell-comparison-pbmc-data).

##### S6.5 Hi-C data with synthetic spike-in DIRs

The real Hi-C interaction matrix contains the pairwise contact intensities of $250$ binned genomic regions in Chromosome 1. It is from the cell line GM12878 and available in the NCBI Gene Expression Omnibus (GEO) under accession number GSE63525. We denote the real interaction matrix as $\mathbf{X}^{\text{real}}$. Because $\mathbf{X}^{\text{real}}$ is symmetric, we only focus on its upper triangular part.

(i) Among the $(250 \times 250 - 250)/2 = 31{,}125$ upper triangular entries (i.e., region pairs), we selected $404$ entries as true up-regulated DIRs, and $550$ entries as true down-regulated DIRs (Fig. S29).

(ii) Next, for the $(i, j)$-th entry, we generated a log fold change, denoted by $f_{ij}$, between the two conditions as follows. We simulated $f_{ij}$ from truncated Normal$(100/|i-j|, 0.5^2)$ with support $[0.05, \infty)$ if the $(i, j)$-th entry is up-regulated, or from truncated Normal$(-100/|i-j|, 0.5^2)$ with support $(-\infty, -0.05]$ if the $(i,j)$-th entry is down-regulated; if the $(i,j)$-th entry is not differential, we set $f_{ij} = 0$.

(iii) Then we specify the mean measurement of the $(i,j)$-th entry under the two conditions as $\mu_{Xij} = [\mathbf{X}^{\text{real}}]_{ij}$ and $\mu_{Yij} = [\mathbf{X}^{\text{real}}]_{ij} \cdot e^{f_{ij}}$, respectively.

(iv) We generated synthetic read counts of the $(i,j)$-th entry from $\text{NB}(\mu_{Xij}, 1000^{-1})$ and $\text{NB}(\mu_{Yij}, 1000^{-1})$ respectively under the two conditions.

(v) We repeated Step (iv) for $200$ times to generate $200$ semi-synthetic datasets.

### S7 Implementation of Clipper in omics data applications

Below we briefly introduce the implementation of Clipper in the four omics data applications. All the results were obtained by running using `R` package `Clipper` (see package vignette for details: `https://github.com/JSB-UCLA/Clipper/blob/master/vignettes/Clipper.pdf`).

#### S7.1 Peak calling from ChIP-seq data

(i) We consider each genomic location, i.e., a base pair, as a feature and each ChIP-seq sample as a replicate under the experimental or background condition. Then we consider the read count of each location in each sample as the corresponding feature's measurement. Doing so, we summarized ChIP-seq data into a $d \times (m+n)$ matrix, where $d$ is the number of locations, and $m$ and $n$ are the numbers of experimental and control samples, respectively. We then applied Clipper to perform an enrichment analysis to obtain the contrast score $C_j$ for each location $j$. In our study, $m = n = 1$, so the default Clipper implementation is Clipper-minus-BC.

(ii) For any target FDR threshold $q$, Clipper gives a cutoff $T_q$ on contrast scores.

(iii) We then used existing peak calling methods, e.g., MACS2 and HOMER, to call candidate peaks with the least stringent q-value cutoff. For example, when we used MACS2, we set the q-value cutoff as 1.

(iv) We computed the contrast score of each candidate peak as the median of the contrast scores of all the locations within.

(v) The candidate peaks with contrast scores greater than or equal to $T_q$ are called discoveries.

It is known that uninteresting regions tend to have larger read counts in the control sample than in the experimental (ChIP) sample, making them more likely to have negative contrast scores than positive ones. However, this phenomenon does not violate Clipper's theoretical assumption (Lemma 1(a) in Section S2).

#### S7.2 Peptide identification from mass spectrometry data

(i) We consider each mass spectrum as a feature and its target/decoy PSM as a replicate under the experimental/background condition respectively. Then we consider $-\log_{10}(\text{q-value} + 0.01)$ as the measurement of each PSM, where the q-value is output by SEQUEST. Doing so, we summarized the SEQUEST output into a $d \times (m+n)$ matrix, where $d$ is the number of mass spectra, and $m$ and $n$ are the numbers of experimental and control samples, respectively. We then applied

Clipper to perform an enrichment analysis to obtain a contrast score $C_j$ for each mass spectrum $j$. If the mass spectrum has no decoy or background measurement, we set $C_j = 0$. In our study, $m = n = 1$, so the default Clipper implementation is Clipper-minus-BC.

(ii) For any target FDR threshold $q$, Clipper gives a cutoff $T_q$ on contrast scores.

(iii) The target PSMs whose mass spectra have contrast scores greater than or equal to $T_q$ are called discoveries.

##### S7.3 DEG identification from bulk RNA-seq data

(i) We consider each gene as a feature and the class label—classical and non-classical human monocytes—as the two conditions. We first performed the TMM normalization method [29]. Then we consider $\log_2$-transformed read counts with a pseudocount $1$ as measurements. Doing so, we summarized the gene expression matrix into a $d \times (m+n)$ matrix, where $d$ is the number of genes, and $m$ and $n$ are the numbers of samples under the two conditions, respectively. We then applied Clipper to perform a differential analysis to obtain a contrast score $C_j$ for each gene. In our study, $m = n = 3$, so the default Clipper implementation is Clipper-max-GZ with $h = 9$, the maximum number of permutations when we have three replicates under both conditions.

(ii) For any target FDR threshold $q$, Clipper gives a cutoff $T_q$ on contrast scores.

(iii) The genes with contrast scores greater than or equal to $T_q$ are called discoveries.

##### S7.4 DEG identification from scRNA-seq data

(i) We consider each gene as a feature and the two cell types—CD4+ T cells and cytotoxic T cells—as the two conditions. We first performed the TMM normalization [29]. Then we consider $\log_2$-transformed read counts with a pseudocount $1$ as measurements. Doing so, we summarized the gene expression matrix into a $d \times (m + n)$ matrix, where $d$ is the number of genes, and $m$ and $n$ are the numbers of samples under the two conditions, respectively. We then applied Clipper to perform differential analysis to obtain a contrast score $C_j$ for each gene $j$. In our study, $m = 1172$, $n = 789$ for Drop-seq dataset and $m = 963$, $n = 694$ for 10x Genomics dataset. The default Clipper implementation is Clipper-max-GZ with $h = 1$, the default number of permutations.

(ii) For any target FDR threshold $q$, Clipper gives a cutoff $T_q$ on contrast scores.

(iii) The genes with contrast scores greater than or equal to $T_q$ are called discoveries.

##### S7.5 DIR identification from Hi-C data

(i) We consider each pair of genomic regions as a feature and manually created two conditions. Then we consider $\log$-transformed read counts as measurements. Doing so, we summarized the gene expression matrix into a $d \times (m + n)$ matrix, where $d$ is the total pairs of genomic regions, and $m$ and $n$ are the numbers of samples under the two conditions, respectively. We then applied Clipper to perform a differential analysis to obtain a contrast score $C_j$ for each pair of genomic regions. In our study, $m = n = 2$, so the default Clipper implementation is Clipper-max-GZ with $h = 1$.

(ii) For any target FDR threshold $q$, Clipper gives a cutoff $T_q$ on contrast scores.

(iii) The pairs of genomic regions with contrast scores greater than or equal to $T_q$ are called discoveries.

### S8 Proofs

#### S8.1 Proof of Theorem 1

We first prove Theorem 1, which relies on Lemmas 1 and 2. Here we only include the proof of Lemma 1 and defer the proof of Lemma 2 to Section S8.3.

**Proof 1 (Proof of Lemma 1)** *Here we prove that Lemma 1 holds when $C_j$ is constructed using (S10); the proof is similar when $C_j$ is constructed using (S11).*

*When input data satisfy (S6) and (S7) and $m = n$, properties (a) and (b) can be derived directly. To prove property (c), it suffices to prove that for any $j \in \mathcal{N}$ with $C_j \neq 0$, $S_j$ is independent of $|C_j|$.*

*Note that $\bar{X}_j$ and $\bar{Y}_j$ are i.i.d for $j \in \mathcal{N}$ when $m = n$. Hence for any measurable set $\mathcal{A} \subset [0, +\infty)$,*

$$\mathbb{P}\left(S_j = 1, |C_j| \in \mathcal{A}\right) = \mathbb{P}\left(t^{\text{minus}}(\boldsymbol{X}_j, \boldsymbol{Y}_j) \in \mathcal{A}\right) = \mathbb{P}\left(t^{\text{minus}}(\boldsymbol{Y}_j, \boldsymbol{X}_j) \in \mathcal{A}\right) = \mathbb{P}\left(S_j = -1, |C_j| \in \mathcal{A}\right).$$

*The first equality holds because $t^{\text{minus}}(\boldsymbol{X}_j, \boldsymbol{Y}_j) = C_j = |C_j|$ when $S_j = 1$. The second equality holds because $t^{\text{minus}}(\boldsymbol{X}_j, \boldsymbol{Y}_j)$ and $t^{\text{minus}}(\boldsymbol{Y}_j, \boldsymbol{X}_j)$ are identically distributed when $j \in \mathcal{N}$. The third equality holds because $t^{\text{minus}}(\boldsymbol{Y}_j, \boldsymbol{X}_j) = -C_j$; if $-C_j \in \mathcal{A}$, then $S_j = -1$.*

*Because $\mathbb{P}\left(S_j = 1, |C_j| \in \mathcal{A}\right) + \mathbb{P}\left(S_j = -1, |C_j| \in \mathcal{A}\right) = \mathbb{P}\left(|C_j| \in \mathcal{A}\right)$, it follows that*

$$\mathbb{P}\left(S_j = 1, |C_j| \in \mathcal{A}\right) = \frac{1}{2}\mathbb{P}\left(|C_j| \in \mathcal{A}\right) = \mathbb{P}(S_j = 1)\mathbb{P}\left(|C_j| \in \mathcal{A}\right),$$

*where the last equality holds because $\mathbb{P}(S_j = 1) = 1/2$ by property (b).*

*Hence, $S_j$ and $|C_j|$ are independent $\forall j \in \mathcal{N}$.*

**Proof 2 (Proof of Theorem 1)** *Define a random subset of $\mathcal{N}$ as $\mathcal{M} := \mathcal{N} \backslash \{j \in \mathcal{N} : C_j = 0\} = \{j \in \mathcal{N} : S_j \neq 0\}$.*

*First note that by Lemma 1(b), $\mathbb{P}(S_j = -1) = \mathbb{P}(C_j < 0) = 1/2$ for all $j \in \mathcal{M} \subset \mathcal{N}$. Assume without loss of generality that $\mathcal{M} = \{1, \ldots, d'\}$. We order $\{|C_j| : j \in \mathcal{M}\}$, from the largest to the smallest, denoted by $|C_{(1)}| \geq |C_{(2)}| \geq \cdots \geq |C_{(d')}|$. Let $J = \sum_{j \in \mathcal{N}} \mathbb{1}\left(|C_j| \geq T^{\text{BC}}\right)$, the number of uninteresting features whose contrast scores have absolute values no less than $T^{\text{BC}}$. When $J > 0$, $|C_{(1)}| \geq \cdots \geq |C_{(J)}| \geq T^{\text{BC}}$. Define $Z_k = \mathbb{1}\left(C_{(k)} < 0\right)$, $k = 1, \ldots, d'$. Then for each order $k$, the following holds*

$$C_{(k)} \geq T^{\text{BC}} \iff \left|C_{(k)}\right| \geq T^{\text{BC}} \text{ and } C_{(k)} > 0 \iff k \leq J \text{ and } Z_k = 0;$$
$$C_{(k)} \leq -T^{\text{BC}} \iff \left|C_{(k)}\right| \geq T^{\text{BC}} \text{ and } C_{(k)} < 0 \iff k \leq J \text{ and } Z_k = 1.$$

*Then*

$$\begin{aligned}
\frac{\text{card}(\{j \in \mathcal{M} : C_j \geq T^{\text{BC}}\})}{\text{card}(\{j \in \mathcal{M} : C_j \leq -T^{\text{BC}}\}) + 1} &= \frac{\sum_{k=1}^{d'} \mathbb{1}\left(C_{(k)} \geq T^{\text{BC}}\right)}{1 + \sum_{k=1}^{d'} \mathbb{1}\left(C_{(k)} \leq -T^{\text{BC}}\right)} \\
&= \frac{\sum_{k=1}^{J} \mathbb{1}\left(C_{(k)} \geq T^{\text{BC}}\right)}{1 + \sum_{k=1}^{J} \mathbb{1}\left(C_{(k)} \leq -T^{\text{BC}}\right)} \\
&= \frac{(1 - Z_1) + \cdots + (1 - Z_J)}{1 + Z_1 + \cdots + Z_J} \\
&= \frac{1 + J}{1 + Z_1 + \cdots + Z_J} - 1.
\end{aligned}$$

*Because $\{S_j\}_{j \in \mathcal{N}}$ is independent of $\mathcal{C}$ (Lemma 1(c)), Lemma 1(a)-(b) still holds after $C_1, \ldots, C_{d'}$ are* *reordered as $C_{(1)}, \ldots, C_{(d')}$. Thus $Z_1, \ldots, Z_{d'}$ are i.i.d. from $\mathrm{Bernoulli}(1/2)$. To summarize, it holds that*

$$\{Z_j\}_{j\in\mathcal{M}} \,\Big|\, \mathcal{M} \overset{\text{i.i.d.}}{\sim} \mathrm{Bernoulli}(1/2)\,.$$

*Then by applying Lemma 2 and making $\rho = 0.5$, we have:*

$$\mathbb{E}\left[\frac{\mathrm{card}(\{j\in\mathcal{M}: C_j \geq T^{\mathrm{BC}}\})}{\mathrm{card}(\{j\in\mathcal{M}: C_j \leq -T^{\mathrm{BC}}\})+1} \,\middle|\, \mathcal{M}\right] \leq 1 \tag{S22}$$

*Then*

$$\begin{aligned}
\mathrm{FDR} &= \mathbb{E}\left[\frac{\mathrm{card}(\{j\in\mathcal{N}: C_j \geq T^{\mathrm{BC}}\})}{\mathrm{card}(\{j: C_j \geq T^{\mathrm{BC}}\})\vee 1}\right] \\
&= \mathbb{E}\left[\frac{\mathrm{card}(\{j\in\mathcal{N}: C_j \geq T^{\mathrm{BC}}\})}{\mathrm{card}(\{j\in\mathcal{N}: C_j \leq -T^{\mathrm{BC}}\})+1}\cdot\frac{\mathrm{card}(\{j\in\mathcal{N}: C_j \leq -T^{\mathrm{BC}}\})+1}{\mathrm{card}(\{j: C_j \geq T^{\mathrm{BC}}\})\vee 1}\right] \\
&\leq \mathbb{E}\left[\frac{\mathrm{card}(\{j\in\mathcal{N}: C_j \geq T^{\mathrm{BC}}\})}{\mathrm{card}(\{j\in\mathcal{N}: C_j \leq -T^{\mathrm{BC}}\})+1}\cdot\frac{\mathrm{card}(\{j: C_j \leq -T^{\mathrm{BC}}\})+1}{\mathrm{card}(\{j: C_j \geq T^{\mathrm{BC}}\})\vee 1}\right] \\
&\leq q\cdot\mathbb{E}\left[\frac{\mathrm{card}(\{j\in\mathcal{N}: C_j \geq T^{\mathrm{BC}}\})}{\mathrm{card}(\{j\in\mathcal{N}: C_j \leq -T^{\mathrm{BC}}\})+1}\right] \\
&\leq q\cdot\mathbb{E}\left[\mathbb{E}\left[\frac{\mathrm{card}(\{j\in\mathcal{M}: C_j \geq T^{\mathrm{BC}}\})}{\mathrm{card}(\{j\in\mathcal{M}: C_j \leq -T^{\mathrm{BC}}\})+1}\,\middle|\,\mathcal{M}\right]\right] \\
&\leq q,
\end{aligned}$$

*where $\mathcal{M}$ is random subset of $\mathcal{N}$ such that for each $j \in \mathcal{M}$, $|C_j| > 0$. The last inequality follows from (S22).*

#### S8.2 Proof of Theorem 2

We then prove Theorem 2, which relies on Lemmas 2 and 3. Here we introduce the proof of Lemma 3 and defer the proof of Lemma 2 to Section S8.3.

**Proof 3 (Proof of Lemma 3)** *With input data satisfying (S6) and (S7), $C_j$ constructed from (S19) or (S20), property (a) can be derived directly.*

*To show property (b), note that for each uninteresting feature $j \in \mathcal{N}$, $\boldsymbol{X}_j$ and $\boldsymbol{Y}_j$ are from the same distribution; thus $\{T_j^{\sigma_\ell}\}_{\ell=0}^h$ are identically distributed. Define an event $\mathcal{E}_j := \left\{\sum_{\ell=0}^h \mathbb{1}(T_j^{\sigma_\ell} = T_j^{(0)}) = 1\right\}$, which indicates that $T_j^{(0)}$, the maximizer of $\{T_j^{\sigma_\ell}\}_{\ell=0}^h$, is unique. Then conditional on $\mathcal{E}_j$, the maximizer is equally likely to be any of $\{0,\ldots,h\}$, and it follows that $\mathbb{P}(S_j = 1 \mid \mathcal{E}_j) = \mathbb{P}(T_j^{\sigma_0} = T_j^{(0)} \mid \mathcal{E}_j) = 1/(h+1)$. Conditioning on that $\mathcal{E}_j$ does not happen, $\mathbb{P}(S_j = 1 \mid \mathcal{E}_j^c) = 0$. Thus $\mathbb{P}(S_j = 1) = \mathbb{P}(S_j = 1 \mid \mathcal{E}_j)\mathbb{P}(\mathcal{E}_j) + \mathbb{P}(S_j = 1 \mid \mathcal{E}_j^c)\mathbb{P}(\mathcal{E}_j^c) \leq 1/(h+1)$.*

*The proof of property (c) is similar to the Proof of Lemma 1(c). It suffices to show that for any $j \in \mathcal{N}$ with $C_j \neq 0$ (that is, $\mathcal{E}_j$ occurs), $S_j$ is independent of $|C_j|$. As $\boldsymbol{X}_j$ and $\boldsymbol{Y}_j$ are from the same distribution, $\{T_j^{\sigma_\ell}\}_{\ell=0}^h$ are identically distributed. Hence for any measurable set $\mathcal{A} \subset [0,+\infty)$,*

$$\begin{aligned}
\mathbb{P}\left(S_j = 1, |C_j| \in \mathcal{A} \mid \mathcal{E}_j\right) &= \mathbb{P}\left(T_j^{\sigma_0} = T_j^{(0)}, |C_j| \in \mathcal{A} \,\middle|\, \mathcal{E}_j\right) \\
&= \frac{1}{h}\mathbb{P}\left(T_j^{\sigma_0} \neq T_j^{(0)}, |C_j| \in \mathcal{A} \,\middle|\, \mathcal{E}_j\right) \\
&= \frac{1}{h}\mathbb{P}\left(S_j = -1, |C_j| \in \mathcal{A} \mid \mathcal{E}_j\right).
\end{aligned}$$

*The first equality holds because $T_j^{\sigma_0} = T_j^{(0)}$ when $S_j = 1$. The second equality holds because $\{T_j^{\sigma_\ell}\}_{\ell=0}^h$ are identically distributed when $j \in \mathcal{N}$. The third equality holds because $T_j^{\sigma_0} \neq T_j^{(0)}$ when $S_j = -1$.*

*Because $\mathbb{P}(S_j = 1, |C_j| \in \mathcal{A} \mid \mathcal{E}_j) + \mathbb{P}(S_j = -1, |C_j| \in \mathcal{A} \mid \mathcal{E}_j) = \mathbb{P}(|C_j| \in \mathcal{A} \mid \mathcal{E}_j)$, it follows that*

$$\mathbb{P}(S_j = 1, |C_j| \in \mathcal{A} \mid \mathcal{E}_j) = \frac{1}{h+1}\mathbb{P}(|C_j| \in \mathcal{A} \mid \mathcal{E}_j) = \mathbb{P}(S_j = 1 \mid \mathcal{E}_j)\mathbb{P}(|C_j| \in \mathcal{A} \mid \mathcal{E}_j) ,$$

*where the last equality holds because $\mathbb{P}(S_j = 1 \mid \mathcal{E}_j) = 1/(h+1)$.*

*Hence, $S_j$ and $|C_j|$ are independent $\forall j \in \mathcal{N}$ with $C_j \neq 0$.*

**Proof 4 (Proof of Theorem 2)** *Define a random subset of $\mathcal{N}$ as $\mathcal{M} := \mathcal{N} \backslash \{j \in \mathcal{N} : C_j = 0\} = \{j \in \mathcal{N} : S_j \neq 0\}$. Assume without loss of generality that $\mathcal{M} = \{1, \ldots, d'\}$. We order $\{|C_j| : j \in \mathcal{M}\}$, from the largest to the smallest, denoted by $|C_{(1)}| \geq |C_{(2)}| \geq \cdots \geq |C_{(d')}|$. Let $J = \sum_{j \in \mathcal{N}} \mathbb{1}\left(|C_j| \geq T^{\mathrm{GZ}}\right)$, the number of uninteresting features whose contrast scores have absolute values no less than $T^{\mathrm{GZ}}$. When $J > 0$, $|C_{(1)}| \geq \cdots \geq |C_{(J)}| \geq T^{\mathrm{GZ}}$. Define $Z_k = \mathbb{1}\left(C_{(k)} < 0\right)$, $k = 1, \ldots, d'$. Then for each order $k$, the following holds:*

$$C_{(k)} \geq T^{\mathrm{GZ}} \iff \left|C_{(k)}\right| \geq T^{\mathrm{GZ}} \text{ and } C_{(k)} > 0 \iff k \leq J \text{ and } Z_k = 0\,;$$
$$C_{(k)} \leq -T^{\mathrm{GZ}} \iff \left|C_{(k)}\right| \geq T^{\mathrm{GZ}} \text{ and } C_{(k)} < 0 \iff k \leq J \text{ and } Z_k = 1\,.$$

*Then it follows that*

$$\begin{aligned}
\frac{\mathrm{card}(\{j \in \mathcal{M} : C_j \geq T^{\mathrm{GZ}}\})}{\mathrm{card}(\{j \in \mathcal{M} : C_j \leq -T^{\mathrm{GZ}}\}) + 1} &= \frac{\sum_{k=1}^{d'} \mathbb{1}\left(C_{(k)} \geq T^{\mathrm{GZ}}\right)}{1 + \sum_{k=1}^{d'} \mathbb{1}\left(C_{(k)} \leq -T^{\mathrm{GZ}}\right)} \\
&= \frac{\sum_{k=1}^{J} \mathbb{1}\left(C_{(k)} \geq T^{\mathrm{GZ}}\right)}{1 + \sum_{k=1}^{J} \mathbb{1}\left(C_{(k)} \leq -T^{\mathrm{GZ}}\right)} \\
&= \frac{(1 - Z_1) + \cdots + (1 - Z_J)}{1 + Z_1 + \cdots + Z_J} \\
&= \frac{1 + J}{1 + Z_1 + \cdots + Z_J} - 1\,.
\end{aligned}$$

*Because $\{S_j\}_{j \in \mathcal{N}}$ is independent of $\mathcal{C}$ (Lemma 1(c)), Lemma 1(a)-(b) still holds after $C_1, \ldots, C_{d'}$ are reordered as $C_{(1)}, \ldots, C_{(d')}$. Thus $Z_1, \ldots, Z_{d'}$ are i.i.d. from $\mathrm{Bernoulli}(\rho_k)$. To summarize, it holds that*

$$\{Z_j\}_{j \in \mathcal{M}} \,\Big|\, \mathcal{M} \overset{\text{i.i.d.}}{\sim} \mathrm{Bernoulli}(\rho_k)\,.$$

*Then by applying Lemma 2 and making $\rho = h/(h+1)$, we have:*

$$\mathbb{E}\left[\frac{\mathrm{card}(\{j \in \mathcal{M} : C_j \geq T^{\mathrm{GZ}}\})}{\mathrm{card}(\{j \in \mathcal{M} : C_j \leq -T^{\mathrm{GZ}}\}) + 1}\right] \leq 1/h\,. \tag{S23}$$

*Then*

$$
\begin{aligned}
\mathrm{FDR} &= \mathbb{E}\left[\frac{\mathrm{card}(\{j\in\mathcal{N}: C_j \geq T^{\mathrm{GZ}}\})}{\mathrm{card}(\{j: C_j \geq T^{\mathrm{GZ}}\})\vee 1}\right] \\
&= \mathbb{E}\left[\frac{\mathrm{card}(\{j\in\mathcal{N}: C_j \geq T^{\mathrm{GZ}}\})}{\mathrm{card}(\{j\in\mathcal{N}: C_j \leq -T^{\mathrm{GZ}}\})+1}\cdot\frac{\mathrm{card}(\{j\in\mathcal{N}: C_j \leq -T^{\mathrm{GZ}}\})+1}{\mathrm{card}(\{j: C_j \geq T^{\mathrm{GZ}}\})\vee 1}\right] \\
&\leq h\cdot\mathbb{E}\left[\frac{\mathrm{card}(\{j\in\mathcal{N}: C_j \geq T^{\mathrm{GZ}}\})}{\mathrm{card}(\{j\in\mathcal{N}: C_j \leq -T^{\mathrm{GZ}}\})+1}\cdot\frac{\frac{1}{h}\mathrm{card}(\{j: C_j \leq -T^{\mathrm{GZ}}\})+\frac{1}{h}}{\mathrm{card}(\{j: C_j \geq T^{\mathrm{GZ}}\})\vee 1}\right] \\
&\leq hq\cdot\mathbb{E}\left[\frac{\mathrm{card}(\{j\in\mathcal{N}: C_j \geq T^{\mathrm{GZ}}\})}{\mathrm{card}(\{j\in\mathcal{N}: C_j \leq -T^{\mathrm{GZ}}\})+1}\right] \\
&\leq hq\cdot\mathbb{E}\left[\mathbb{E}\left[\frac{\mathrm{card}(\{j\in\mathcal{M}: C_j \geq T^{\mathrm{GZ}}\})}{\mathrm{card}(\{j\in\mathcal{M}: C_j \leq -T^{\mathrm{GZ}}\})+1}\;\middle|\;\mathcal{M}\right]\right] \\
&\leq q\,,
\end{aligned}
$$

*where the second inequality follows from the definition of* $T_{\mathrm{GZ}}$ (S21) *and the last inequality follows from* (S23).

#### S8.3 Proof of Lemma 2

Finally, we derive Lemma 2 by following the same proof same as in [8], which relies on Lemma 4 and Corollary 1.

**Lemma 4** *Suppose that* $Z_1,\ldots,Z_d \overset{i.i.d.}{\sim} \mathrm{Bernoulli}(\rho)$*. Let* $J$ *be a stopping time in reverse time with respect to the filtration* $\{\mathcal{F}_j\}$*, where* $\mathcal{F}_j = \sigma\left(\{(Z_1+\cdots+Z_j),\, Z_{j+1},\,\cdots,\, Z_d\}\right)$ *with* $\sigma(\cdot)$ *denoting a* $\sigma$*-algebra, and the variables* $Z_1,\ldots,Z_j$ *are exchangeable with respect to* $\{\mathcal{F}_j\}$*. Then*

$$
\mathbb{E}\left[\frac{1+J}{1+Z_1+\cdots+Z_J}\right] \leq \rho^{-1}.
$$

**Proof 5 (Proof of Lemma 4)** *Define*

$$
Y_j = Z_1+\cdots+Z_j \in \mathcal{F}_j
$$

*and define the process*

$$
M_j = \frac{1+j}{1+Z_1+\cdots+Z_j} = \frac{1+j}{1+Y_j} \in \mathcal{F}_j.
$$

*In [3], it is shown that* $\mathbb{E}[M_d] \leq \rho^{-1}$*. Therefore, by the optional stopping time theorem it suffices to show that* $\{M_j\}$ *is a supermartingale with respect to* $\{\mathcal{F}_j\}$*. As* $\{Z_1,\ldots,Z_{j+1}\}$ *are exchangeable with respect to* $\mathcal{F}_{j+1}$*, we have*

$$
\mathbb{P}\left(Z_{j+1}=1 \mid \mathcal{F}_{j+1}\right) = \frac{Y_j+1}{1+j}.
$$

*Therefore, if $Y_{j+1} > 0$,*

$$
\begin{aligned}
\mathbb{E}\left[M_j \mid \mathcal{F}_{j+1}\right] &= \frac{1+j}{1+Y_{j+1}} \cdot \mathbb{P}\left(Z_{j+1}=0 \mid \mathcal{F}_{j+1}\right) + \frac{1+j}{1+Y_{j+1}-1} \cdot \mathbb{P}\left(Z_{j+1}=1 \mid \mathcal{F}_{j+1}\right) \\
&= \frac{1+j}{1+Y_{j+1}} \cdot \frac{1+j-Y_{j+1}}{1+j} + \frac{1+j}{1+Y_{j+1}-1} \cdot \frac{Y_{j+1}}{1+j} \\
&= \frac{1+j-Y_{j+1}}{1+Y_{j+1}} + 1 \\
&= \frac{1+(j+1)}{1+Y_{j+1}} \\
&= M_{j+1}.
\end{aligned}
$$

*If instead $Y_{j+1} = 0$, then trivially $Y_j = 0$, and $M_j = 1 + j < 2 + j = M_{j+1}$. This proves that $\{M_j\}$ is a supermartingale with respect to $\{\mathcal{F}_j\}$ as desired.*

**Corollary 1** *Suppose that $\mathcal{A} \subseteq \{1, \ldots, d\}$ is fixed, while $Z_1, \ldots, Z_d \overset{i.i.d.}{\sim} \text{Bernoulli}(\rho)$. Let $J$ be a stopping time in reverse time with respect to the filtration $\{\mathcal{F}_j\}$, where $\mathcal{F}_j = \sigma\left(\{\sum_{k \leq j, k \in \mathcal{A}} Z_k\} \cup \{Z_k : j < k < d, k \in \mathcal{A}\}\right)$ with $\sigma(\cdot)$ denoting a $\sigma$-algebra, and the variables $\{Z_k : k \leq j, k \in \mathcal{A}\}$ are exchangeable with respect to $\mathcal{F}_j$. Then*

$$
\mathbb{E}\left[\frac{1 + \text{card}\left(\{k : k \leq J, k \in \mathcal{A}\}\right)}{1 + \sum_{k \leq J, k \in \mathcal{A}} Z_k}\right] \leq \rho^{-1}.
$$

**Proof 6 (Proof of Corollary 1)** *Let $\mathcal{A} = \{j_1, \ldots, j_m\}$ where $1 \leq j_1 < \cdots < j_m \leq d$. Then by considering the i.i.d. sequence*

$$
Z_{j_1}, \ldots, Z_{j_m}
$$

*in place of $Z_1, \ldots, Z_d$, we see that this result is equivalent to Lemma 4.*

**Proof 7 (Proof of Lemma 2)** *[From [3]] We may assume $\rho < 1$ to avoid the trivial case. We first introduce a different definition for $\{Z_j\}_{j=1}^d$ by defining a random set $\mathcal{A} \subseteq \{1, \ldots, d\}$ where for each $j$, independently,*

$$
\mathbb{P}\left(j \in \mathcal{A}\right) = \frac{1-\rho_j}{1-\rho}.
$$

*We then define random variables $Q_1, \ldots, Q_d \overset{i.i.d.}{\sim} \text{Bernoulli}(\rho)$, which are generated independently of the random set $\mathcal{A}$. Finally, we define*

$$
Z_j = Q_j \cdot \mathbb{1}\left(j \in \mathcal{A}\right) + \mathbb{1}\left(j \notin \mathcal{A}\right). \tag{S24}
$$

*Then $\{Z_j\}_{k=1}^d$ are mutually independent and $\mathbb{P}\left(Z_j = 1\right) = 1 - \mathbb{P}(j \in \mathcal{A}) \cdot \mathbb{P}(Q_j = 0) = \rho_j$, that is, $Z_j \sim \text{Bernoulli}(\rho_j)$. This new definition of $\{Z_j\}_{j=1}^d$ meet all the conditions required by Lemma 2, so that we can apply this new definition in the following proof.*

*As $Z_j = Q_j \cdot \mathbb{1}\left(j \in \mathcal{A}\right) + \mathbb{1}\left(j \notin \mathcal{A}\right)$ for all $j$, we have*

$$
\frac{1+J}{1+Z_1+\cdots+Z_J} = \frac{1 + \text{card}\left(\{j \leq J : j \in \mathcal{A}\}\right) + \text{card}\left(\{j \leq J : j \notin \mathcal{A}\}\right)}{1 + \sum_{j \leq J, j \in \mathcal{A}} Q_j + \text{card}\left(\{j \leq J : j \notin \mathcal{A}\}\right)} \leq \frac{1 + \text{card}\left(\{j \leq J : j \in \mathcal{A}\}\right)}{1 + \sum_{j \leq J, j \in \mathcal{A}} Q_j}, \tag{S25}
$$

*where the last step uses the identify $\frac{a+c}{b+c} \leq \frac{a}{b}$ whenever $0 < b \leq a$ and $c \geq 0$. Therefore, it will be sufficient to prove that*

$$
\mathbb{E}\left[\frac{1 + \text{card}\left(\{j \leq J : j \in \mathcal{A}\}\right)}{1 + \sum_{j \leq J, j \in \mathcal{A}} Q_j} \,\middle|\, \mathcal{A}\right] \leq \rho^{-1}, \tag{S26}
$$

*To prove (S26), first let $\widetilde{Q}_j = Q_j \cdot \mathbb{1}(j \in \mathcal{A})$, and define a filtration $\{\mathcal{F}'_j\}$ where $\mathcal{F}'_j$ is the $\sigma$-algebra generated as*

$$\mathcal{F}'_j = \sigma\left(\left\{\widetilde{Q}_1 + \cdots + \widetilde{Q}_j, \widetilde{Q}_{j+1}, \cdots, \widetilde{Q}_d, \mathcal{A}\right\}\right).$$

*Next for any $j$, by (S24) we see that*

$$Z_1 + \cdots + Z_j, Z_{j+1}, \ldots, Z_d \in \mathcal{F}'_j \Rightarrow \mathcal{F}_j \subseteq \mathcal{F}'_j,$$

*so $J$ is a stopping time (in reverse time) with respect to $\mathcal{F}'_j$. Finally, since the $Q_j$'s are independent of $\mathcal{A}$, (S26) follows from Corollary 1 after conditioning on $\mathcal{A}$.*

### S9 Supplementary figures

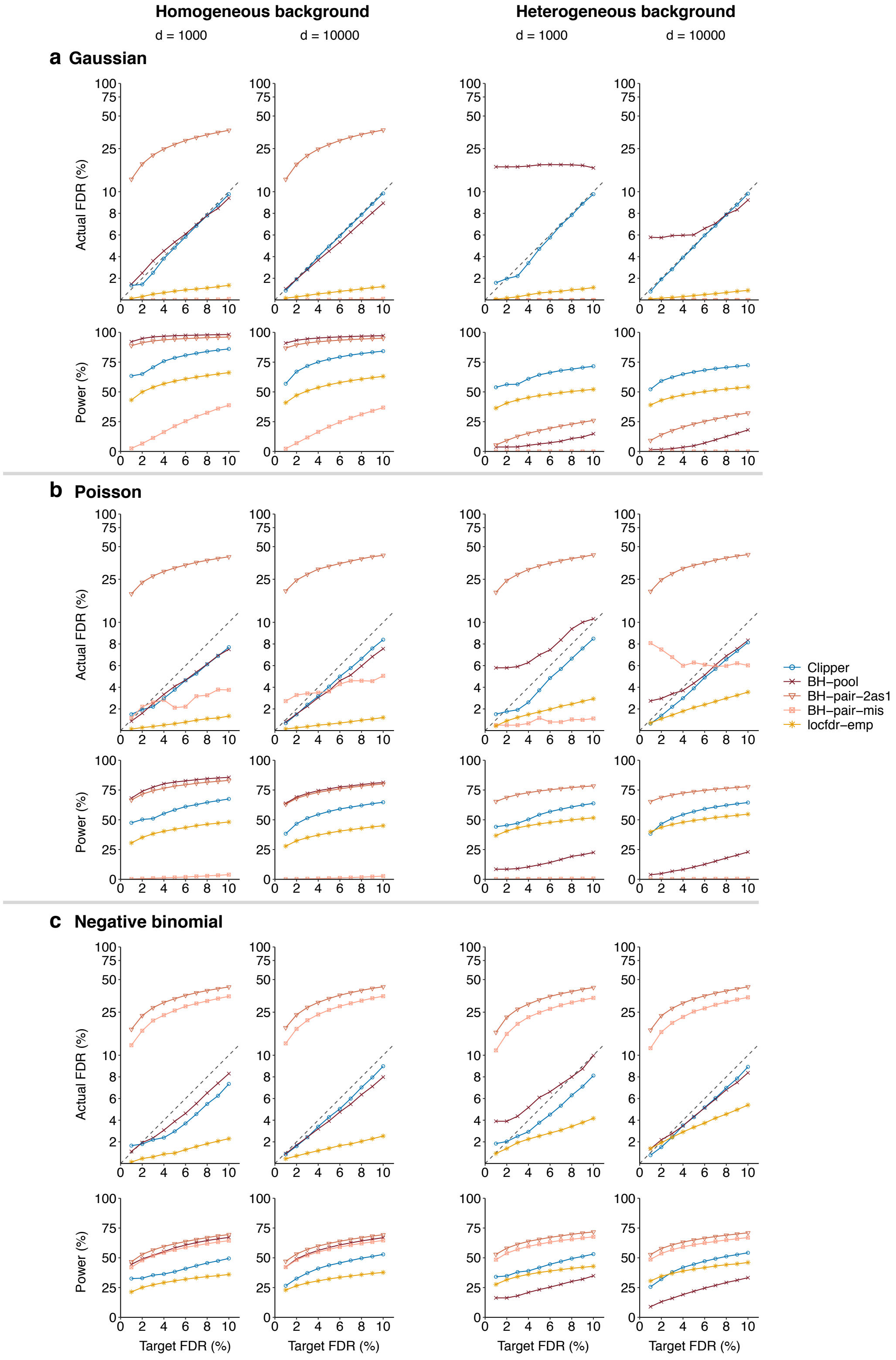


**Figure S1:** In the 1vs1 enrichment analysis, comparison of Clipper and four other generic FDR control methods (BH-pool, BH-pair-2as1, BH-pair-mis, and locfdr-emp) in terms of their FDR control and power. At target FDR thresholds $q \in \{1\%, 2\%, \cdots, 10\%\}$, each method's actual FDRs and power are evaluated on $200$ simulated datasets with $d = 1000$ or $10{,}000$ features generated from (**a**) the Gaussian distribution, (**b**) the Poisson distribution, or (**c**) the negative binomial distribution under homogeneous (two left columns) and heterogeneous (two right columns) background scenarios. Among the methods that control the FDR, Clipper is the second most powerful for homogeneous background and the most powerful for heterogeneous background.

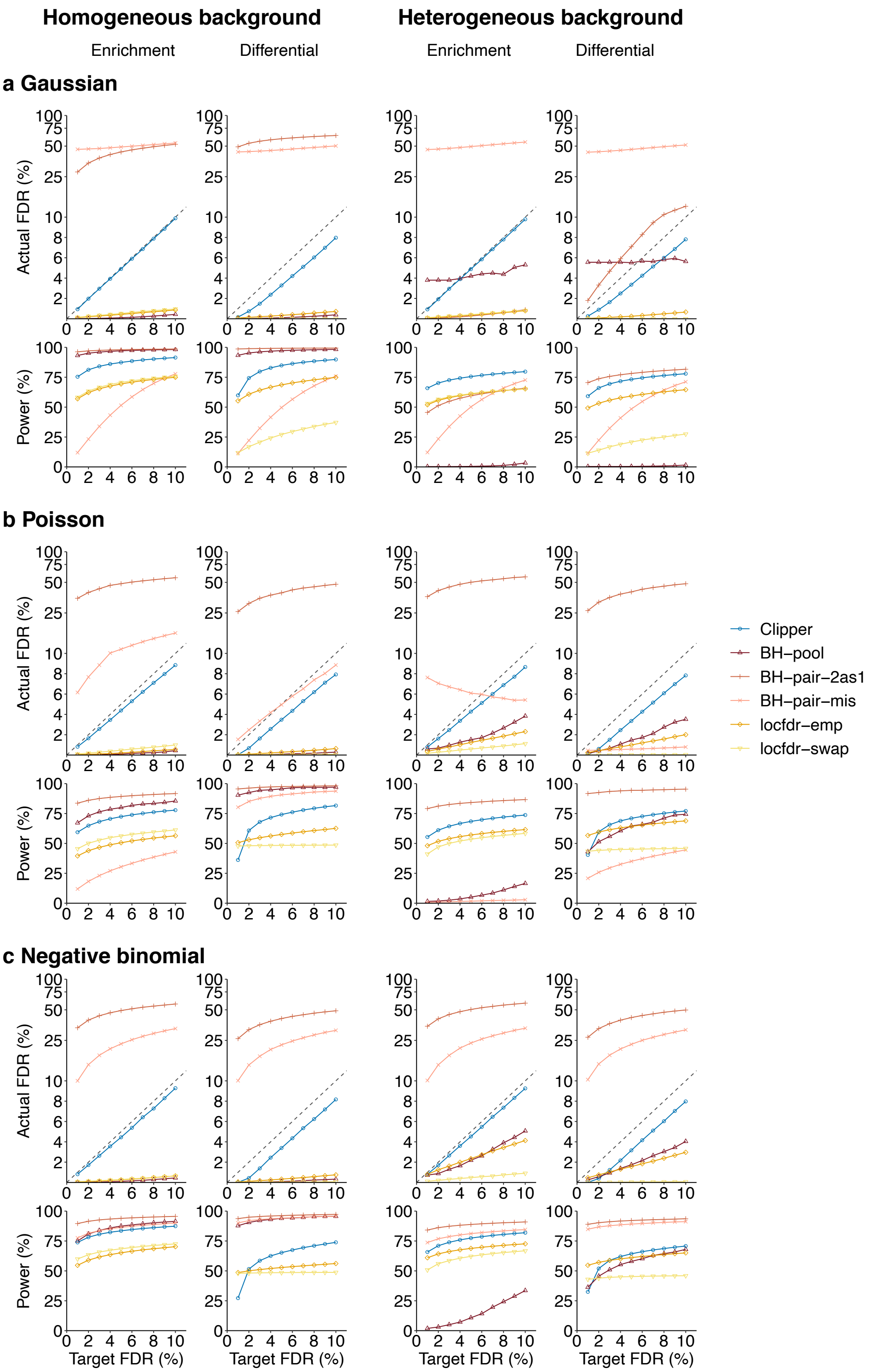


**Figure S2:** In the 2vs1 enrichment analysis (columns 1 and 3) and differential analysis (columns 2 and 4), comparison of Clipper and five other generic FDR control methods (BH-pooled, BH-pair-2as1, BH-pair-mis, locfdr-emp, and locfdr-swap) in terms of their FDR control and power. At target FDR thresholds $q \in \{1\%, 2\%, \cdots, 10\%\}$, each method's actual FDRs and power are evaluated on $200$ simulated datasets with $d = 10{,}000$ features generated from (**a**) the Gaussian distribution, (**b**) the Poisson distribution, or (**c**) the negative binomial distribution under homogeneous(two left columns) and heterogeneous (two right columns) background scenarios. Among the methods that control the FDR, Clipper is the second most powerful for homogeneous background and the most powerful for heterogeneous background (except for differential analysis with $q \leq 2\%$).

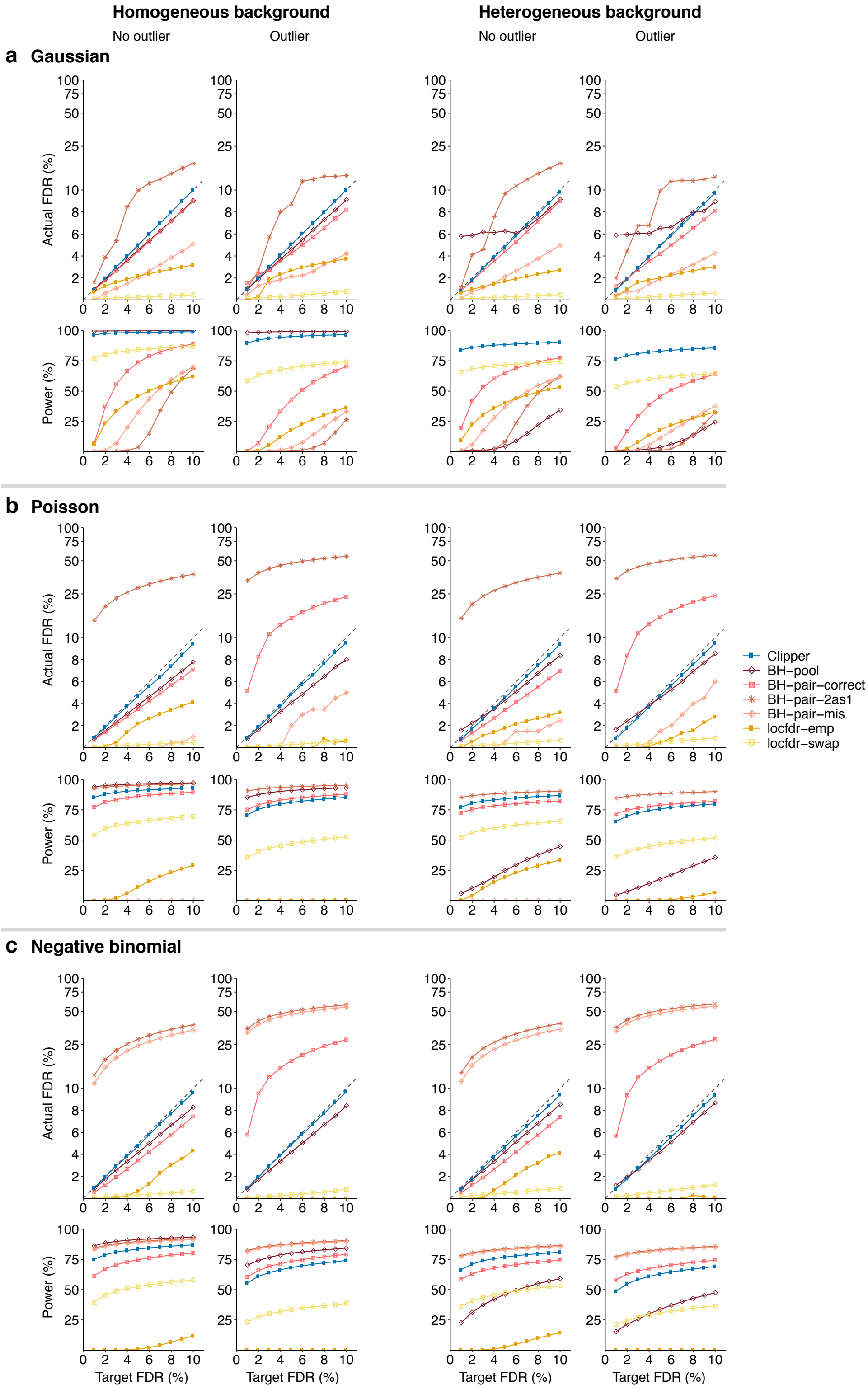


**Figure S3:** In the 3vs3 enrichment analysis without (columns 1 and 3) or with outliers (columns 2 and 4), comparison of Clipper and six other generic FDR control methods (BH-pooled, BH-pair-correct, BH-pair-2as1, BH-pair-mis, locfdr-emp, and locfdr-swap)in terms of their FDR control and power in 3vs3 enrichment analysis with possible outliers. At target FDR thresholds $q \in \{1\%, 2\%, \cdots, 10\%\}$, each method's actual FDRs and power are evaluated on $200$ simulated datasets with $d = 10{,}000$ features generated from (**a**) the Gaussian distribution, (**b**) the Poisson distribution, or (**c**) the negative binomial distribution under homogeneous (two left columns) and heterogeneous (two right columns) background scenarios. $10\%$ of the features are interesting features. Among the methods that control the FDR, Clipper is the second most powerful for homogeneous background and the most powerful for heterogeneous background.

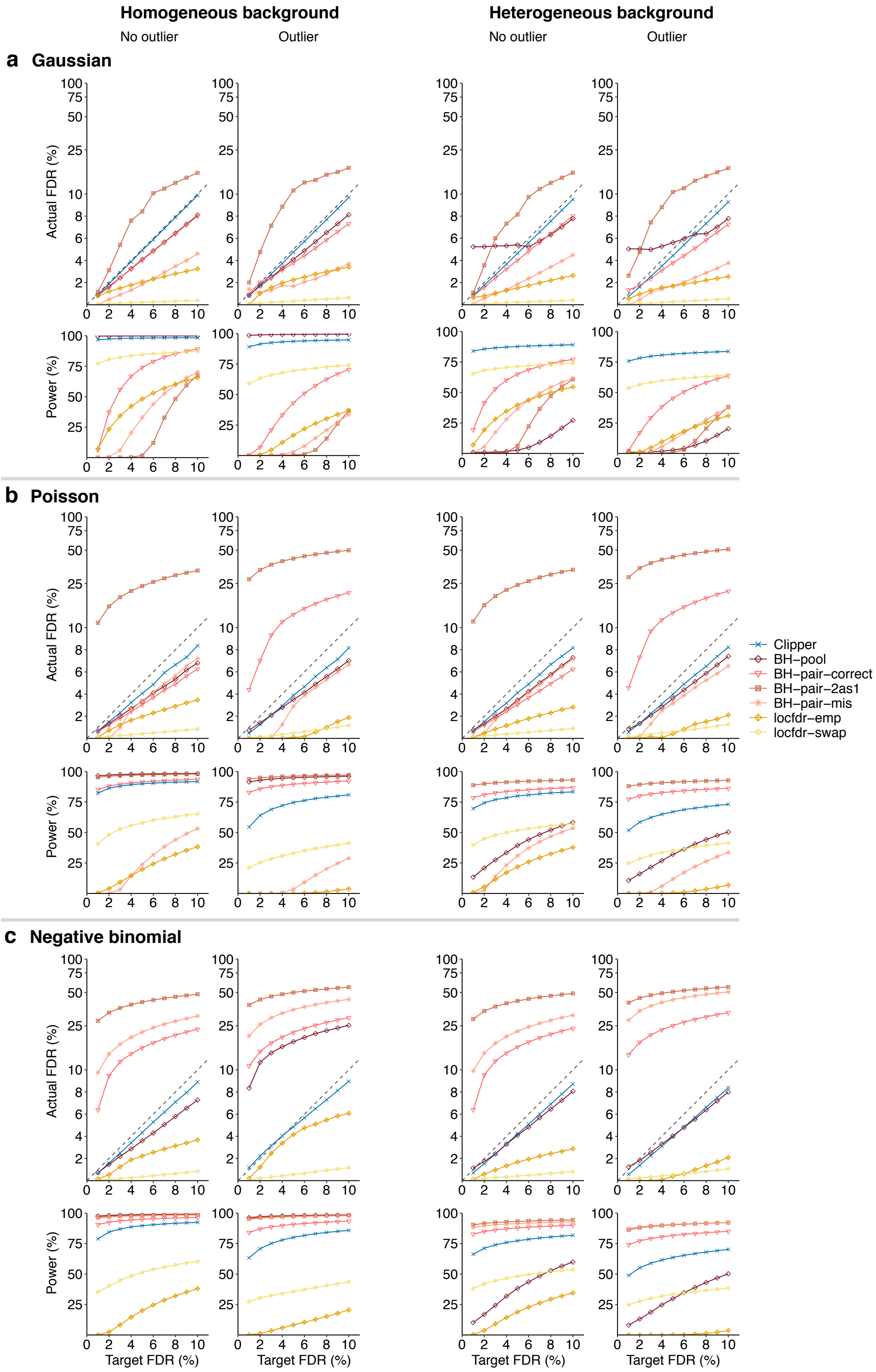


**Figure S4:** In the 3vs3 differential analysis without (columns 1 and 3) or with outliers (columns 2 and 4), comparison of Clipper and six other generic FDR control methods (BH-pooled, BH-pair-correct, BH-pair-2as1, BH-pair-mis, locfdr-emp, and locfdr-swap) in terms of their FDR control and power. At target FDR thresholds $q \in \{1\%, 2\%, \cdots, 10\%\}$, each method's actual FDRs and power are evaluated on $200$ simulated datasets with $d = 10{,}000$ features generated from (**a**) the Gaussian distribution, (**b**) the Poisson distribution, or (**c**) the negative binomial distribution under homogeneous (two left columns) and heterogeneous (two right columns) background scenarios. Among the methods that control the FDR, Clipper is the second most powerful for homogeneous background and the most powerful for heterogeneous background (except for Poisson distribution where Clipper is second to BH-pair-correct, an idealistic method).

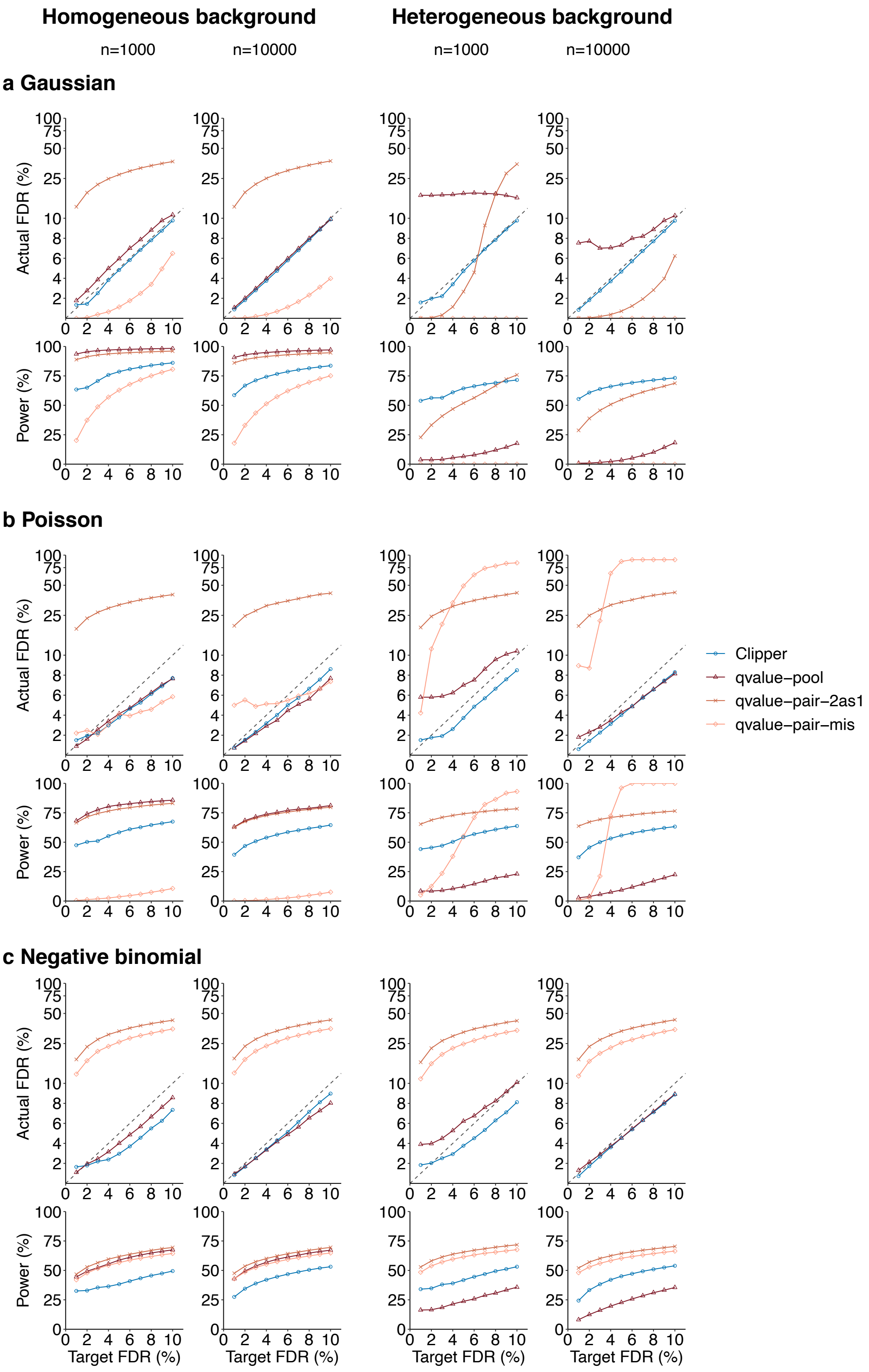


**Figure S5:** In the 1vs1 enrichment analysis, comparison of Clipper and three other generic FDR control methods using Storey's q-value (qvalue-pool, qvalie-pair-2as1, and qvalue-pair-mis) in terms of their FDR control and power. At target FDR thresholds $q \in \{1\%, 2\%, \cdots, 10\%\}$, each method's actual FDRs and power are evaluated on $200$ simulated datasets with $d = 1000$ or $10{,}000$ features generated from (**a**) the Gaussian distribution, (**b**) the Poisson distribution, or (**c**) the negative binomial distribution under homogeneous (two left columns) and heterogeneous (two right columns) background scenarios. Among the methods that control the FDR, Clipper is the second most powerful for homogeneous background and the most powerful for heterogeneous background.

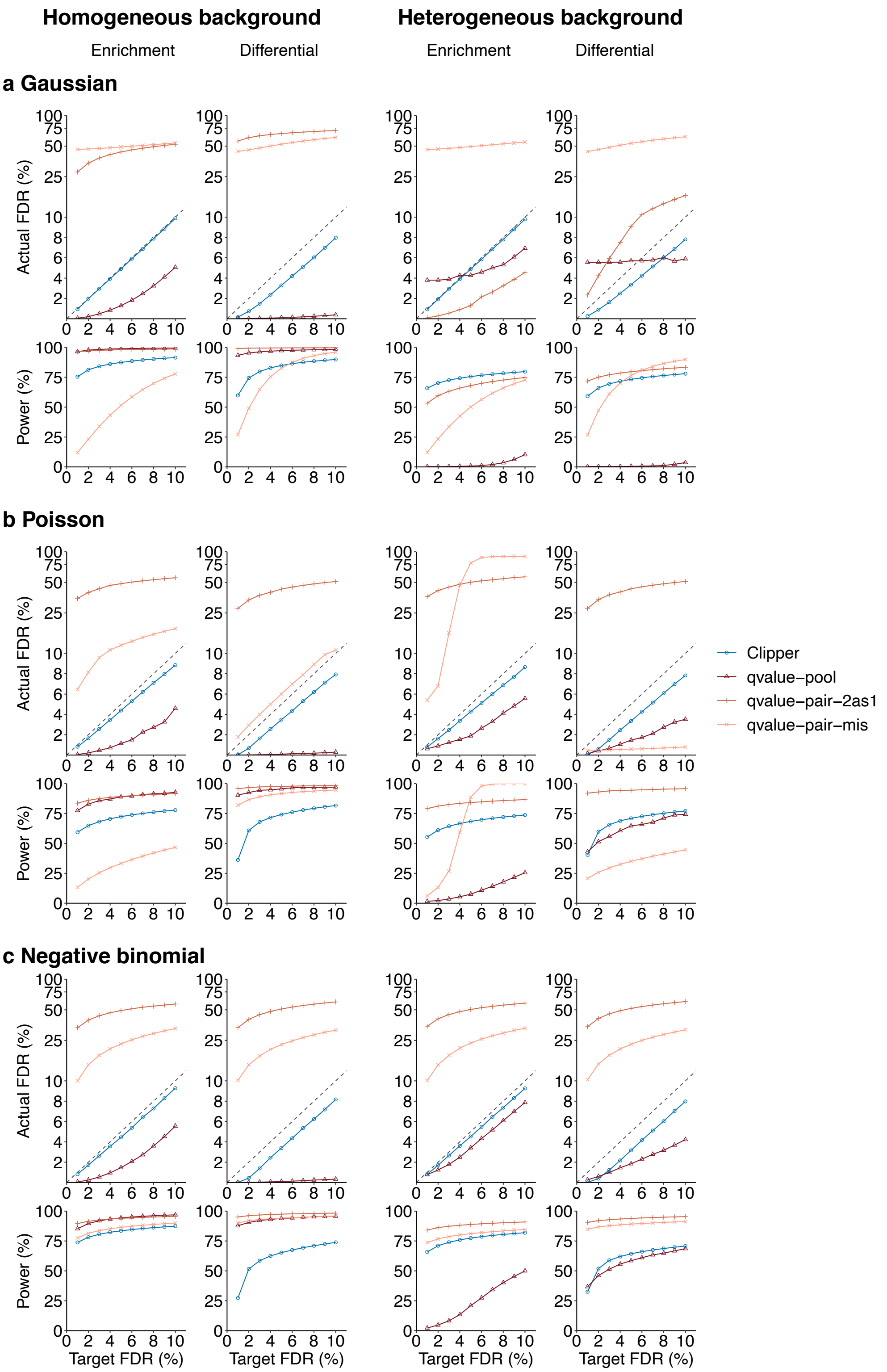


**Figure S6:** In the 2vs1 enrichment analysis (columns 1 and 3) and differential analysis (columns 2 and 4), comparison of Clipper and three other generic FDR control methods using Storey's q-value (qvalue-pool, qvalie-pair-2as1, and qvalue-pair-mis) in terms of their FDR control and power. At target FDR thresholds $q \in \{1\%, 2\%, \cdots, 10\%\}$, each method's actual FDRs and power are evaluated on $200$ simulated datasets with $d = 10{,}000$ features generated from (**a**) the Gaussian distribution, (**b**) the Poisson distribution, or (**c**) the negative binomial distribution under homogeneous (two left columns) and heterogeneous (two right columns) background scenarios. Among the methods that control the FDR, Clipper is the second most powerful for homogeneous background and the most powerful for heterogeneous background.

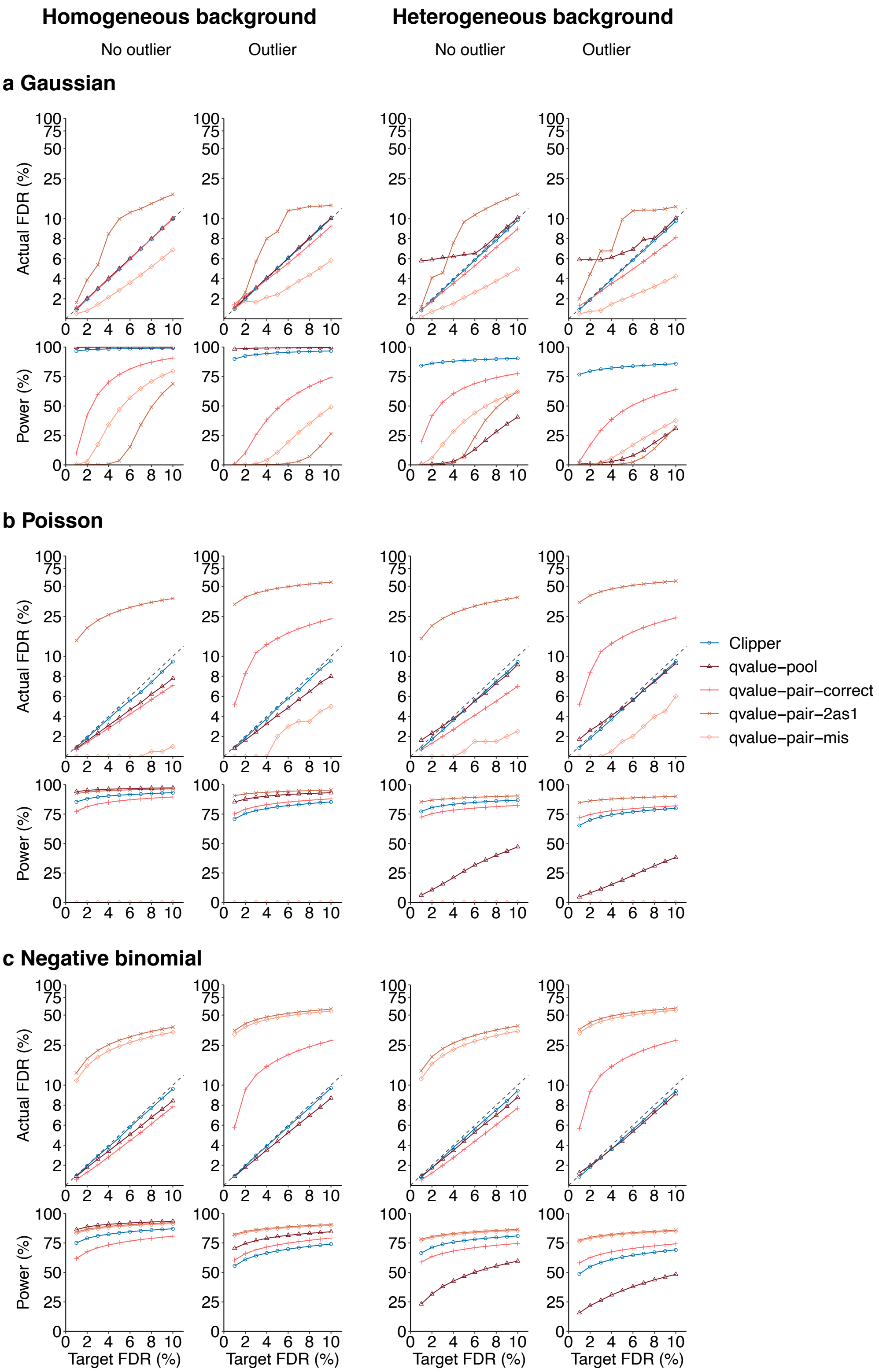


**Figure S7:** In the 3vs3 enrichment analysis without (columns 1 and 3) or with outliers (columns 2 and 4), comparison of Clipper and four other generic FDR control methods using Storey's q-value (qvalue-pooled, qvalue-pair-correct, qvalue-pair-2as1, and qvalue-pair-mis) in terms of their FDR control and power in 3vs3 enrichment analysis with possible outliers. At target FDR thresholds $q \in \{1\%, 2\%, \cdots, 10\%\}$, each method's actual FDRs and power are evaluated on $200$ simulated datasets with $d = 10{,}000$ features generated from (**a**) the Gaussian distribution, (**b**) the Poisson distribution, or (**c**) the negative binomial distribution under homogeneous (two left columns) and heterogeneous (two right columns) background scenarios. Among the methods that control the FDR, Clipper is the second most powerful for homogeneous background and the most powerful for heterogeneous background.

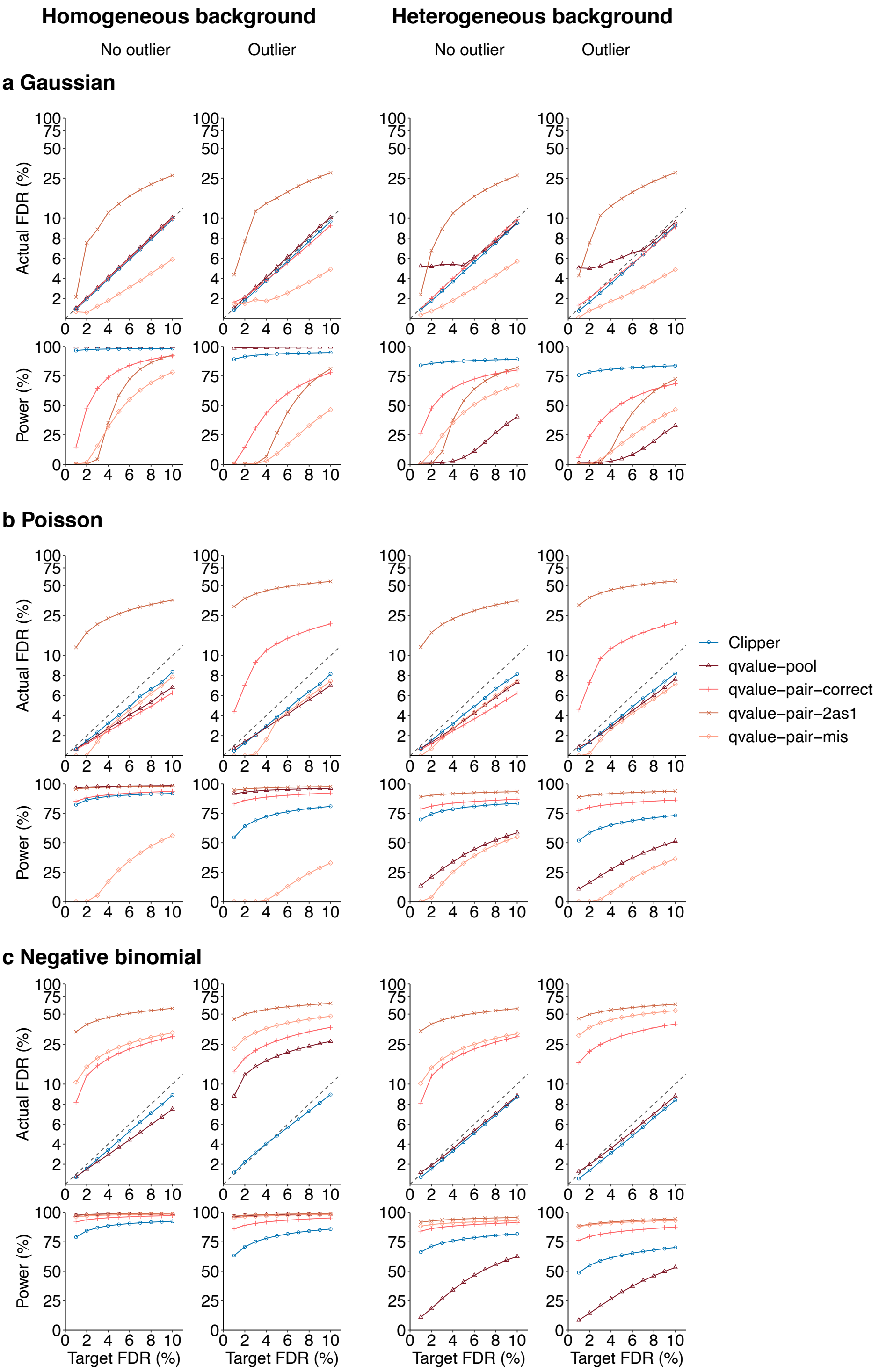


**Figure S8:** In the 3vs3 differential analysis without (columns 1 and 3) or with outliers (columns 2 and 4), comparison of Clipper and four other generic FDR control methods using Storey's q-value (qvalue-pooled, qvalue-pair-correct, qvalue-pair-2as1, and qvalue-pair-mis) in terms of their FDR control and power. At target FDR thresholds $q \in \{1\%, 2\%, \cdots, 10\%\}$, each method's actual FDRs and power are evaluated on $200$ simulated datasets with $d = 10{,}000$ features generated from (**a**) the Gaussian distribution, (**b**) the Poisson distribution, or (**c**) the negative binomial distribution under homogeneous (two left columns) and heterogeneous (two right columns) background scenarios. Among the methods that control the FDR, Clipper is the second most powerful for homogeneous background and the most powerful for heterogeneous background (except for Poisson distribution where Clipper is second to qvalue-pair-correct, an idealistic method).

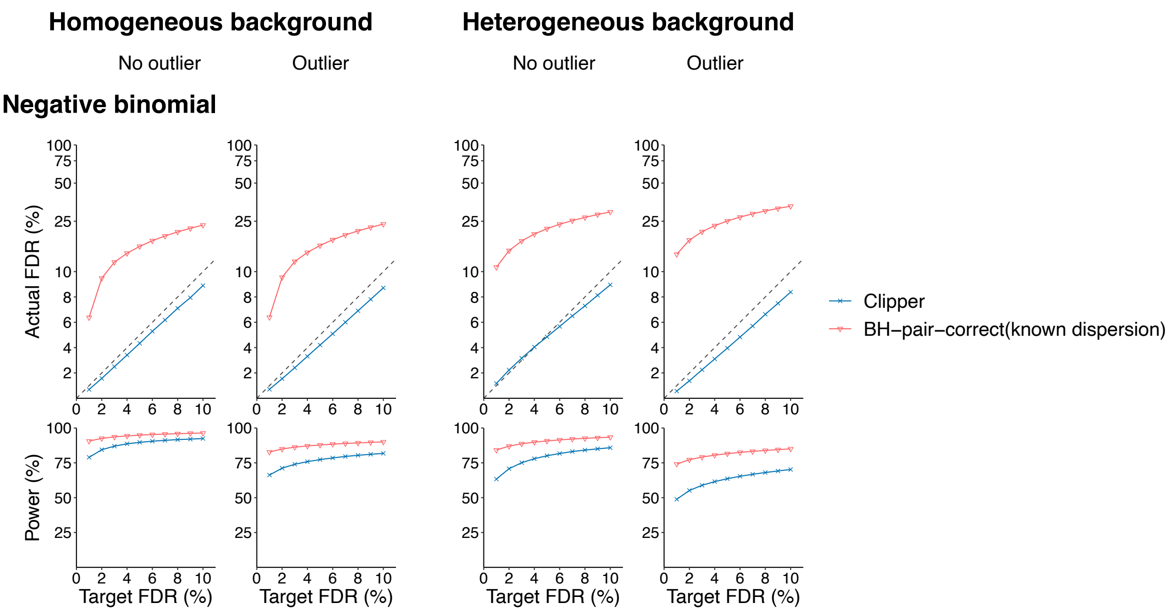


**Figure S9:** In the 3vs3 differential analysis without (columns 1 and 3) or with outliers (columns 2 and 4), comparison of Clipper, BH-pair-correct (known dispersion), and BH-pair-correct (unknown dispersion) in terms of their FDR control and power. At target FDR thresholds $q \in \{1\%, 2\%, \cdots, 10\%\}$, each method's actual FDRs and power are evaluated on $200$ simulated datasets with $d = 10{,}000$ features generated from the negative binomial distribution under homogeneous (two left columns) and heterogeneous (two right columns) background scenarios. BH-pair-correct (unknown dispersion) cannot control the FDR in all settings. In contrast, Clipper is consistently the most powerful for homogeneous and heterogeneous background.

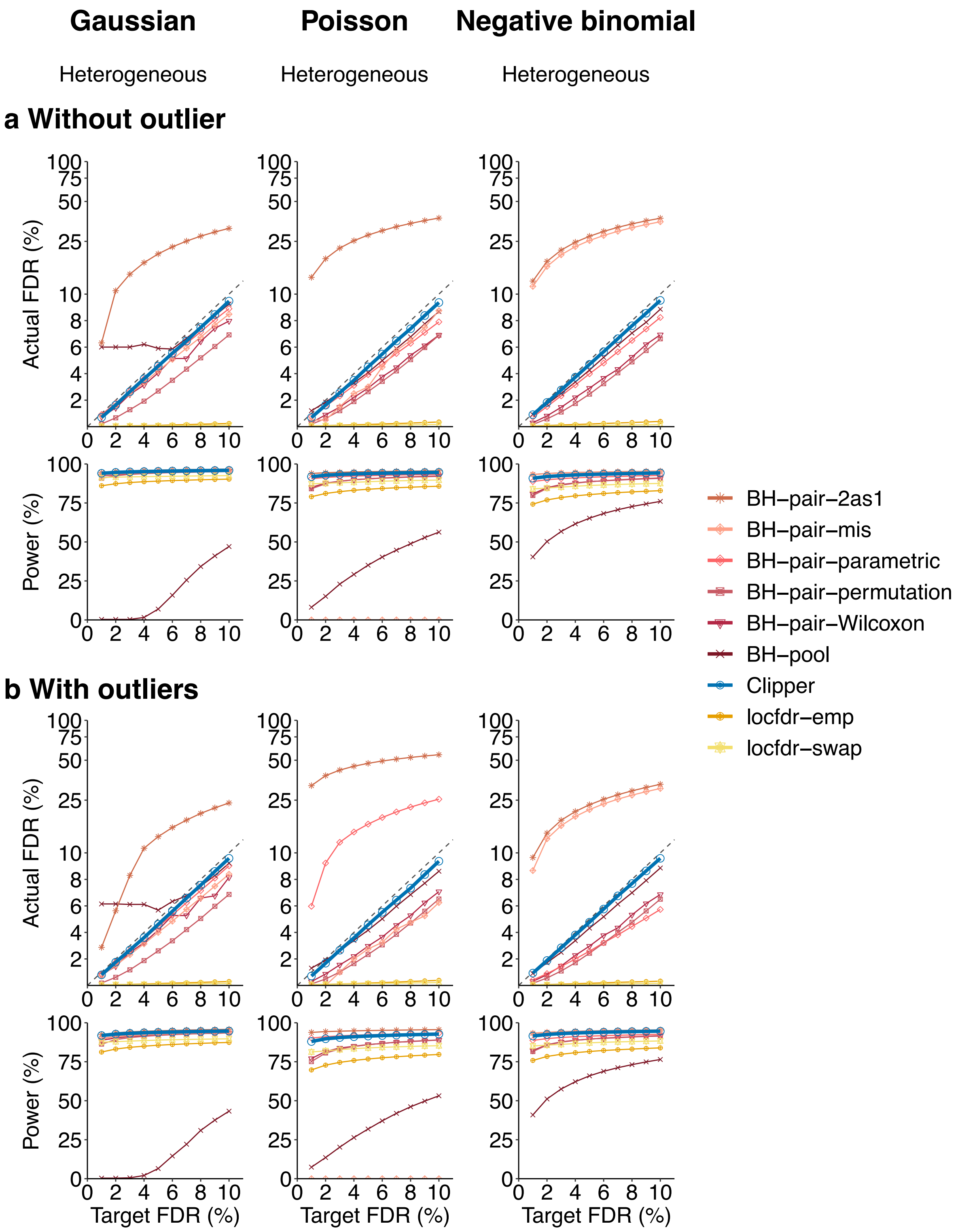


**Figure S10:** In the 10vs10 enrichment analysis with and without outliers, comparison of Clipper and eight generic FDR control methods (BH-pooled, BH-pair-Wilcoxon, BH-pair-parametric, and BH-pair-permutation, BH-pair-2as1, BH-pair-mis, locfdr-emp, and locfdr-swap) in terms of their FDR control and power. At target FDR thresholds $q \in \{1\%, 2\%, \cdots, 10\%\}$, each method's actual FDRs and power are evaluated on $200$ simulated datasets with $d = 10{,}000$ features generated from the Gaussian distribution (left), the Poisson distribution (middle), or the negative binomial distribution (right) under heterogeneous background scenarios. Clipper achieves the highest power for all three distributions.

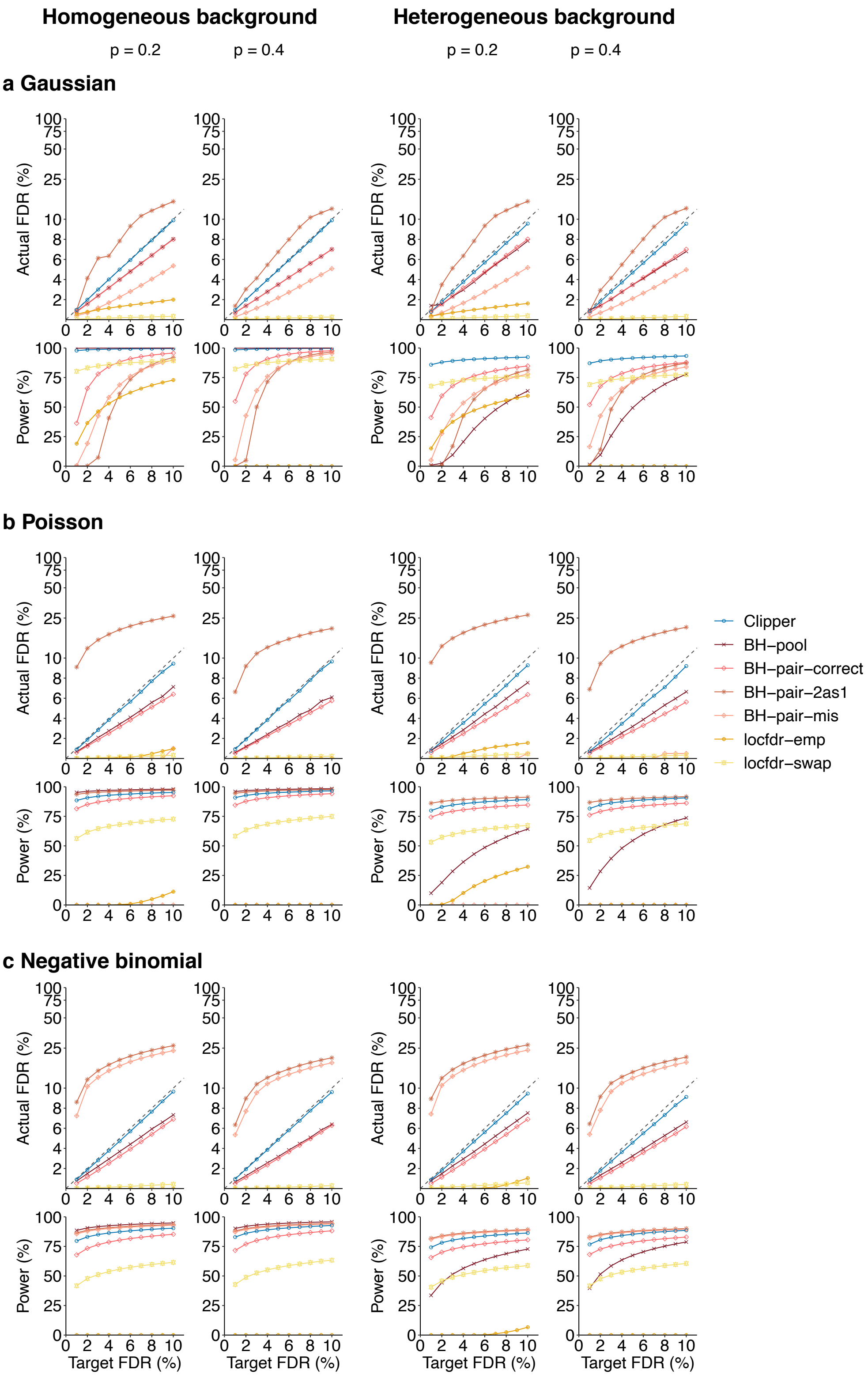


**Figure S11:** In 3vs3 enrichment analysis with different proportions of interesting features without outliers, comparison of Clipper and six generic FDR control methods (BH-pooled, BH-pair-correct, BH-pair-2as1, BH-pair-mis, locfdr-emp, and locfdr-swap) in terms of their FDR control and power. At target FDR thresholds $q \in \{1\%, 2\%, \cdots, 10\%\}$, each method's actual FDRs and power are evaluated on $200$ simulated datasets with $d = 10{,}000$ features generated from the Gaussian distribution, the Poisson distribution, or the negative binomial distribution, with the proportion of interesting features being $0.2$ (columns 1 and 3) or $0.4$ (columns 2 and 4) under homogeneous (columns 1 and 2) and heterogeneous (columns 3 and 4) background scenarios. Clipper achieves the highest power for all distributions.

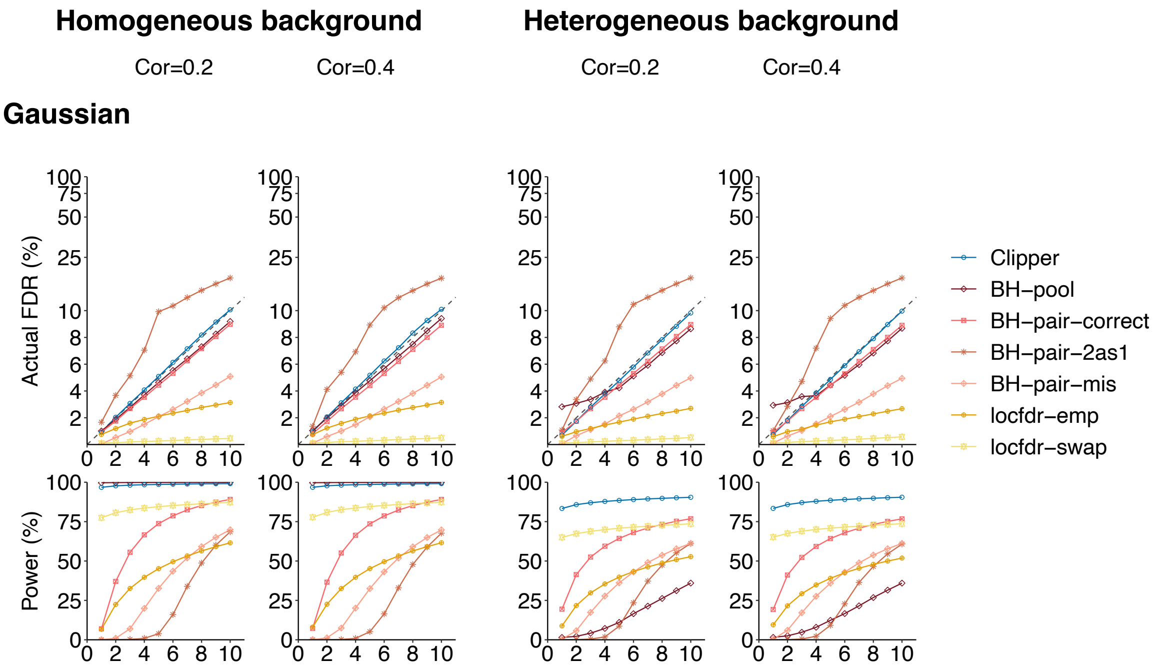


**Figure S12:** In the 3vs3 enrichment analysis with correlated features, comparison of Clipper and six other generic FDR control methods (BH-pooled, BH-pair-correct, BH-pair-2as1, BH-pair-mis, locfdr-emp, and locfdr-swap) in terms of their FDR control and power in 3vs3 enrichment analysis. At target FDR thresholds $q \in \{1\%, 2\%, \cdots, 10\%\}$, each method's actual FDRs and power are evaluated on $200$ simulated datasets with $d = 10{,}000$ features generated from a multivariate Gaussian distribution with a correlation $0.2$ (columns 1 and 3) or $0.4$ (columns 2 and 4) between features. Among the methods that control the FDR, Clipper is the second most powerful for homogeneous background and the most powerful for heterogeneous background.

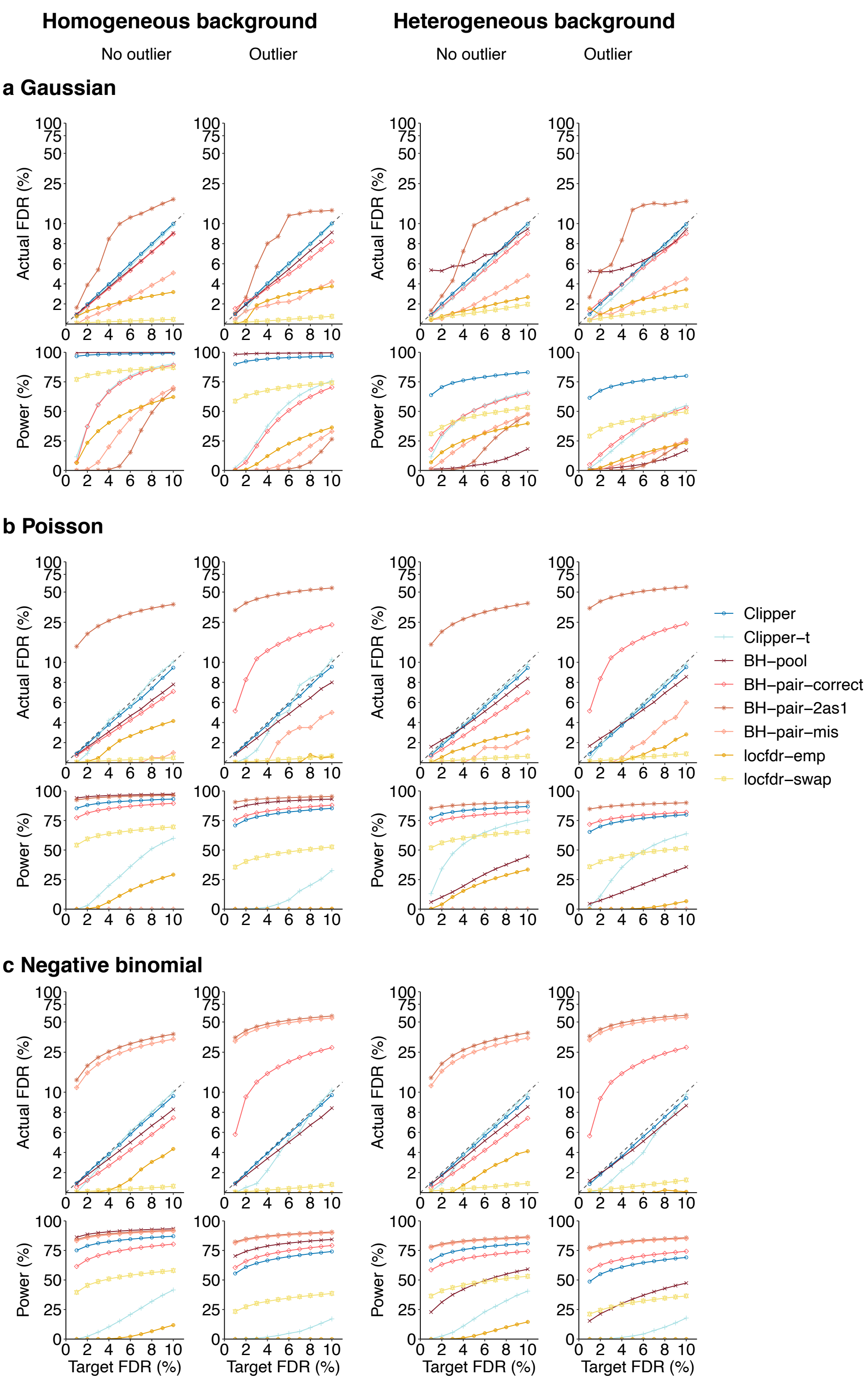


**Figure S13:** In the 3vs3 enrichment analysis with and without outliers, comparison of the default Clipper, the Clipper variant using the $t$ statistic as the contrast score (Clipper-t), and six generic FDR control methods (Clipper BH-pooled, BH-pair-correct, BH-pair-2as1, BH-pair-mis, locfdr-emp, and locfdr-swap) in terms of their FDR control and power. At target FDR thresholds $q \in \{1\%, 2\%, \cdots, 10\%\}$, each method's actual FDRs and power are evaluated on $200$ simulated datasets with $d = 10{,}000$ features generated from the Gaussian distribution, the Poisson distribution, or the negative binomial distribution under homogeneous (columns 1 and 2) and heterogeneous (columns 3 and 4) background scenarios. Clipper achieves higher power than Clipper-t does.

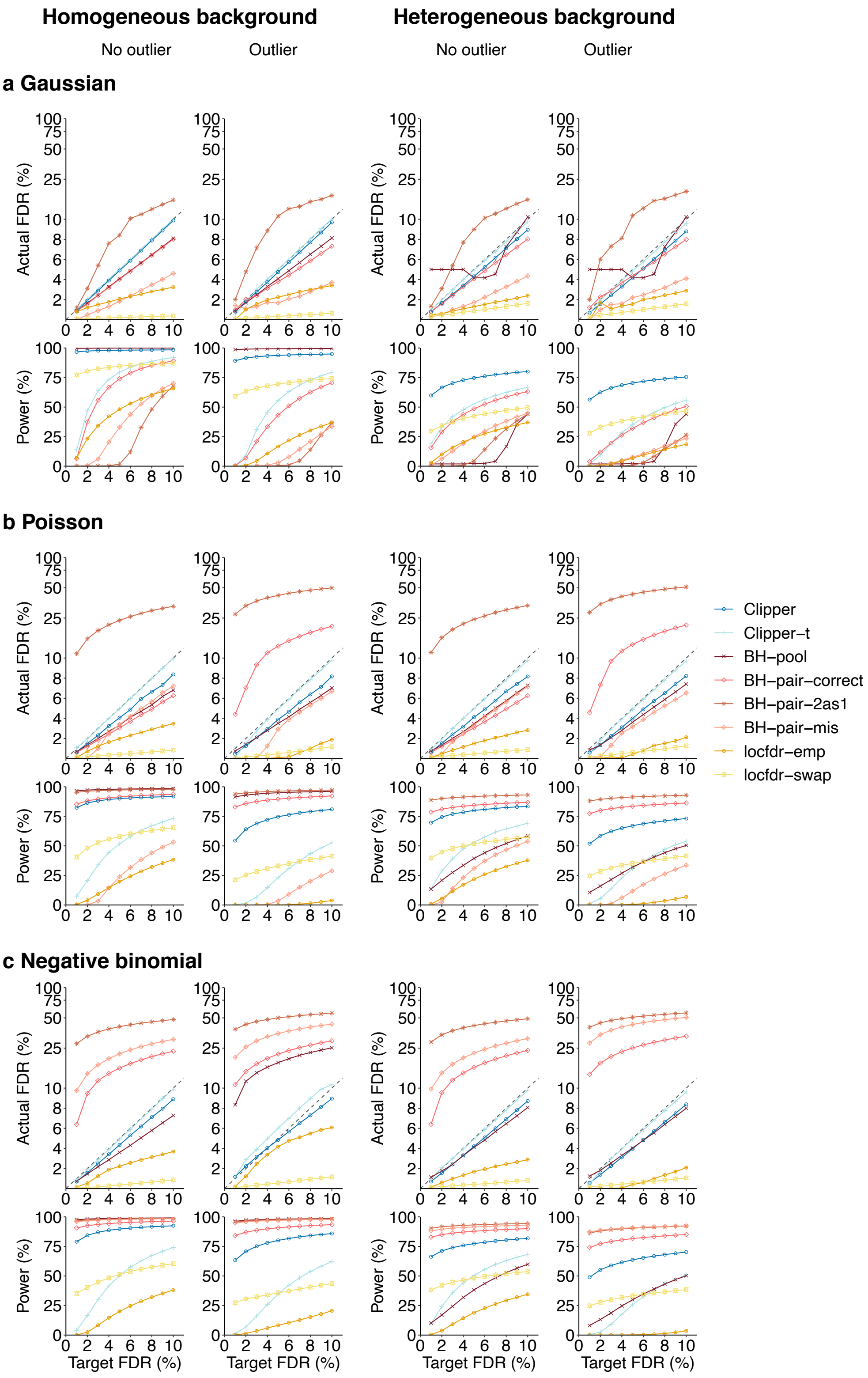


**Figure S14:** In the 3vs3 differential analysis with and without outliers, comparison of the default Clipper, the Clipper variant using the $t$ statistic to calculate the degree of interestingness (Clipper-t), and six generic FDR control methods (Clipper BH-pooled, BH-pair-correct, BH-pair-2as1, BH-pair-mis, locfdr-emp, and locfdr-swap) in terms of their FDR control and power. At target FDR thresholds $q \in \{1\%, 2\%, \cdots, 10\%\}$, each method's actual FDRs and power are evaluated on $200$ simulated datasets with $d = 10{,}000$ features generated from the Gaussian distribution, the Poisson distribution, or the negative binomial distribution under homogeneous (columns 1 and 2) and heterogeneous (columns 3 and 4) background scenarios. Clipper achieves higher power than Clipper-t does.

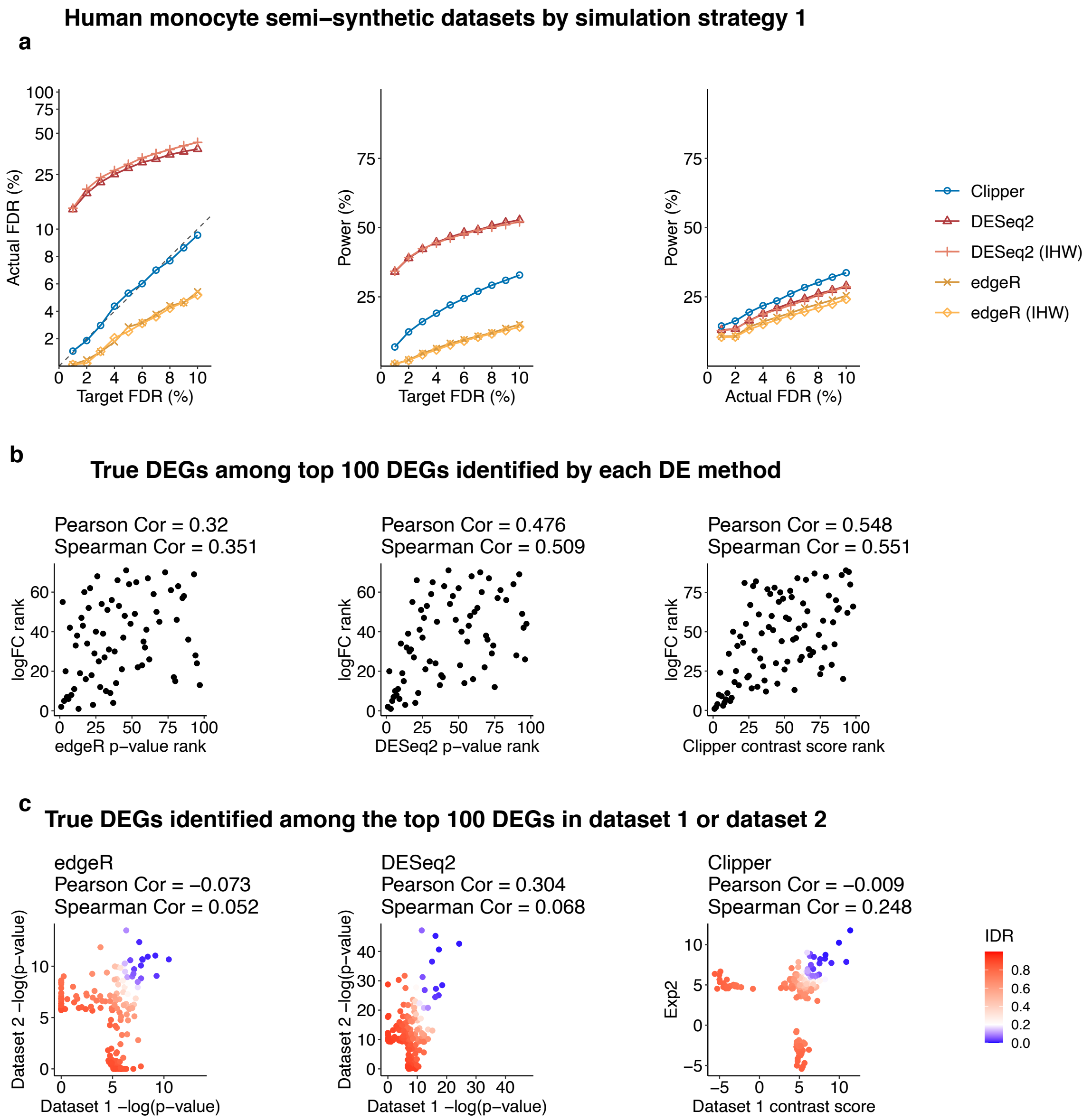


**Figure S15:** Comparison of Clipper and two popular DEG identification methods—edgeR and DESeq2—in DEG analysis on semi-synthetic bulk RNA-seq data (generated from human monocyte real data using simulation strategy 1 in Supp. Section S6.3). **(a)** FDR control, power given the same target FDR, and power given the same actual FDR. **(b)** Ranking consistency of the true DEGs among the top $100$ DEGs identified by each method. The consistency is defined between the genes' ranking based on edgeR/DESeq2's p-values or Clipper's contrast scores and their ranking based on true expression fold changes. **(c)** Reproducibility between two semi-synthetic datasets as technical replicates. Three reproducibility criteria are used: the IDR, Pearson correlation, and Spearman correalation. Each criterion is calculated for edgeR/DESeq2's p-values or Clipper's contrast scores on the two semi-synthetic datasets. Among the three methods, only Clipper controls the FDR, and Clipper achieves the highest power, the best gene ranking consistency, and the best reproducibility.

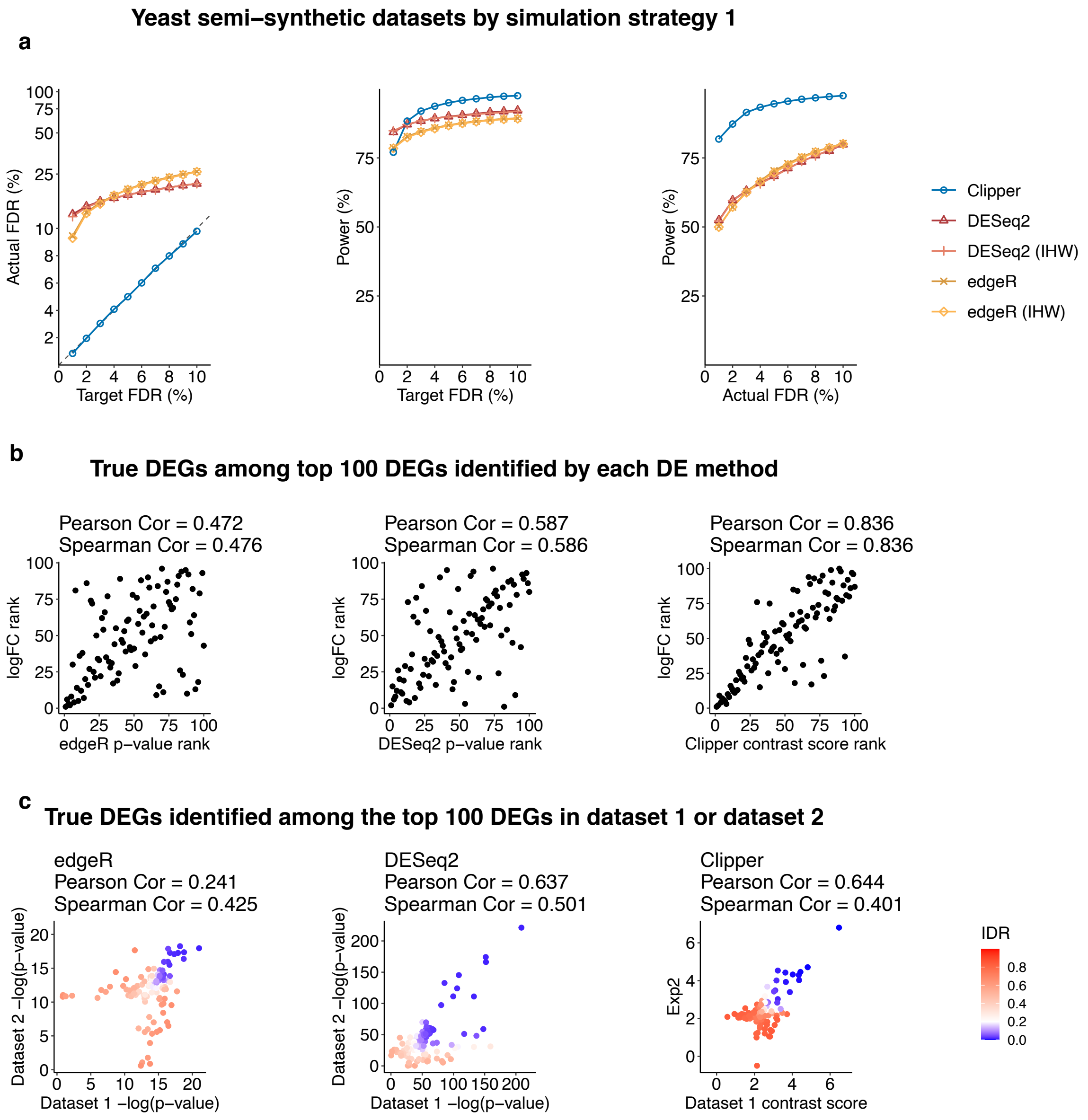


**Figure S16:** Comparison of Clipper and two popular DEG identification methods—edgeR and DESeq2—in DEG analysis on semi-synthetic bulk RNA-seq data (generated from yeast real data using simulation strategy 1 in Supp. Section S6.3). **(a)** FDR control, power given the same target FDR, and power given the same actual FDR. **(b)** Ranking consistency of the true DEGs among the top $100$ DEGs identified by each method. The consistency is defined between the genes' ranking based on edgeR/DESeq2's p-values or Clipper's contrast scores and their ranking based on true expression fold changes. **(c)** Reproducibility between two semi-synthetic datasets as technical replicates. Three reproducibility criteria are used: the IDR, Pearson correlation, and Spearman correalation. Each criterion is calculated for edgeR/DESeq2's p-values or Clipper's contrast scores on the two semi-synthetic datasets. Among the three methods, only Clipper controls the FDR, and Clipper achieves the highest power, the best gene ranking consistency, and the best reproducibility.

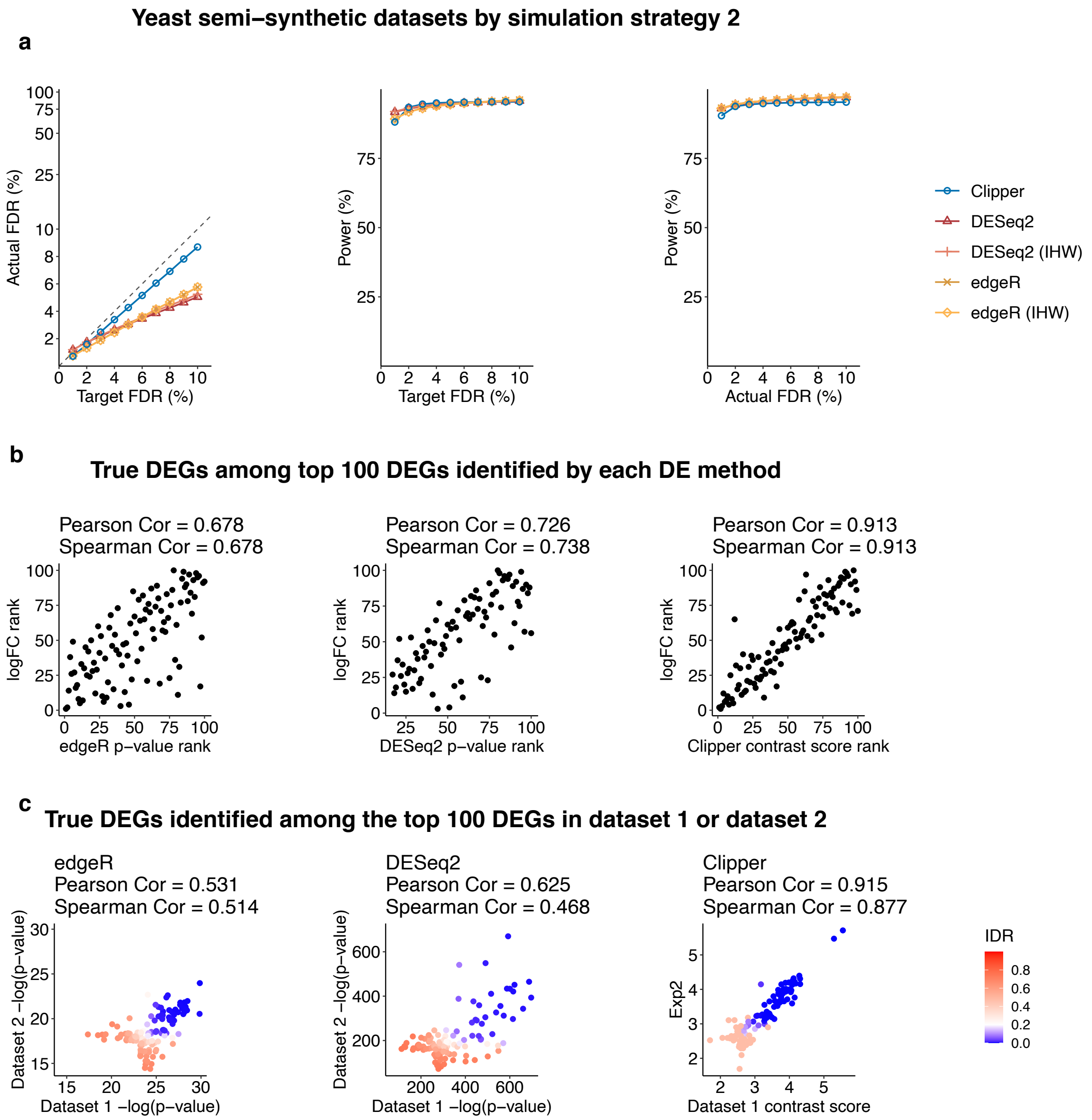


**Figure S17:** Comparison of Clipper and two popular DEG identification methods—edgeR and DESeq2—in DEG analysis on semi-synthetic bulk RNA-seq data (generated from yeast real data using simulation strategy 2 in Supp. Section S6.3). **(a)** FDR control, power given the same target FDR, and power given the same actual FDR. **(b)** Ranking consistency of the true DEGs among the top $100$ DEGs identified by each method. The consistency is defined between the genes' ranking based on edgeR/DESeq2's p-values or Clipper's contrast scores and their ranking based on true expression fold changes. **(c)** Reproducibility between two semi-synthetic datasets as technical replicates. Three reproducibility criteria are used: the IDR, Pearson correlation, and Spearman correalation. Each criterion is calculated for edgeR/DESeq2's p-values or Clipper's contrast scores on the two semi-synthetic datasets. Among the three methods, only Clipper controls the FDR, and Clipper achieves the highest power, the best gene ranking consistency, and the best reproducibility.

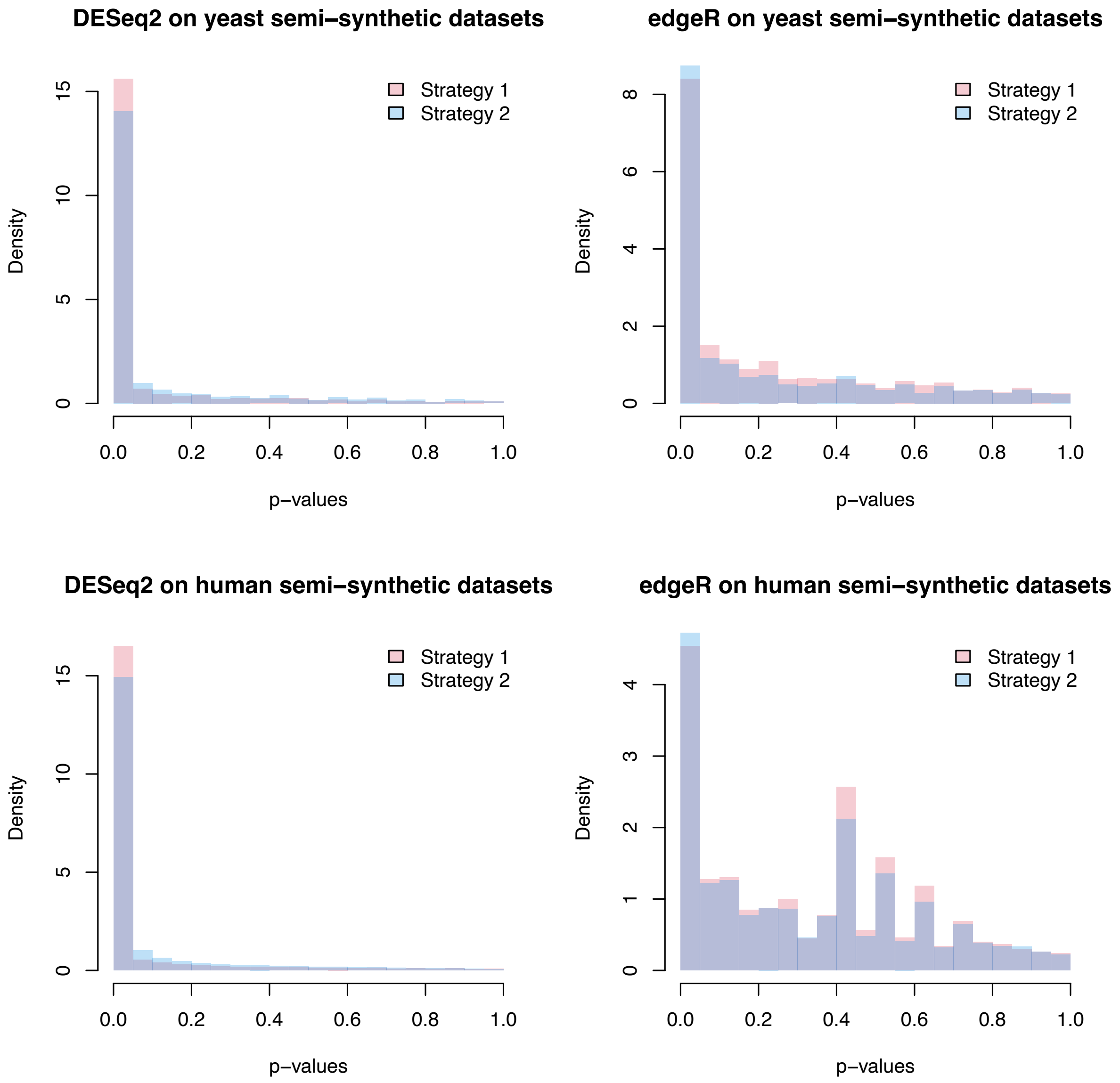


**Figure S18:** Histograms of p-values (one p-value per non-DEG) that are obtained by testing whether each non-DEG's p-values (output by DESeq2 or edgeR) follow a uniform distribution. For each real dataset (yeasts on the top and human monocytes on the bottom) and each simulation strategy (red for strategy 1 and blue for strategy 2), a histogram is plotted for DESeq2 (left) or edgeR (right); each p-value is calculated across $100$ semi-synthetic datasets (excluding NA p-values). In each panel, more right skewed histograms are considered better.

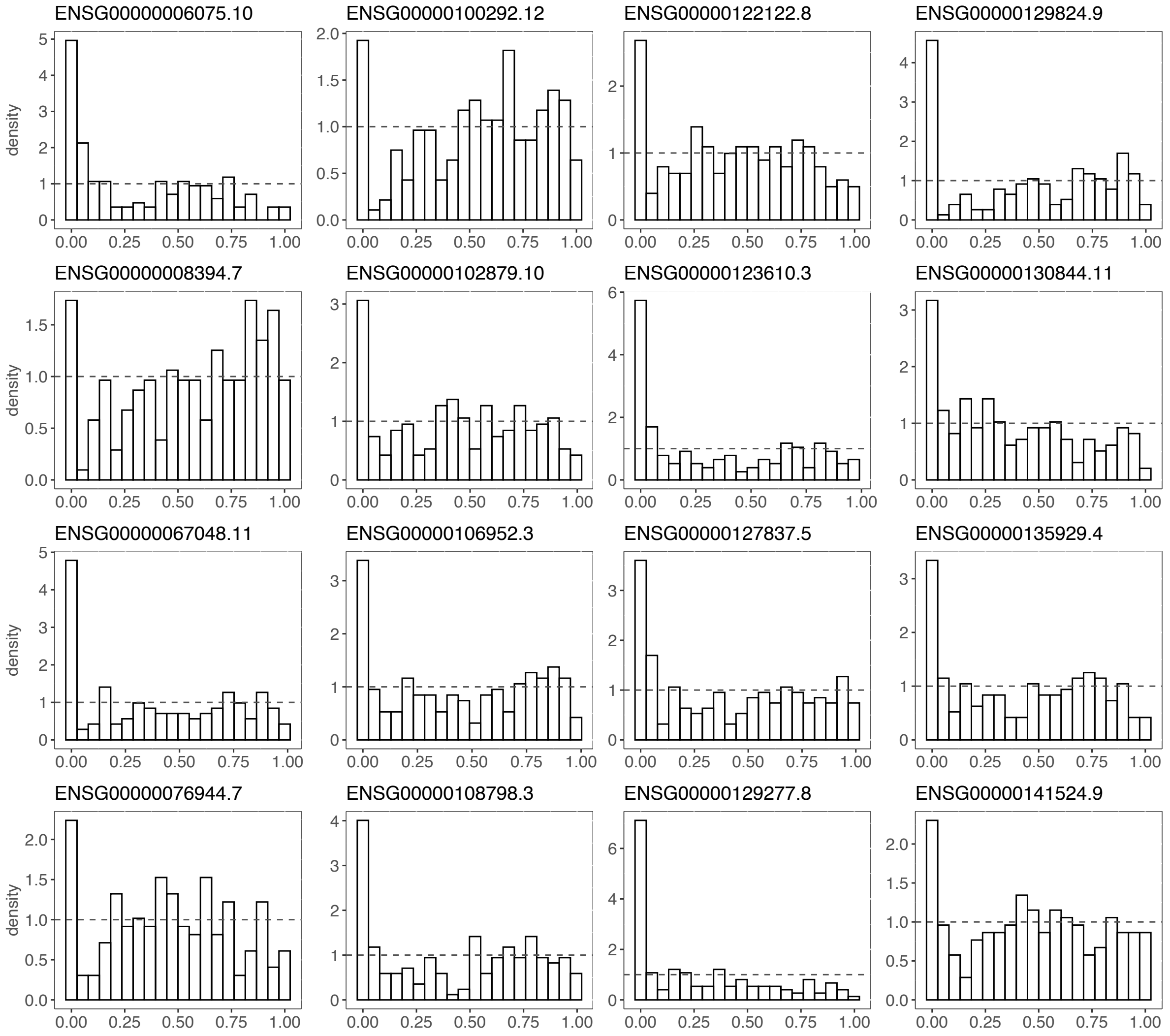


**Figure S19:** The p-value distributions of $16$ non-DEGs that are most frequently identified by DESeq2 at $q = 5\%$ from $200$ semi-synthetic datasets. The p-values of these $16$ genes tend to be overly small, and their distributions are non-uniform with a mode close to $0$.

a

GO terms enriched in Clipper−specific DEGs in Clipper vs. DESeq2 comparison

| GO term (ID) | qvalue (Clipper) |
|---|---|
| neutrophil activation (GO:0042119) | 3.104557e−10 |
| granulocyte activation (GO:0036230) | 3.104557e−10 |
| neutrophil degranulation (GO:0043312) | 8.587750e−10 |
| neutrophil activation involved in immune response (GO:0002283) | 8.591455e−10 |
| neutrophil mediated immunity (GO:0002446) | 3.104557e−10 |

b

GO terms enriched in Clipper−specific DEGs in Clipper vs. edgeR comparison

| GO term (ID) | qvalue (Clipper) |
|---|---|
| neutrophil degranulation (GO:0043312) | 8.587750e−10 |
| neutrophil activation involved in immune response (GO:0002283) | 8.591455e−10 |
| neutrophil activation (GO:0042119) | 3.104557e−10 |
| neutrophil mediated immunity (GO:0002446) | 3.104557e−10 |
| granulocyte activation (GO:0036230) | 3.104557e−10 |
| cellular response to chemical stress (GO:0062197) | 2.157116e−03 |
| response to oxidative stress (GO:0006979) | 3.141033e−03 |
| cellular response to oxidative stress (GO:0034599) | 2.902893e−03 |

**Figure S20:** Enrichment q-values of GO terms that are found enriched in the DEGs that are uniquely identified by Clipper in pairwise comparison of **(a)** Clipper vs. edgeR and **(b)** Clipper vs. DESeq2. These GO terms are all related to immune response and thus biologically meaningful.

a

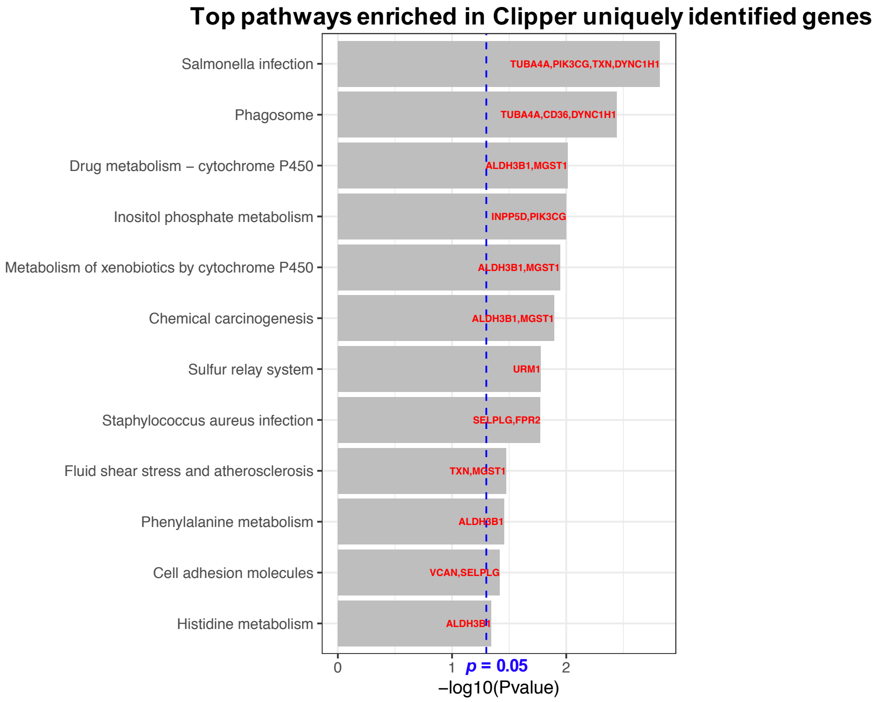


b

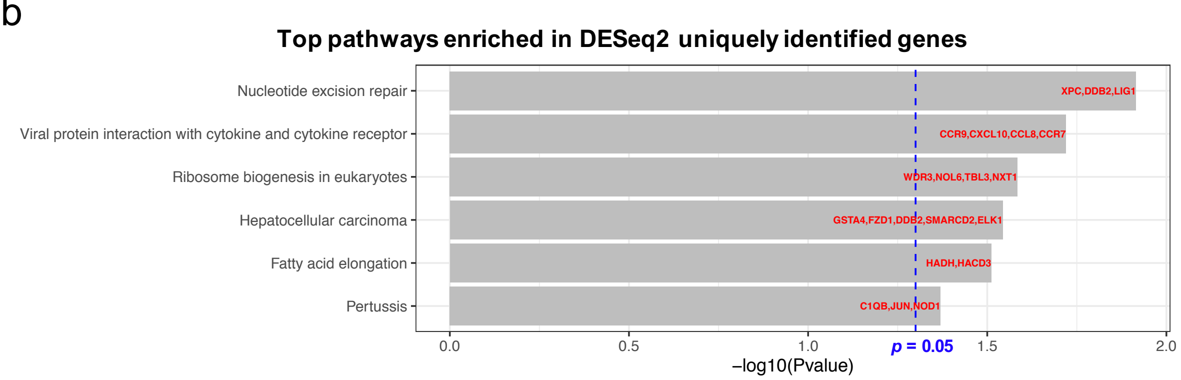


**Figure S21:** The p-values of the top enriched pathways in the DEGs that are uniquely identified by **(a)** Clipper and **(b)** DESeq2; i.e., the DEGs that are only identified by one method by missed by the other two methods. There are more immune-related pathways enriched in (a) than (b).

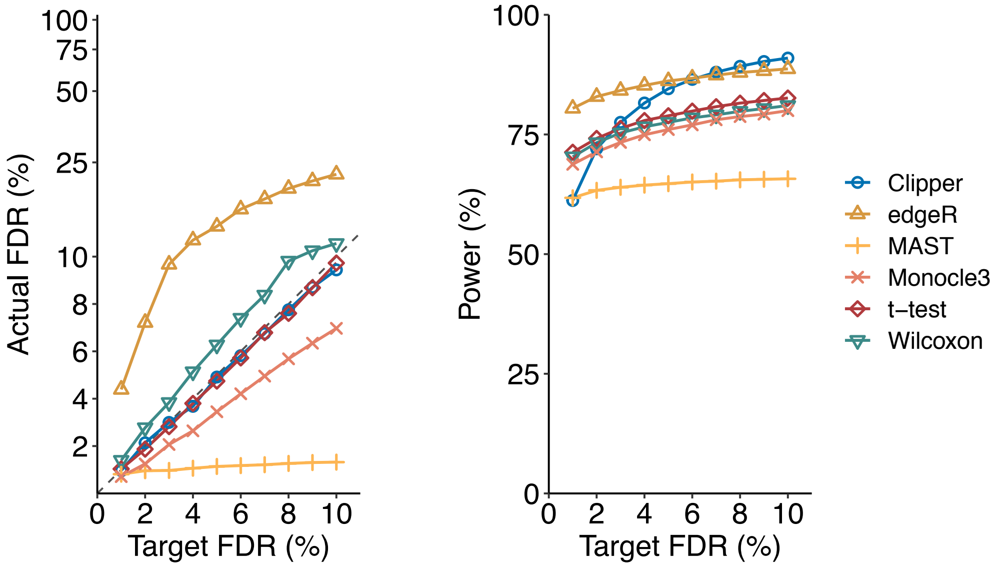


**Figure S22:** Comparison of Clipper and five scRNA-seq DEG identification methods on semi-synthetic Drop-seq data generated by scDesign2 (based on a real Drop-seq dataset of PBMCs). The target FDR threhold $q$ ranges from $1\%$ to $10\%$. In the "Actual FDR vs. Target FDR" plot (left), points above the dashed diagonal line indicate failed FDR control. Clipper controls the FDR while maintaining high power, demonstrating Clipper's good performance in single-cell DE analyses.

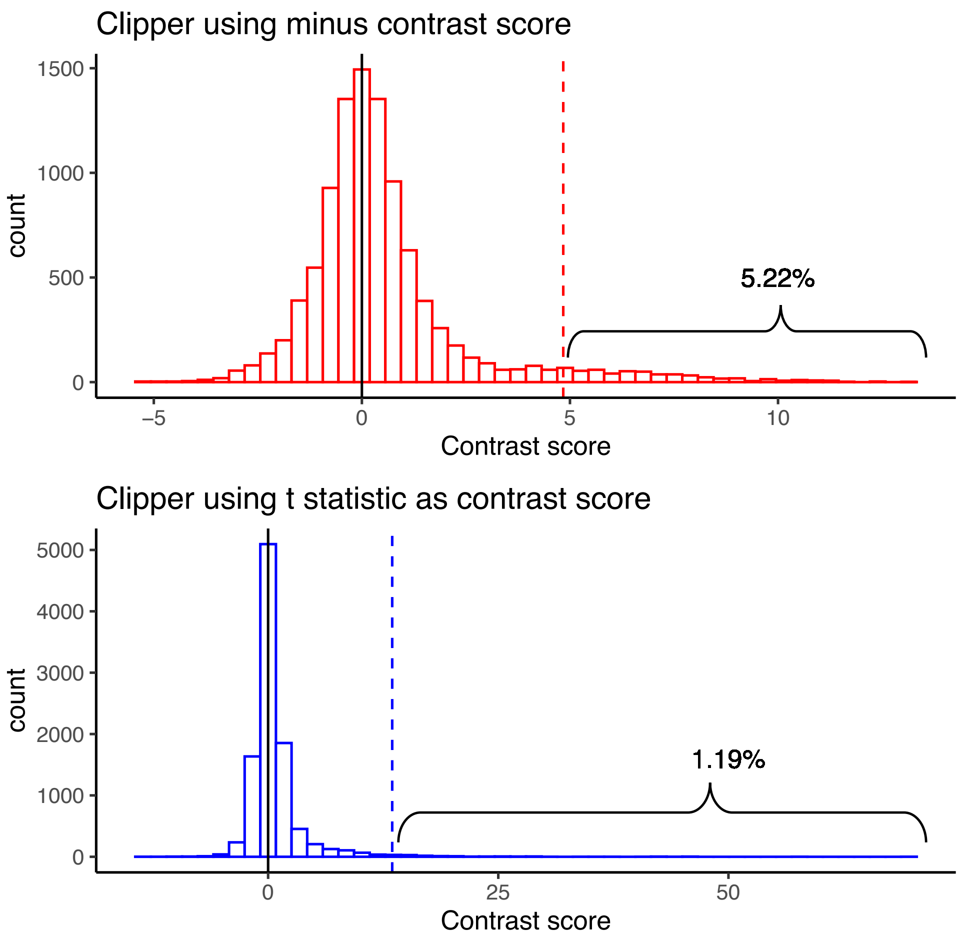


**Figure S23:** In the 3vs3 enrichment analysis, distributions of contrast scores used by two Clipper variants: the default Clipper using the minus contrast score (top) and the Clipper variant using the two-sample $t$ statistic (bottom). Features are generated from the Gaussian distribution under the heterogeneous background scenario (see Supp. Section S4). The vertical dashed lines indicate the contrast score cutoffs found by the BC procedure at the target FDR threshold $q = 1\%$. The distribution of the minus contrast scores has a heavier right tail ($5.22\%$) than that of the distribution of the $t$ statistic contrast scores ($1.19\%$).

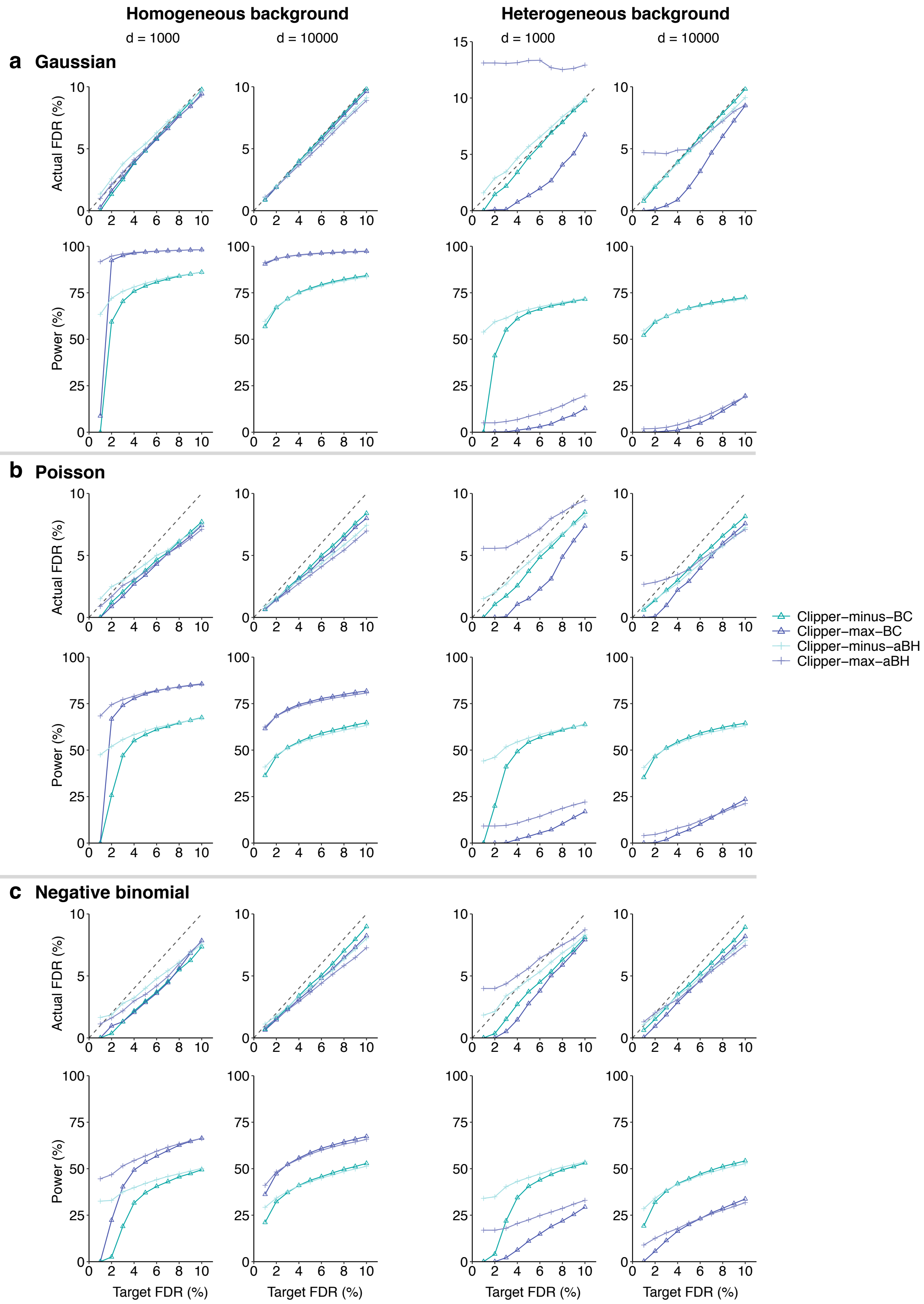


**Figure S24:** In 1vs1 enrichment analysis, comparison of four Clipper variant algorithms (Clipper-minus-BC, Clipper-minus-aBH, Clipper-max-BC, and Clipper-max-aBH) in terms of their FDR control and power. At target FDR thresholds $q \in \{1\%, 2\%, \cdots, 10\%\}$, each method's actual FDRs and power are evaluated on $200$ simulated datasets with $d = 1000$ or $10{,}000$ features generated from (**a**) the Gaussian distribution, (**b**) the Poisson distribution, or (**c**) the negative binomial distribution under homogeneous (two left columns) and heterogeneous (two right columns) background scenarios. Clipper-minus-BC is chosen as the default implementation under this scenario.

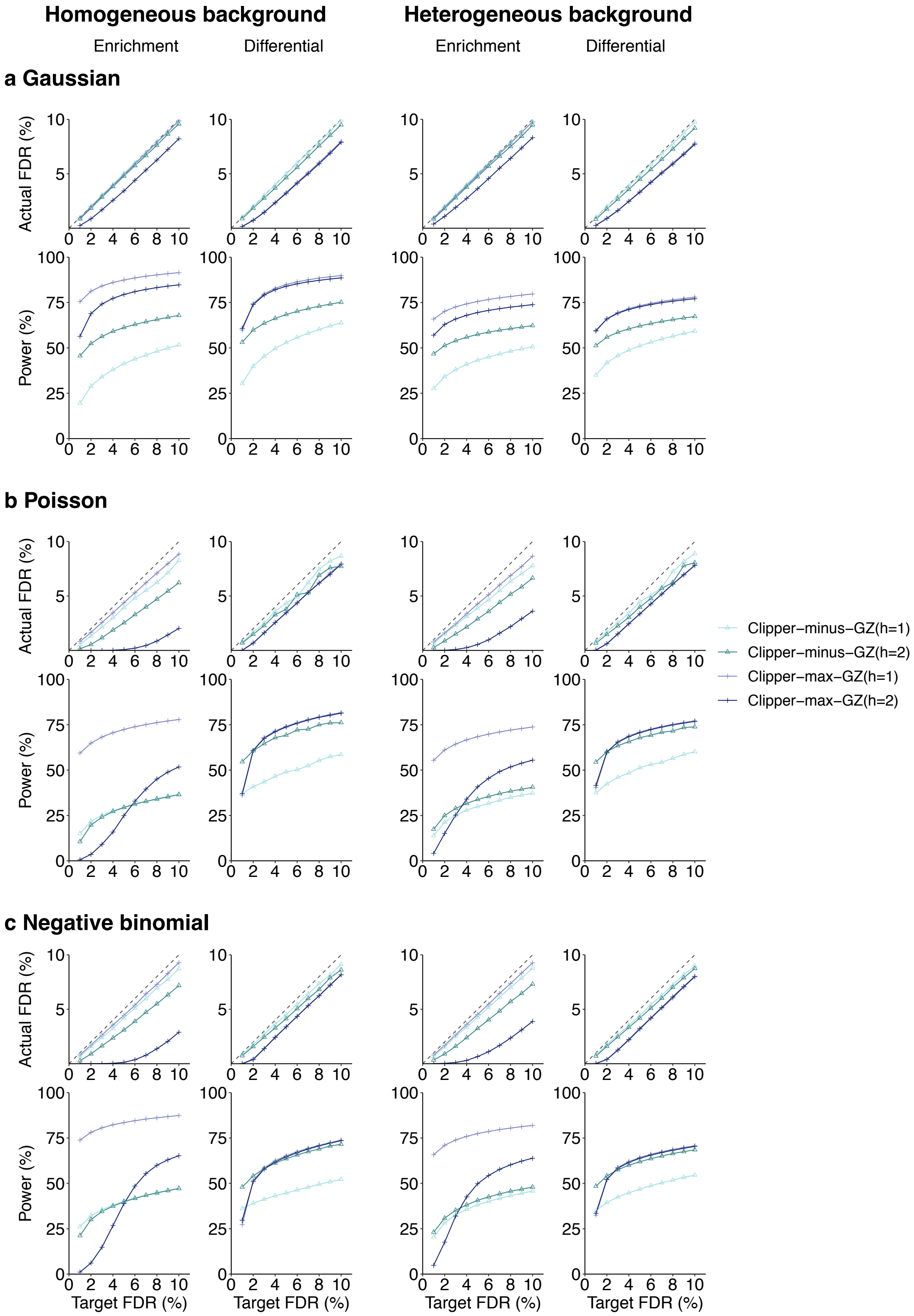


**Figure S25:** In the 2vs1 enrichment analysis (columns 1 and 3) and differential analysis (columns 2 and 4), comparison of four Clipper variant algorithms (Clipper-minus-GZ(h=1), Clipper-minus-GZ(h=2), Clipper-max-GZ(h=1), and Clipper-max-GZ(h=2)) in terms of their FDR control and power. At target FDR thresholds $q \in \{1\%, 2\%, \cdots, 10\%\}$, each method's actual FDRs and power are evaluated on $200$ simulated datasets with $d = 10{,}000$ features generated from (**a**) the Gaussian distribution, (**b**) the Poisson distribution, or (**c**) the negative binomial distribution under homogeneous (two left columns) and heterogeneous (two right columns) background scenarios. Clipper-max-GZ(h=1) is chosen as the default implementation under this scenario.

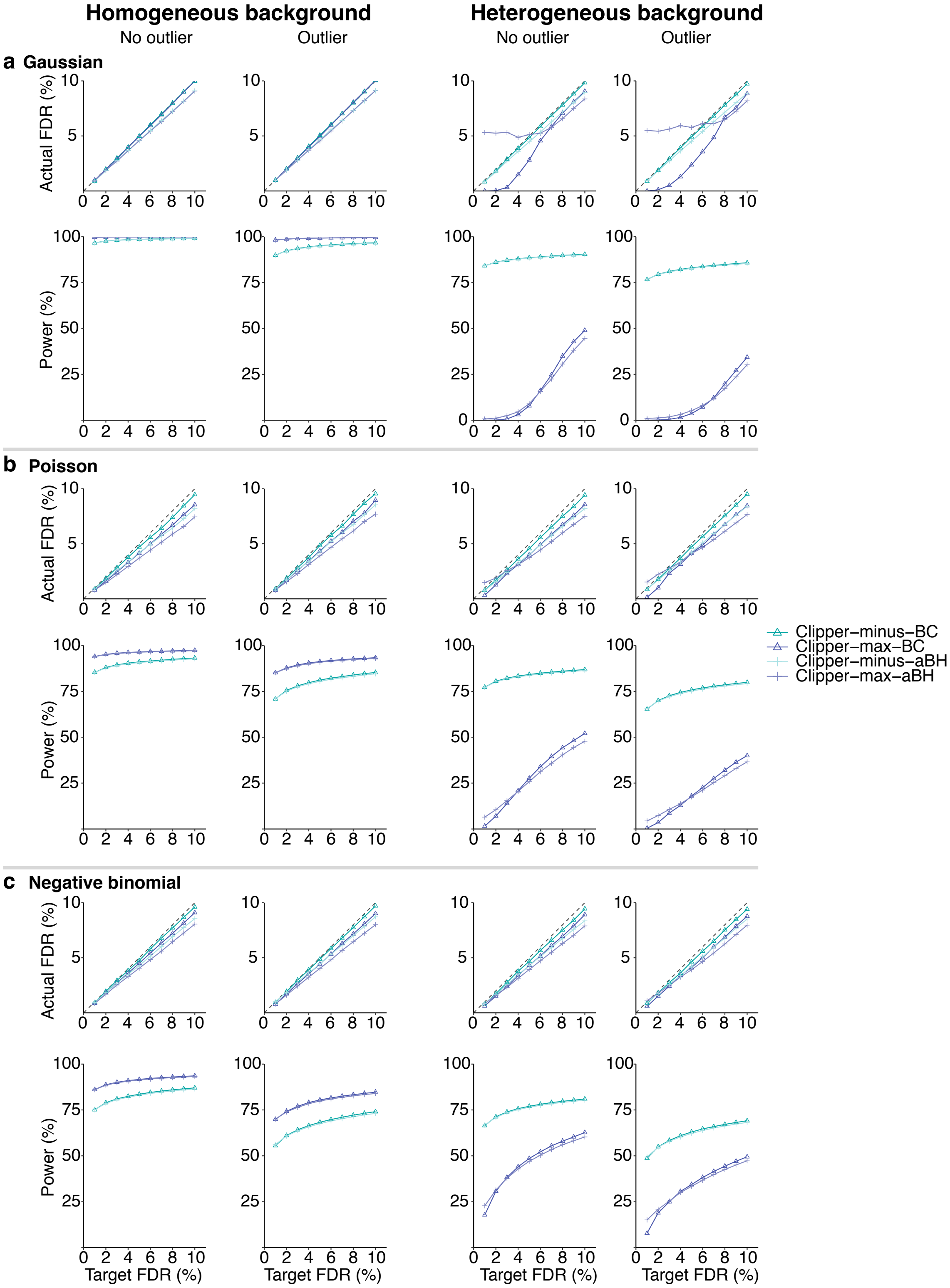


**Figure S26:** In 3vs3 enrichment analysis without (columns 1 and 3) or with outliers (columns 2 and 4), comparison of four Clipper variant algorithms (Clipper-minus-BC, Clipper-minus-aBH, Clipper-max-BC, and Clipper-max-aBH) in terms of their FDR control and power. At target FDR thresholds $q \in \{1\%, 2\%, \cdots, 10\%\}$, each method's actual FDRs and power are evaluated on $200$ simulated datasets with $d = 10{,}000$ features generated from (**a**) the Gaussian distribution, (**b**) the Poisson distribution, or (**c**) the negative binomial distribution under homogeneous (two left columns) and heterogeneous (two right columns) background scenarios. Clipper-minus-BC is chosen as the default implementation under this scenario.

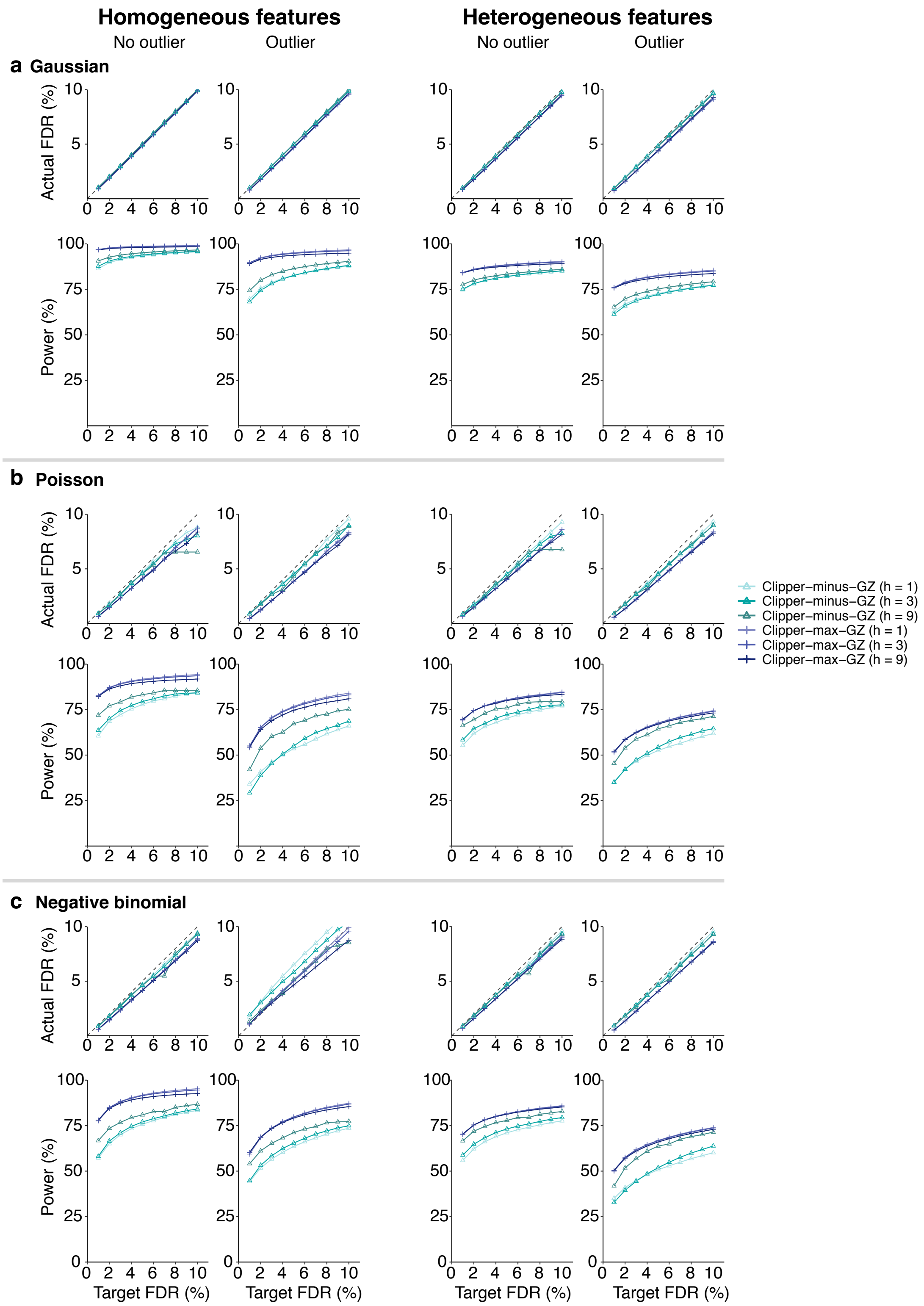


**Figure S27:** In 3vs3 differential analysis without (columns 1 and 3) or with outliers (columns 2 and 4), comparison of six Clipper variant algorithms (Clipper-minus-GZ(h=1), Clipper-minus-GZ(h=3), Clipper-minus-GZ(h=9), Clipper-max-GZ(h=1), Clipper-max-GZ(h=3), and Clipper-max-GZ(h=9)) in terms of their FDR control and power. At target FDR thresholds $q \in \{1\%, 2\%, \cdots, 10\%\}$, each method's actual FDRs and power are evaluated on $200$ simulated datasets with $d = 10{,}000$ features generated from (**a**) the Gaussian distribution, (**b**) the Poisson distribution, or (**c**) the negative binomial distribution under homogeneous (two left columns) and heterogeneous (two right columns) background scenarios. Clipper-max-GZ(h=1) is chosen as the default implementation under this scenario.

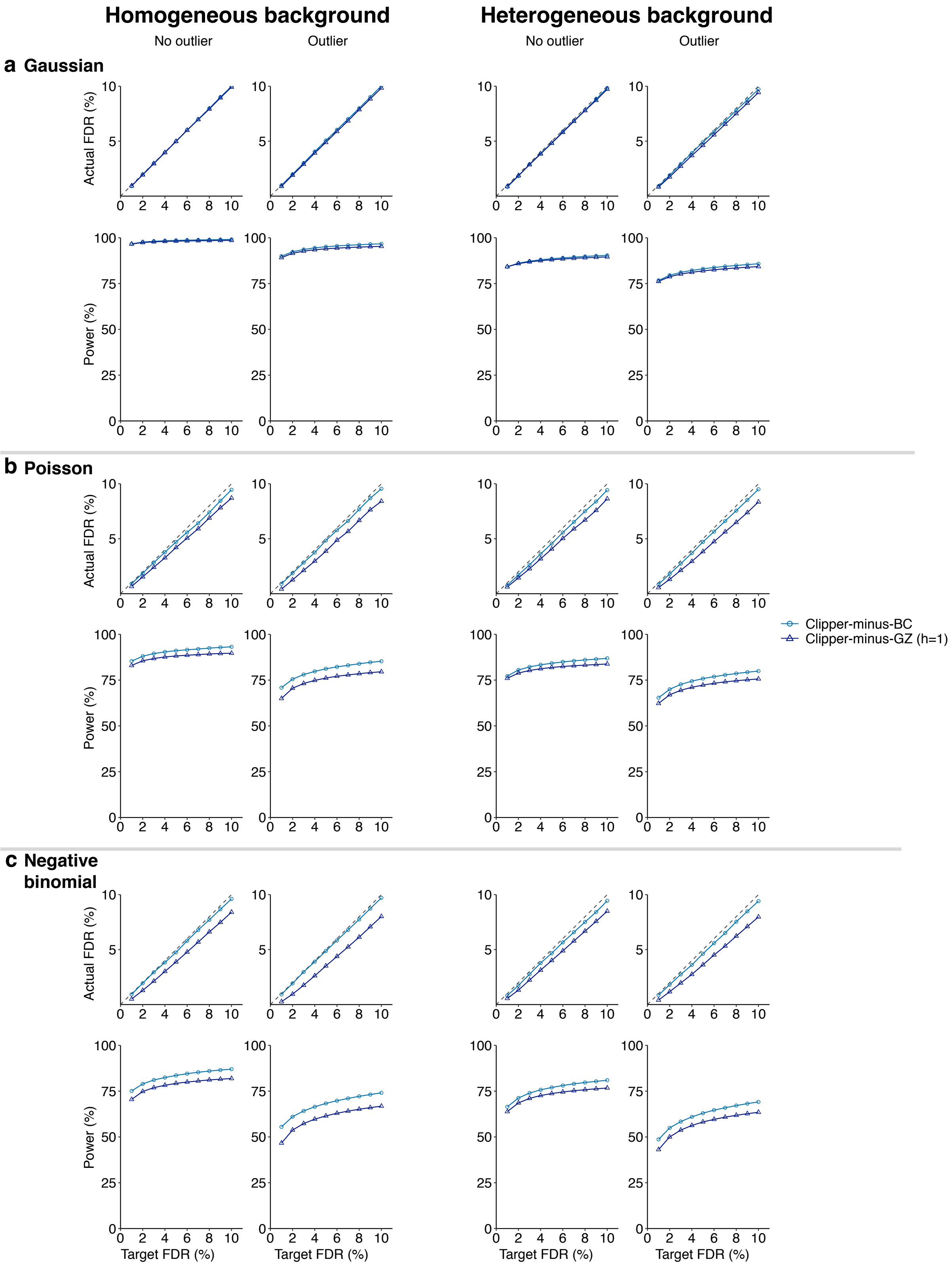


**Figure S28:** In the 3vs3 enrichment analysis without (columns 1 and 3) or with outliers (columns 2 and 4), comparison of two Clipper variant algorithms (Clipper-minus-BC, Clipper-max-GZ(h=1)) in terms of their FDR control and power. At target FDR thresholds $q \in \{1\%, 2\%, \cdots, 10\%\}$, each method's actual FDRs and power are evaluated on $200$ simulated datasets with $d = 10{,}000$ features generated from (**a**) the Gaussian distribution, (**b**) the Poisson distribution, or (**c**) the negative binomial distribution under homogeneous (two left columns) and heterogeneous (two right columns) background scenarios. Clipper-minus-BC is chosen as the default implementation under this scenario.

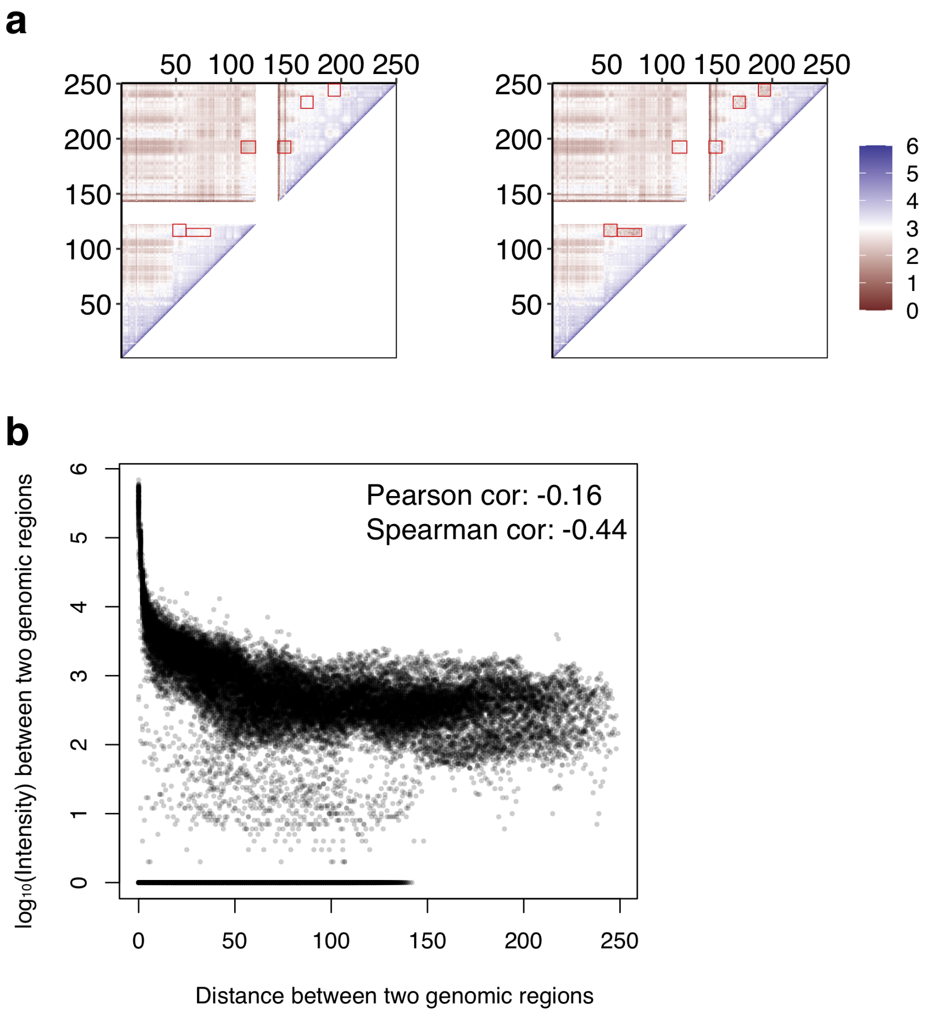


**Figure S29:** **(a)** $\log_{10}$-transformed mean Hi-C interaction matrices ($\mu_X$ and $\mu_Y$ in Section S6.5) under the two conditions. DIR regions are highlighted in red squares. **(b)** In one randomly picked Hi-C semi-synthetic dataset, closer genomic regions tend to have higher contact intensities.
